# Distinct patterns of genetic variation at low-recombining genomic regions represent haplotype structure

**DOI:** 10.1101/2021.12.22.473882

**Authors:** Jun Ishigohoka, Karen Bascón-Cardozo, Andrea Bours, Janina Fuß, Arang Rhie, Jacquelyn Mountcastle, Bettina Haase, William Chow, Joanna Collins, Kerstin Howe, Marcela Uliano-Silva, Olivier Fedrigo, Erich D. Jarvis, Javier Pérez-Tris, Juan Carlos Illera, Miriam Liedvogel

**Affiliations:** Max Planck Institute for Evolutionary Biology, Plön, Germany; Institute of Clinical Molecular Biology (IKMB), Kiel University, Kiel, Germany; Genome Informatics Section, Computational and Statistical Genomics Branch, National Human Genome Research Institute, National Institutes of Health, Bethesda, MD, USA; The Vertebrate Genome Lab, Rockefeller University, New York, NY, USA; Wellcome Sanger Institute, Cambridge, UK; Laboratory of Neurogenetics of Language, Rockefeller University, New York, NY, USA; The Howards Hughes Medical Institute, Chevy Chase, MD, USA; Department of Biodiversity, Ecology and Evolution, Complutense University of Madrid, Madrid, Spain; Biodiversity Research Institute (CSIC-Oviedo University-Principality of Asturias), Oviedo University, Mieres, Spain; Institute of Avian Research, Wilhelmshaven, Germany

**Author notes:** Correspondence: Jun Ishigohoka < >, Miriam Liedvogel < >.

## Abstract

Genetic variation of the entire genome represents population structure, yet individual loci can show distinct patterns. Such deviations identified through genome scans have often been attributed to effects of selection instead of randomness. This interpretation assumes that long enough genomic intervals average out randomness in underlying genealogies, which represent local genetic ancestries. However, an alternative explanation to distinct patterns has not been fully addressed: too few genealogies to average out the effect of randomness. Specifically, distinct patterns of genetic variation may be due to reduced local recombination rate, which reduces the number of genealogies in a genomic window. Here, we associate distinct patterns of local genetic variation with reduced recombination rates in a songbird, the Eurasian blackcap (*Sylvia atricapilla*), using genome sequences and recombination maps. We find that distinct patterns of local genetic variation reflect haplotype structure at low-recombining regions either shared in most populations or found only in a few populations. At the former species-wide low-recombining regions, genetic variation depicts conspicuous haplotypes segregating in multiple populations. At the latter population-specific low-recombining regions, genetic variation represents variance among cryptic haplotypes within the low-recombining populations. With simulations, we confirm that these distinct patterns of haplotype structure evolve due to reduced recombination rate, on which the effects of selection can be overlaid. Our results highlight that distinct patterns of genetic variation can emerge through evolution of reduced local recombination rate. Recombination landscape as an evolvable trait therefore plays an important role determining the heterogeneous distribution of genetic variation along the genome.

## Introduction

Patterns of genetic variation in the genome represent ancestries of sequences and are influenced by population history. While genome-wide genetic variation represents population structure (McVean, 2009; Patterson et al., 2006), randomness in genealogies also contributes to fluctuation of local genetic variation along recombining chromosomes. Specifically, genealogies can differ between loci even under the same population history (Dutheil et al., 2009; Martin & Van Belleghem, 2017; McVean & Cardin, 2005; Pamilo & Nei, 1988; Wakeley, 2008, 2020; Wiuf & Hein, 1999). This is because realisation of a genealogy under a given population history is a probabilistic process: an ancestral haplotype for a set of individuals at one locus is not necessarily a common ancestor of the same set of individuals at another locus (Shipilina et al., 2023). Patterns of local genetic variation along the genome tend to conform with the population structure with random fluctuation (Fig. 1).

**Figure 1:**
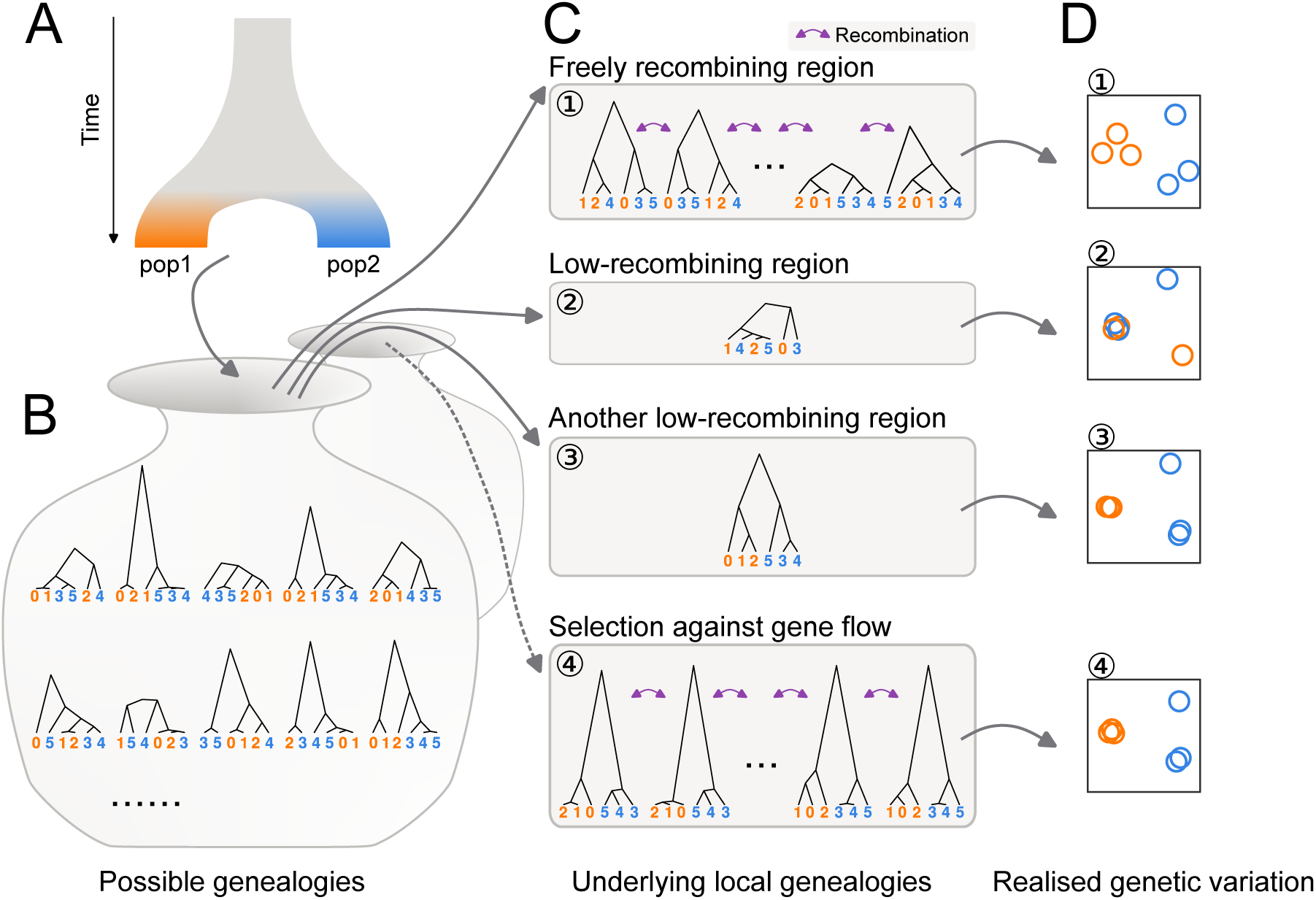
Distinct patterns of genetic variation can be due to reduced recombination rate. Population history (**A**) affects the distribution of possible genealogies (**B**) from which local genealogies are drawn (**C**). The number of genealogies in a genomic interval with a fixed physical length depends on the local recombination rate (**C**). Mutations occurring on the genealogies (not shown) determine the patterns of realised genetic variation. The realised genetic variation can be summarised and visualised with various methods such as PCA (**D**). (1) In freely recombining neutral regions, mutations represent many genealogies and hence the pattern of genetic variation converges to the population structure. (2, 3) In low-recombining neutral regions, mutations represent few genealogies covering the region leading to patterns of genetic variation distinct from the population structure. (3) Due to randomness in sampling of genealogies, some of such distinct patterns can be similar to patterns expected at targets of selective factors (c.f. 4). (4) At targets of selection, distribution of possible genealogies is different from that at neutral regions, which is depicted as a different set of possible genealogies in **B** and the dotted arrow.

Inference of population structure as well as other genome-wide analyses based on genetic variation take advantage of a sufficient number of unlinked variable sites (e.g. single nucleotide polymorphisms (SNPs)) to eliminate the effect of randomness. One of the most common methods to summarise population structure based on this approach is principal component analysis (PCA) applied on a whole-genome genotype table (McVean, 2009; Price et al., 2006). In a whole-genome PCA, variation among individuals based on variable sites of the entire genome are usually projected onto a few major axes (some analyses use many more axes), and the distances among individuals on these reduced dimensions represent genetic differences. Summarising population structure and other related measures using the entire genome has been proven to be an effective approach to eliminate random fluctuation of genealogies along the genome (Bhatia et al., 2013; Cao et al., 2020; Fedorova et al., 2013; Peter, 2022; Shao et al., 2023).

However, some fundamental biological questions concern selective factors that systematically bias the shape of genealogies at a genomic local scale, shifting the expected patterns of genetic variation from the population structure. For example, patterns of local genetic variation are distinct under selection against gene flow (Fig. 1C4), positive selection and adaptive introgression because they affect coalescence rate, topology, and branch lengths of the underlying genealogies (Hejase et al., 2020; Martin et al., 2015; Setter et al., 2020; Speidel et al., 2019; Wolf & Ellegren, 2017). Empirically, genome scans of population genetic summary statistics have been commonly used to identify regions with distinct patterns of genetic variation (Delmore et al., 2018; Irwin et al., 2018; Kawakami et al., 2017; Roesti et al., 2013; Rougemont et al., 2021). Many of these have identified regions with distinct patterns, such as elevated differentiation and reduced diversity, within low-recombining genomic regions (Geraldes et al., 2011; Kawakami et al., 2017; Renaut et al., 2013; Roesti et al., 2013, 2013; Rougemont et al., 2021). Distinct patterns at low-recombining regions can influence the chromosome-wide (Knief et al., 2016; Neafsey et al., 2010) and even genome-wide population structure (Mérot et al., 2021). These associations between distinct patterns of genetic variation at “outlier regions” or “genomic islands” and reduced recombination rate is often interpreted as linked selection (Burri et al., 2015; Burri, 2017; Delmore et al., 2015, 2018; Irwin et al., 2018; Kawakami et al., 2017; Roesti et al., 2013; Rougemont et al., 2021; Van Doren et al., 2017). However, a non-selective explanation is equally conceivable and yet often overlooked: the focal genomic region may contain too few underlying genealogies for a genome scan to eliminate the effect of random fluctuation simply due to low recombination rate, which is represented as the distinct patterns of genetic variation (Booker et al., 2020; Lotterhos, 2019). Specifically, it has not been well studied what aspects of distinct patterns of genetic variation can be explained by reduced recombination rate, and what other aspects reflect the effect of selection.

We address the effect of reduced recombination rate on local genetic variation using a songbird species, Eurasian blackcap (*Sylvia atricapilla*, hereafter “blackcap”), which is characterised by variability in seasonal migration across its distribution range (Berthold, 1988, 1991; Delmore et al., 2020a; Helbig, 1991). Populations with diverged migratory phenotypes split as recently as ∼30,000 years ago, likely corresponding to the last glacial period and now exhibit population structure (Fig. 2A-C, Sup. Fig. 1) (Delmore et al., 2020b). Due to their recent split and relatively large effective population size, genetic differentiation is very low among blackcap populations (Delmore et al., 2020b). The presence of population structure albeit with the low levels of differentiation makes the blackcap a perfect system to investigate local deviations of genetic variation: even the slightest effects of factors that change local genetic variation are likely detectable because such effects are not obscured by population structure. In addition, fine-scale recombination maps for multiple populations are available for this species (Bascón-Cardozo et al., 2022a), facilitating investigation of the relationship between changes in the recombination landscape and locally distinct patterns of genetic variation.

**Figure 2:**
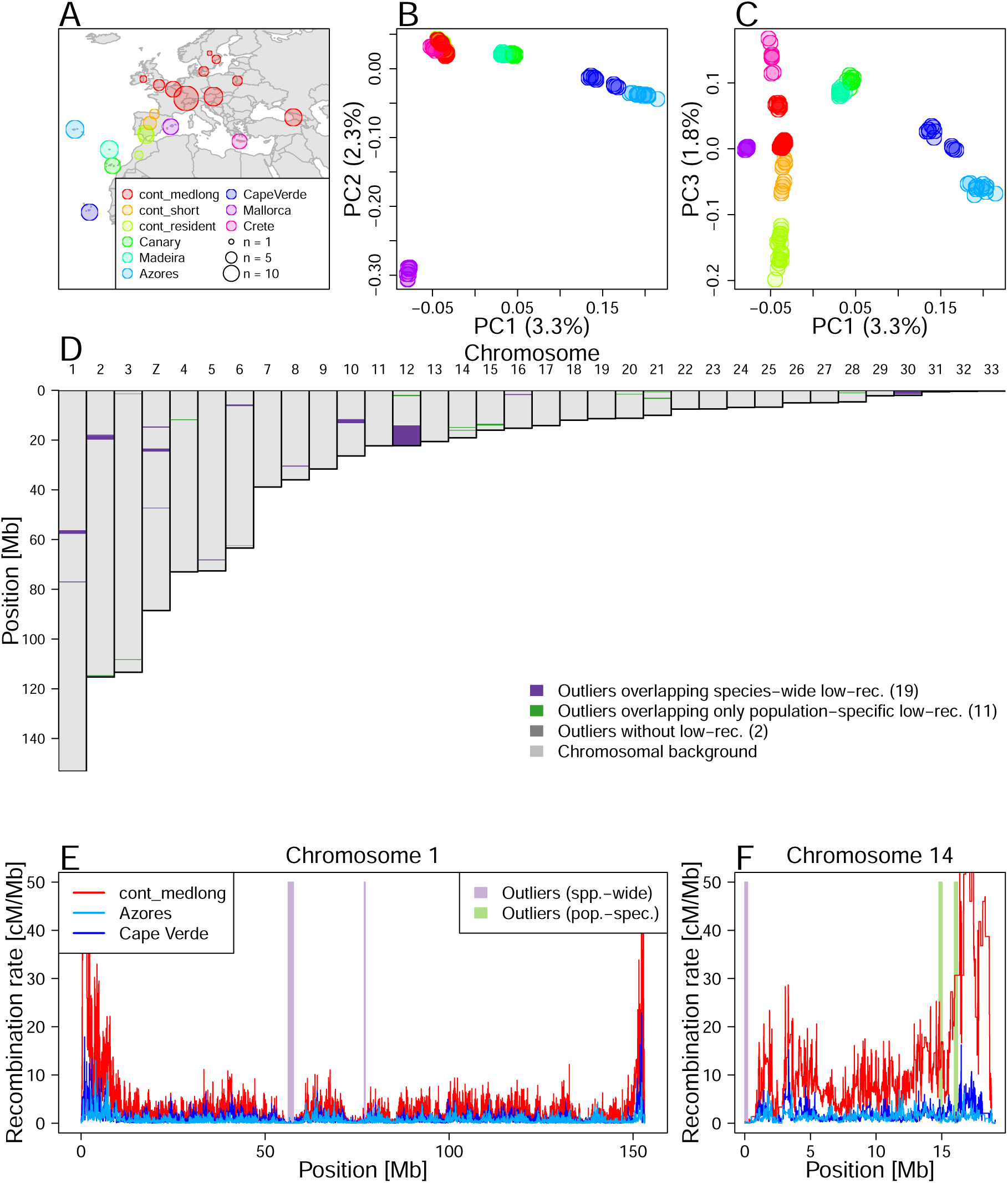
Local PCA outliers coincide with species-wide and population-specific low-recombining regions A. Geographic location of blackcap populations included in this study. Each point on the map represents a sampling location where multiple individuals were sampled. Populations were defined based on the geographic location, migratory phenotype, and genomic-wide population structure. **B, C.** Genome-wide PCA illustrating population structure. **D.** Distribution of outlier regions based on local PCA using `lostruct`. **E, F** Inferred recombination rates along two exemplified chromosomes (chromosomes 1 and 14) in three blackcap populations (cont_medlong, Azores, and Cape Verde). In **D-F**, purple and green shades respectively indicate positions of outliers that coincide with species-wide and population-specific low-recombining regions. The two green shades in **F** both overap with Azores and Cape Verde-specific low-recombining regions. cont_medlong: medium and long distance migrant population breeding on the continent; cont_short: short distance migrant population breeding on the continent; cont_res: resident (non-8migrant) population breeding on the continent. All island populations (Canary, Madeira, Azores, Cape Verde, Mallorca and Crete) are resident.

By leveraging a large-scale genomic re-sequencing dataset, we first systematically explore distinct patterns of local genetic variation along the blackcap genome, and compare these with genomic regions exhibiting reduced recombination rate. We further investigate the patterns of genetic variation in outlier regions and associate them with the prevalence of recombination suppression across populations. We also conduct simulations to analyse how reduced local recombination rate in the entire species and in a subpopulation with and without selection affects patterns of genetic variation through time. Finally, we propose a model of local genetic variation representing haplotype structure corresponding to evolutionary changes in local recombination rate.

## Results

### Chromosome-level reference assembly

To allow population genomic analyses in the blackcap system, we generated a chromosome-level reference genome using the Vertebrate Genomes Project pipeline v1.5 (Rhie et al., 2021). We collected blood of a female blackcap from Tarifa, Spain population. We generated contigs from Pacbio long reads, sorted haplotypes, and scaffolded them with 10X Genomics linked reads, Bionano Genomics optical mapping, and Arima Genomics Hi-C linked reads. Base call errors were polished with both PacBio long reads and Arrow short reads to achieve above Q40 accuracy (no more than 1 error every 10,000 bp). Manual curation identified 33 autosomes and Z and W chromosomes (plus 1 unlocalised W). Autosomes were named in decreasing order of size, and all had counterparts in the commonly used VGP reference zebra finch assembly (Sup. Table 2). The final 1.1 Gb assembly had 99.14% assigned to chromosomes, with a contig N50 of 7.4 Mb, and scaffold N50 of 73 Mb, indicating a high-quality assembly that fulfills the VGP standard metrics. The primary and alternate haplotype assemblies are provided under NCBI BioProject PRJNA558064, accession numbers GCA_009819655.1 and GCA_009819715.1.

### Deviation of genetic variation coincides with low-recombining regions

To investigate the genome-wide distribution of genetic variation, we mapped short reads of the whole-genomes of 179 blackcaps including 69 newly sequenced individuals (Sup. Table 1) on a *de novo*-assembled reference genome generated through the Vertebrate Genomes Project (VGP, Rhie et al., 2021), and called SNPs (Materials and Methods). To characterise genome-wide genetic variation, we performed PCA using SNPs in all autosomes, revealing population structure. While PC1 and PC2 represented differentiation of island populations (Fig. 2B), PC3 represented structure within continental populations with different migratory phenotypes (Fig. 2C). To identify genomic regions with patterns of genetic variation distinct from the population structure, we performed local PCA using `lostruct` (Li & Ralph, 2019). Briefly, `lostruct` performs PCA in sliding genomic windows and dissimilarity of PCA among windows are summarised with multidimensionality scaling (MDS). Distinct patterns of genetic variation of windows relative to the background are represented by extreme values along the MDS axes. Multiple windows with correlated patterns of genetic variation distinct from the population structure are represented by extreme values along the same MDS axis. This approach allowed systematic and unbiased exploration unaffected by our definition of populations of the blackcaps. We performed `lostruct` on both genotype and phased haplotype data with window size of 1,000 SNPs. We identified outlier windows by applying threshold MDS values (the mode of the distribution ± 0.3). We further identified genomic regions with distinct patterns of genetic variation by finding genomic intervals longer than 100 kb with at least five outlier windows based on the same MDS axis and merging the intervals based on the genotype- and phased haplotype-based approaches. This yielded 32 genomic regions with distinct patterns of variation (hereafter “outlier regions”, Fig. 2D, Sup. Table 3, Sup. Fig. 3). Their size ranged from 0.12 to 8.11 Mb (mean and median of 0.71 and 0.29 Mb), and each region contained 5,000 to 356,000 SNPs. Comparing the genomic distribution of these outlier regions to population-level recombination maps, we found that low-recombining regions (nominally recombination rate lower than the 20 percentile of each chromosome) were significantly enriched in the outlier regions (permutation tests with n = 1,000, p-value = 0.000 (Sup. Fig. 10)). Among these 32 outlier regions, 19 coincided with regions in which recombination rate was reduced in most tested populations (“species-wide” low-recombining regions), 11 coincided with regions in which recombination rate was reduced in one or two populations (“population-specific” low-recombining regions), and two did not coincide with low-recombining regions in any population (Fig. 2E, F, Sup. Fig. 9).

To further investigate the outlier regions, we separately performed PCA using SNPs in each region, revealing diverse patterns of distinct genetic variation (Fig. 3A-C top). First, species-wide low-recombining regions showed different levels of clustering of individuals in PCA. Specifically, the PCA projections consisted of either three distinct clusters (Fig. 3A top, Sup. Fig. 6), six loose clusters (Fig. 3B top, Sup. Fig. 6), or mixture of all individuals without apparent clustering (Sup. Fig. 6), suggesting that they represent haplotype structure with different numbers of low-recombining alleles. These clusters did not clearly separate populations, indicating a greater contribution of haplotype structure than the population structure. Four of these (e.g. Fig. 3A top, Sup. Figs. 6, 11) had the clearest clustering patterns with three groups of individuals in PCA, which is expected for a haplotype block with two distinct alleles (Huang et al., 2020; Ma & Amos, 2012; Todesco et al., 2020). Two of these regions showed LD patterns consistent with segregating inversions (Fig. 3A bottom, Sup. Fig. 12), and the other two showed patterns of non-inversion haplotype blocks (Sup. Fig. 12), indicating that recombination suppression with different mechanisms resulted in similar patterns of genetic variation due to presence of two distinct segregating haplotypes.

**Figure 3:**
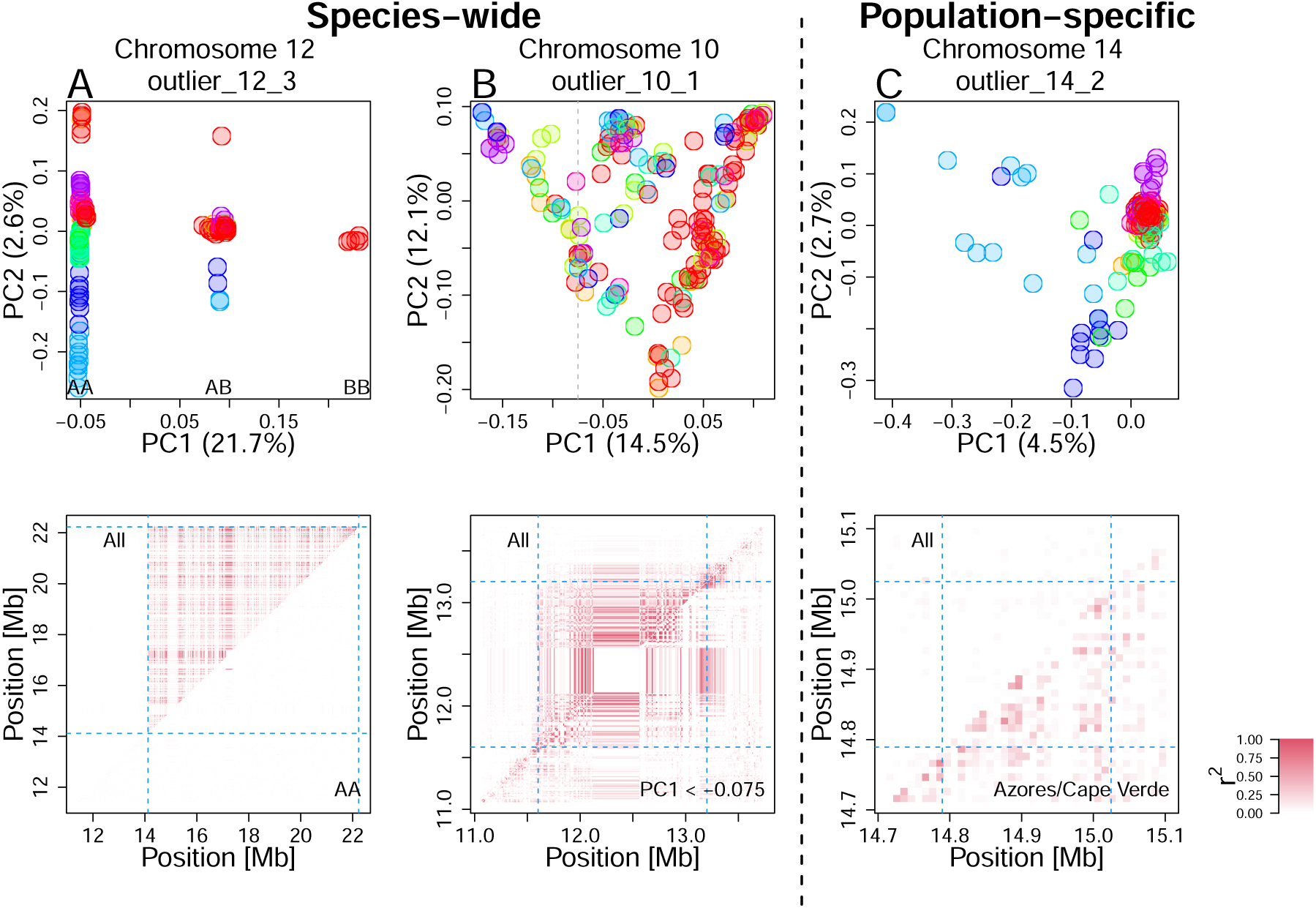
Patterns of genetic variation and linkage disequilibrium at local PCA outliers Top: PCA at exemplified outlier regions visualising the patterns of local genetic variation. Data points represent blackcap individuals colour-coded by population as depicted in Fig. 2. **Bottom**: LD calculated for all individuals (top-left diagonal) and for subset individuals (bottom-right diagonal). **A.** A putative inversion. Three clusters correspond to combination of two non-recombining alleles possessed by individuals, depicted as AA, AB, and BB. LD calculated using AA individuals is not elevated, in line with heterozygote-specific recombination suppression at an inversion locus (Sup. Fig.14). **B.** A species-wide low-recombining region with six loose clusters of individuals. LD calculated using subset individuals was elevated, suggesting genotype-non-specific recombination suppression. **C.** A population-specific low-recombining region. The variance in genetic distances between individuals of the low-recombining populations (Azores (blue) and Cape Verde (light blue)) is greater than between other pairs of individuals (top). LD calculated using individuals of the low-recombining populations is elevated (bottom).

Second, population-specific low-recombining regions exhibited distinct patterns of genetic variation consistently across the outlier regions. While individuals from the low-recombining populations were spread in PCA projections, individuals of other populations were more densely clustered (Fig. 3C top). This pattern indicates that the variance in genetic distances between a pair of individuals of the low-recombining populations is greater than between individuals of normally recombining populations. LD was elevated only in the low-recombining populations (Fig. 3C bottom), supporting population-specific reduction in recombination rate.

### Reduced recombination rate generates distinct patterns of genetic variation

To discern the effect of reduced recombination rate, demographic history, and unequal sample sizes among population on outlier regions, we performed neutral coalescent simulations using `msprime` (Baumdicker et al., 2022). We prepared 11 scenarios differing in the presence/absence of population subdivision, equal/unequal sizes of populations, presence/absence of gene flow between populations, and recombination rate in the middle of the chromosome relative to the chromosomal background (Sup. Fig. 19, Sup. Table 8). We applied `lostruct` on the simulated data to identify outlier regions. In all 1,000 replicates, reduced local recombnation rate resulted in distinct patterns of genetic variation irrespective of the population structure and demographic history (Sup. Fig. 20). We also asked whether population genetic summary statistics are affected. The mean nucleotide diversity (*π*), Tajima’s D, and F_ST_ were not affected, yet the variance of these statistics was greater within the low-recombining region than in the chromosomal background (Sup. Figs. 21, 22, 23).

To address how species-wide and population-specific reduction in recombination rate affect the patterns of genetic variation over time, we performed forward simulations using `SLiM` (Haller & Messer, 2022). First, to investigate the effects of species-wide reduction in local recombination rate, we simulated one ancestral population of 1,000 diploids with a low-recombining genomic region that splits into three subpopulations (pop1, pop2, pop3. Fig. 4A). We sampled individuals over time after the populations split and conducted PCA both in the low-recombining and normally recombining genomic regions. PCA patterns at low-recombining regions (Fig. 4B, C, Sup. Fig. 24) were distinct from normally recombining regions (Fig. 4D). The low-recombining regions exhibited three, six, or more clusters of individuals resembling our empirical results. The clusters of individuals represented genotypes consisting of different combinations of ancestral haplotypes (Sup. Fig. 25). The distinct patterns representing haplotype structure persisted until population structure started to emerge along the PC axes (Fig. 4B, C). Accordingly, the percentages of variation explained by PC1 and PC2 were higher at low-recombining regions than in normally recombining region until this transition (Fig. 4C). Distinct patterns in the low-recombining regions persisted over longer times than it took for population structure in normally recombining region to emerge (Fig. 4D). These results suggest that distinct patterns of genetic variation in species-wide low-recombining regions represent haplotype structure whose transition to the population structure is slower than in normally recombining regions.

**Figure 4:**
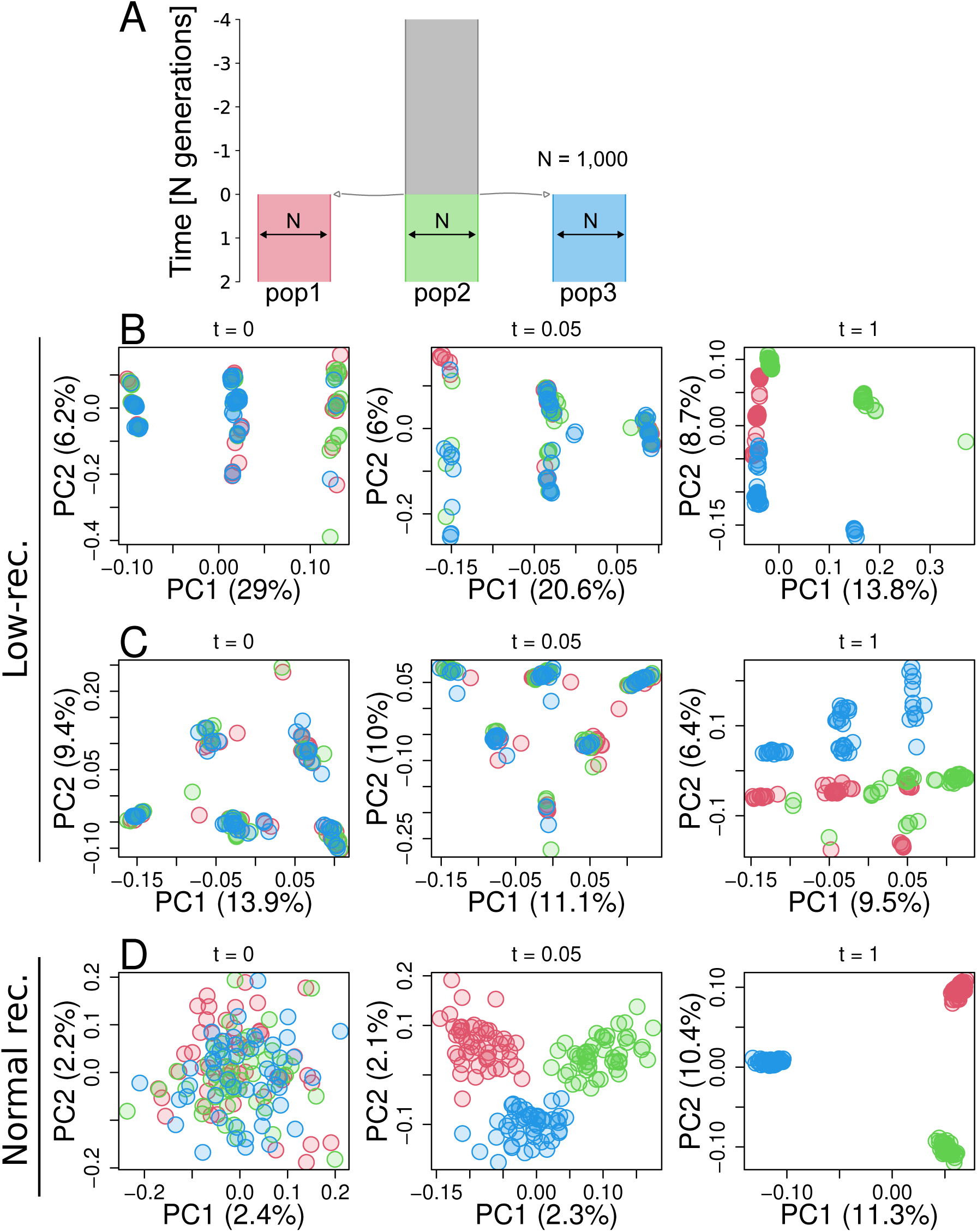
Simulation of a species-wide low-recombining region. **A.** Simulated demography scenario. Our simulated genome contained two chromosomes, one with a low-recombining region and the other without. **B, C.** PCA showing patterns of genetic variation at the species-wide low-recombining region at three time points in three exemplified simulation replicates. **D.** PCA showing patterns of genetic variation at a normally recombining chromosome at three time points in the same replicates as **B**.

Second, to investigate the effects of population-specific reduction in local recombination rate, we performed forward simulations. Three populations (pop1, pop2, and pop3) and their ancestral population had 1,000 diploid individuals, and pop1 evolved a reduced local recombination rate. We considered two cases with respect to when the population-specific reduction in recombination rate is introduced: before or after differentiation of populations. In the first scenario (Sup. Fig. 26), recombination suppression was introduced at the same time as the three populations split, while in the second scenario (Fig. 5A) recombination suppression was introduced 4,000 generations after the split. We conducted PCA in genomic regions with and without population-specific recombination suppression using individuals sampled over time. In both scenarios, the genomic region with population-specific recombination suppression transiently showed distinct patterns of genetic variation (Fig. 5B, Sup. Fig. 26B) resembling the empirical results, while regions without population-specific suppression showed population structure (Fig. 5C). Haplotype structure was not as conspicuous as in species-wide low-recombining regions (Sup. Fig. 27B, F, c.f. Sup. Fig. 25) due to standing genetic variation. Mutations originating in the non-recombining population were enriched in the set of mutations that have the greatest contribution to the distinct pattern of PCA (Sup. Fig. 27C, G. *χ*^2^ tests, p-value = 1.14 × 10*^−^*^12^ for model 1 and p-value = 2.30 × 10*^−^*^32^ for model 2). These mutations were significantly associated with each other in the underlying genealogy sharing common branches compared to other mutations originating in the same population (Sup. Fig. 27D, H. Materials and Methods, Kolmogorov-Smirnov tests, p-value = 7.74 × 10*^−^*^6^ for model 1 and p-value = 0.0012 for model 2), indicating that the distinct pattern of genetic variation represents sets of mutations that occurred in ancestral haplotypes. Associations between these population-specific mutations on ancestral haplotypes would have eventually decayed by recombination events, but in the low-recombining population the association was maintained due to suppressed recombination, resulting in the cryptic haplotype structure.

**Figure 5:**
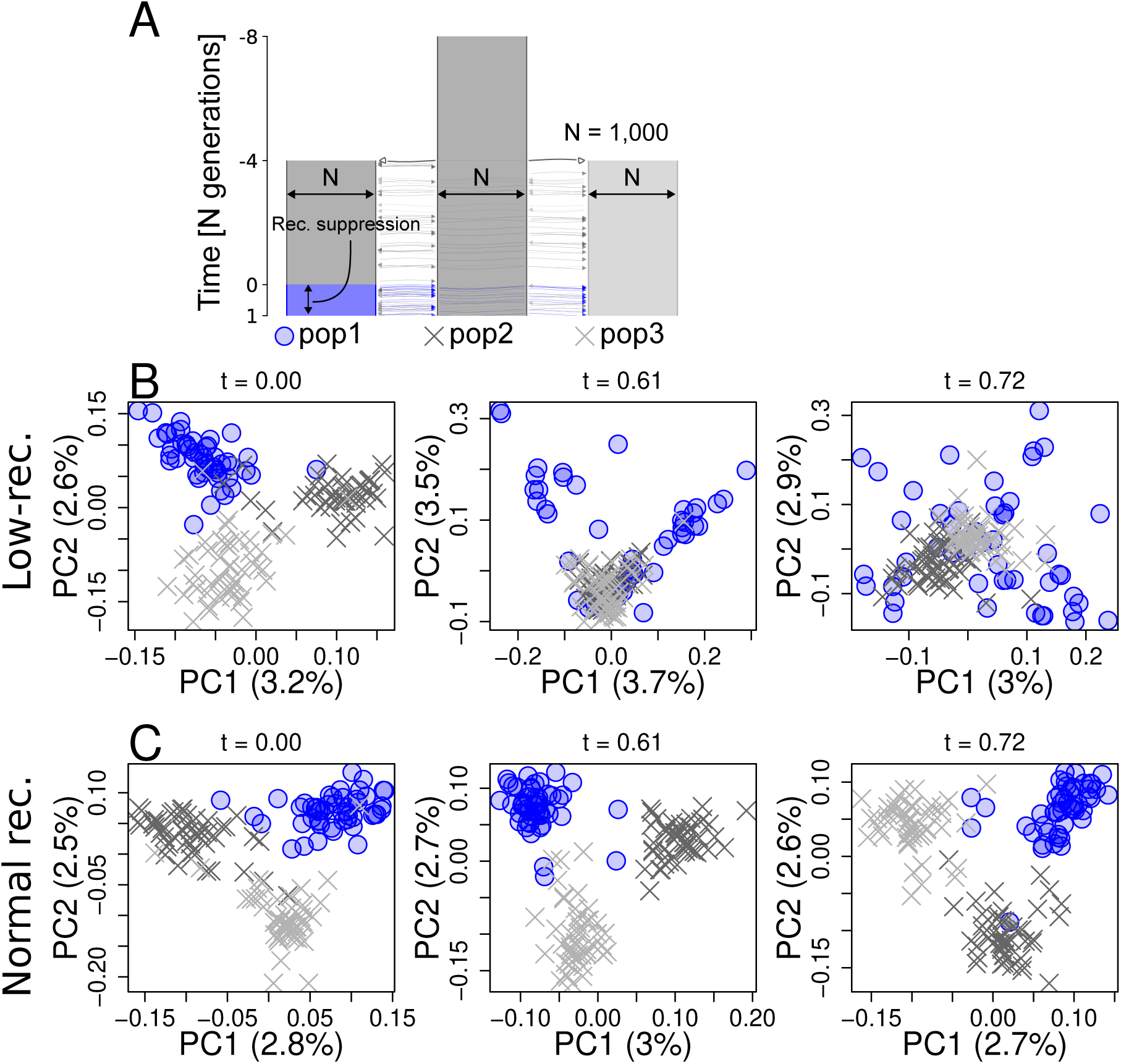
Simulation of a population-specific low-recombining region. **A.** Simulated scenario. Simulated genome contained two chromosomes, one with a population-specific low-recombining region and the other without. **B, C.** PCA showing patterns of genetic variation at the population-specific low-recombining region (**B**) and the normally recombining chromosome (**C**) at three time points in one exemplified simulation replicate.

### Effect of selection on patterns of genetic variation

Selection is known to cause distinct patterns of genetic variation (Nielsen, 2005). To test whether the outlier regions based on `lostruct` identified in the blackcap genome are also targets of selection, we measured nucleotide diversity (*π*) and Tajima’s D in each population, as well as ratio between non-synonymous and synonymous substitutions (d_N_*/*d_S_) for annotated genes. Many species-wide low-recombining regions showed reduced nucleotide diversity (Sup. Fig. 29) and Tajima’s D (Sup. Fig. 28), suggesting that they are under either positive or purifying selection. Most genes within outlier regions had d_N_*/*d_S_ below 0 (Sup. Fig. 30) with a few genes with positive d_N_*/*d_S_, indicating that most genes are under purifying selection and a few others are under positive selection. Furthermore, sequence analysis indicated that some but not all species-wide low-recombining outlier regions coincide with putative pericentromeric regions with enrichment of long tandem repeats (Sup. Figs. 33, 34). These results indicate that the outlier regions may experience effects of selection in addition to reduced recombination rates.

We asked whether the distinct patterns of local genetic variation at the outlier regions observed in blackcaps represent the effect of selection instead of reduced recombination rates. Specifically, we addressed wheather the distinct patterns of genetic variation representing haplotype structure could be caused by (i) purifying or (ii) positive selection alone or if they primarily represent the effect of reduced recombination rate. To this end, we used `SLiM` to simulate purifying and positive selection with and without reduction in recombination rate, and investigated local genetic variation over time by PCA. First, to investigate the effect of purifying selection, we simulated two chromosomes with and without a species-wide low-recombining region under the same demographic history as the neutral scenario (Fig. 4A) but with different strength of purifying selection by introducing mutations with different ratios between the rates of neutral and deleterious mutations (Materials and Methods). Distinct patterns of genetic variation representing haplotype structure evolved only in scenarios where recombination rate was reduced irrespective of the distribution of fitness effects (DFE) (Sup. Fig. 31). Stronger purifying selection (DFE with more frequent deleterious mutations in our simulation) decreased the time for distinct patterns of genetic variation at low-recombining regions to be overtaken by population structure (Sup. Fig. 31A, C). Second, to investigate the effect of positive selection, we simulated a chromosome with or without a species-wide low-recombining region under the same demographic history, and introduced a beneficial mutation 100 generations after the population split in one population (Sup. Fig. 32A) or 100 generations before the split in the ancestral population (Sup. Fig. 32D). For simulations in which the beneficial mutation persisted, we recorded the patterns of local genetic variation by PCA over time. Although positive selection affected patterns of genetic variation compared to the neutral scenario, distinct patterns of genetic variation representing discrete haplotypes were unique to scenarios with reduced recombination rate in both cases (Sup. Fig. 32B-E). These results indicate that distinct patterns of genetic variation represented in local PCA, as in the blackcap outlier regions, primarily reflect haplotype structure due to reduced recombination rate, on which the effect of selection can be overlaid.

## Discussion

### Distinct patterns of genetic variation at low-recombining regions: Genealogical interpretations

#### Genealogical noise, genealogical bias, and mutational noise

A number of empirical population genomics studies have identified ecologically and evolutionarily important genomic regions by locating outlier regions with distinct patterns of genetic variation (Jones et al., 2012; Lamichhaney et al., 2016; Lawniczak et al., 2010; Lundberg et al., 2021; Malinsky et al., 2015). Genomic windows in such studies are assumed to be both large enough to eliminate the effect of random fluctuation in local genetic variation and small enough to capture the localised signatures of selection. We showed empirically that genomic regions with distinct patterns of genetic variation identified by a population genomic scan based on principal component analysis (PCA) highly overlap with low-recombining genomic regions (Fig. 2). With simulations, we showed that although selection may affect the amount and pattern of local genetic variation around the target locus, the distinct patterns of genetic variation represented by PCA at low-recombining regions can be primarily explained by haplotype structure due to reduced recombination rate (Figs. 4, 5). We discuss our findings from the perspective of underlying genealogies.

We first define three terms: (1) genealogical noise, (2) genealogical bias, and (3) mutational noise. (1) By “genealogical noise” we refer to the fact that gene genealogies vary along the genome following a null distribution given a population history (Dutheil et al., 2009; Martin & Van Belleghem, 2017; McVean & Cardin, 2005; Wakeley, 2008, 2020; Wiuf & Hein, 1999). (2) By “genealogical bias” we refer to the fact that selective processes can systematically shift the distribution of local genealogies away from the null distribution. For example, genealogies under positive selection, selection against gene flow, adaptive introgression, and balancing selection are biased due to bursts of coalescence, faster lineage sorting, and introduction and maintenance of long branches (Barton & Etheridge, 2004; Guerrero et al., 2012; Hejase et al., 2020; Martin et al., 2019; Setter et al., 2020; Speidel et al., 2019; Taylor, 2013). On top of these, (3) randomness in the process of mutation causes additional noise in realised genetic variation (Ralph et al., 2020), which we call “mutational noise”. For example, the first and the second halves of a chromosomal interval with a single genealogy can still have slightly different patterns of genetic variation because they represent some finite numbers of different mutations.

#### Species-wide low-recombining regions

We showed in blackcaps that some distinct patterns of genetic variation are associated with species-wide low-recombining regions (Fig. 2). This is in line with previous studies reporting negative correlation between recombination rate and genetic differentiation (Burri et al., 2015; Burri, 2017; Delmore et al., 2015, 2018; Irwin et al., 2018; Kawakami et al., 2017; Roesti et al., 2013; Rougemont et al., 2021; Van Doren et al., 2017). To investigate what factors affect distinct patterns of gentic variation at low-recombining regions (Fig. 3) in more detail, we performed simulations of low-recombining regions with and without selection, and demonstrated that haplotype structure underlies the distinct patterns which persists only transiently until the effect of the population structure emerges (Figs. 4, 5). This transiency reflects a shift from local genetic variation primarily representing haplotype structure (Lotterhos, 2019; Ma & Amos, 2012) to that representing population structure, which can be interpreted based on the underlying genealogies. Low-recombining regions have few underlying genealogies per interval of a fixed physical length and haplotype structure at such regions tends to reflect their basal branches because basal branches tend to be longer than peripheral branches (Wakeley, 2008). At a time point soon after a population split event, peripheral branches covering more recent times than the population split harbour fewer mutations than basal branches. Therefore, the realised pattern of genetic variation at this stage has the greatest contributions by mutations on the long basal branches undifferentiated among populations (i.e. consisting standing genetic variation), representing a few ancestral haplotypes that descend the current sample. As time passes after the population split, the proportion of mutations that have occurred after the population split increases while some ancestral haplotypes can be lost by chance (i.e. drift), increasing the contribution of population structure on genetic variation. This type of distinct patterns of genetic variation arises predominantly in low-recombining regions but less so in normally recombining regions. This is because haplotype structure representing a few ancestral lineages would become less prominent with recombination as different segments of a current haplotype can follow distinct ancestries and thus the genealogical noise is effectively averaged out.

Some low-recombining regions may have genealogies with much shorter basal branches than other low-recombining regions because the variance in the basal branch length is greater than peripheral branches (Wakeley, 2008). The over-representation of a few ancestral haplotypes in genetic variation requires long basal branches in the underlying genealogies, and thus low-recombining regions with relatively short basal branches cannot accommodate sufficient mutations to represent distinct ancestral haplotypes. This decreases the relative contribution of genealogical noise compared to mutational noise (Supplementary Notes 1.1). Distinct patterns of genetic variation with varying levels of clustering of individuals in PCA in our empirical results (Sup. Fig. 6) may correspond to different ratios between genealogical and mutational noise due to large variance in the basal branch lengths of underlying genealogies. Specifically, some outlier regions with mixture of individuals from multiple populations without distinct clusters and population subdivision in PCA may have underlying genealogies with short basal branches leading to greater contributions of mutational noise on the realised genetic variation.

#### Population-specific low-recombining regions

We both empirically and with simulations showed that population-specific low-recombining regions exhibit distinct patterns of genetic variation in which individuals of low-recombining and normally recombining populations have different variance in genetic distances (Fig. 3C, Fig. 5). This unequal variance in low-recombining and normally recombining populations can be interpreted based on the underlying genealogies (Sup. Fig. 35). We consider the ancestry of current samples of low-recombining and normally recombining populations and split the ancestry at the time *T* when the population-specific recombination suppression initiated (Sup. Fig. 35A). At time *T*, there were *n*_1_ and *n*_2_ ancestral haplotypes that descend all current samples in low-recombining and normally recombining populations. At times older than *T*, the ancestors of the *n*_1_ and *n*_2_ haplotypes may freely recombine within each set, making the genetic distances among ancestral haplotypes within each population close to equidistant (Sup. Fig. 35B). After the initiation of the population-specific reduction in recombination rate, the ancestry of one current sequence of the low-recombining population can be traced back to either one of the *n*_1_ ancestral haplotypes present at the time *T* (Sup. Fig. 35A). On the contrary, the ancestry of one current sequence of the normally recombining population can be traced back to multiple ancestral haplotypes of the *n*_2_ sequences because of the presence of recombination (Sup. Fig. 35A). From the perspective of mutations, in the low-recombining population, mutations that arose on the same haplotype tend to be linked until the present time because of the suppressed recombination. On the other hand, in the normally recombining population, mutations that arose on the same ancestral haplotype less likely stay linked until the present time because recombination can dissociate them. Because shuffling of haplotypes reduces the variance of genetic distances among sequences, population-specific reduction in recombination rates leads to greater variance in low-recombining population than in normally recombining population as observed in our empirical results and simulations. In short, because of the different recombination rates between the populations, genealogical noise is more efficiently eliminated in the normally recombining population than in the low-recombining population.

The haplotype structure at population-specific low-recombining region is only cryptic and less apparent than in species-wide low-recombining regions because other standing mutations coexist on the same haplotype, which are older than the initiation of the population-specific recombination suppression (Sup. Fig. 27). The elevated PC loadings at linked mutations originating in the low-recombining population could be informative to study evolutionary change in local recombination rate: the ages of such mutations mapped on inferred genealogies might be useful to estimate the timing at which the population-specific recombination suppression initiated.

In our empirical analyses in blackcaps, we detected the effect of population-specific reduction of recombination rate in Azores and Cape Verde island populations (Fig. 3C, Sup. Fig. 7). It remains unclear why reduced recombination rate in certain populations but not others is reflected as distinct patterns of genetic variation by `lostruct`. The recent split of Azores and Cape Verde populations from other populations, accompanied by reduction in population size and the level of isolation (Delmore et al., 2020b) may have contributed to more efficient spread of reduced recombination rate.

### Recombination landscape as a driver of evolution of local genetic variation

Species-wide and population-specific recombination suppression underlying distinct patterns of local genetic variation are probably not independent: reduction in recombination rates that initiates formation of haplotype blocks likely originates from one population and may spread to multiple populations. For example, local recombination rate may be initially reduced in one population in which a segregating inversion originates before it may spread in multiple populations by gene flow (Faria et al., 2019). In line with this view of recombination map as an evolvable trait diverging across populations according to subdivision, recent studies find that divergence in local recombination rate among populations is correlated with genetic divergence (Bascón-Cardozo et al., 2022a; Roesti et al., 2013; Spence & Song, 2019). Future work on the effects of transition from population-specific to species-wide suppression of recombination will fill the gap between the two states.

Besides spread of recombination suppression across populations, there are other paths along which patterns of local genetic variation may change over time. First, change in frequency of one haplotypic variant by drift or gene flow and selection and accumulation of novel mutations may shift the distinct pattern of genetic variation (Rubin et al., 2022). Second, an increase in recombination rate in the region may resolve the distinct pattern of genetic variation and result in emergence of the population structure, because recombination breaks down discrete haplotypes and generates mixed types whereby reducing the variance of genetic variation (Hudson, 1983). These two types of shifts in distinct patterns of genetic variation are not mutually exclusive. For example, fixation of an inversion results in elevated recombination rate (Smukowski Heil et al., 2015; Stevison et al., 2011) because there are no longer non-recombining heterozygotes in the population. Due to resumed recombination, patterns of local genetic variation in such regions are expected to reflect population structure eventually. The question of how long it takes for an outlier region with distinct patterns of genetic variation to disappear after these events should be focally studied in the future.

In Fig. 6A, we illustrate a model for the evolution of local genetic variation that changes according primarily to the evolution of local recombination rates. Local genetic variation can become distinct from the population structure first by representing emerging haplotype structure associated with population-specific recombination suppression or other types of haplotype blocks (e.g. inversions) in one population. If this recombination suppression spreads throughout all populations, then local genetic variation will start to reflect species-wide haplotype structure. Once the relative contribution of haplotype structure on local genetic variation is reduced by differentiation or disappears by elevated recombination rates, then genetic variation returns to reflect the population structure and consequently the outlier region disappears. The effect of selection on local genetic variation may be overlaid on top (Supplementary Notes 1.2).

**Figure 6:**
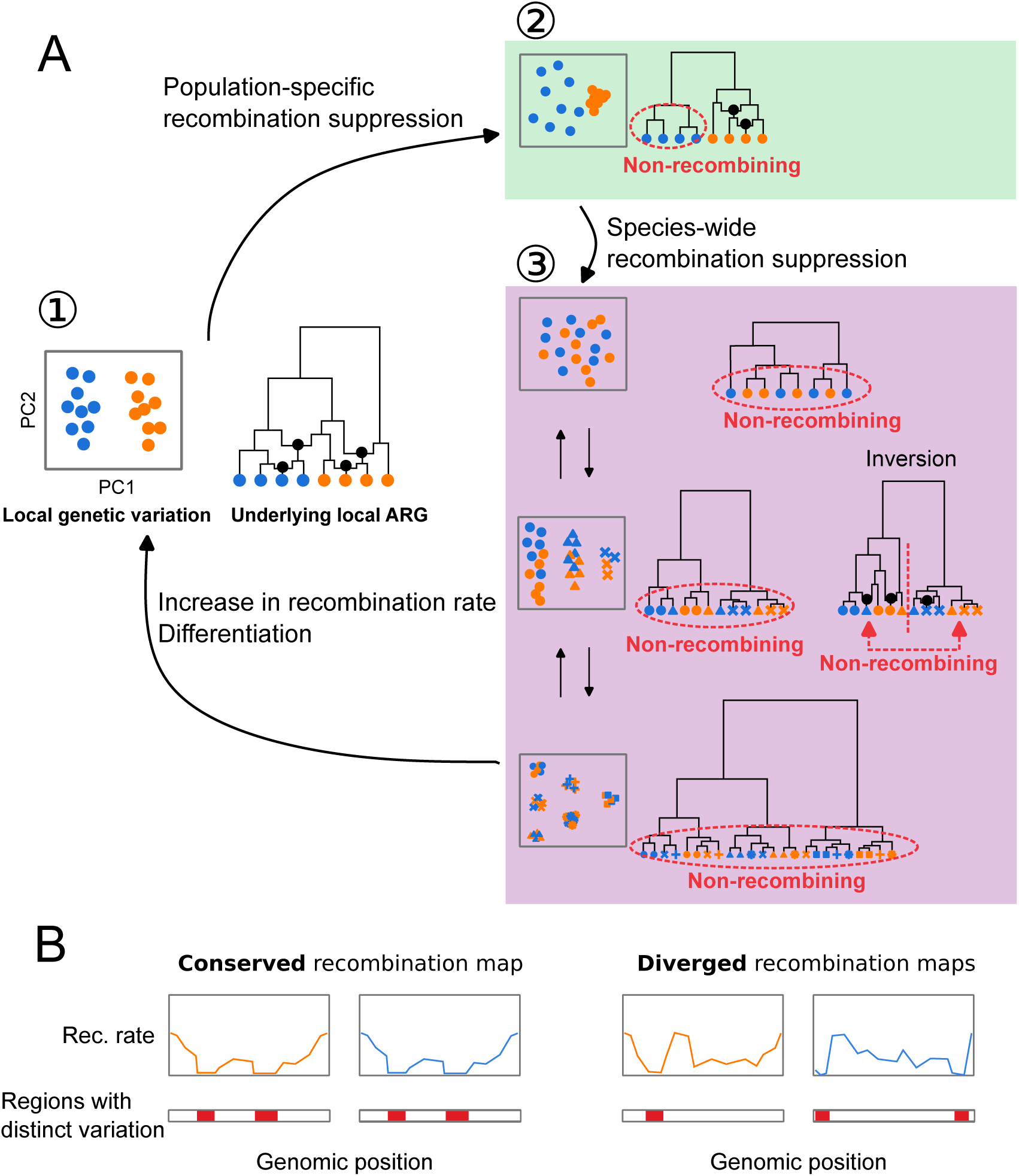
Evolutionary changes in local recombination rate influence evolution of local genetic variation. **A.** Local genetic variation is shown in hypothetical PCA plots. Their underlying genealogies are shown in simplified ancestral recombination graphs (ARGs,(Griffiths & Marjoram, 1997; reviewed in Lewanski et al., 2024)), on which black dots represent ancestral recombination events contributing to the sampled sequences. Points in PCA depict diploid individuals, while those on the ARGs represent haploid sequences. Two colours of these points (blue and orange) indicate two populations. (1) Local genetic variation concordant to population structure. Genetic variation shows separation of individuals from two populations. ARG shows that recombination is suppressed in neither population. (2) Population-specific recombination suppression in the blue population. ARG shows that recombination is suppressed in the blue population. (3) Species-wide recombination suppression. Top: A case in which there are few mutations representing the basal splits of the underlying genealogy at species-wide low-recombining region. Middle: A case in which there are two haplotypic variants at the species-wide low-recombining region. If this is due to presence of an inversion (right ARG), recombination is suppressed between but not within the two clades representing two alleles. Bottom: A case in which there are three haplotypic variants at the species-wide low-recombining region. **B** Evolution of recombination map influences differe2n3ce in genomic distributions of distinct patterns of genetic variation between species/populations.

### Implications

Finally, we discuss technical and biological implications of our study. The technical implication concerns interpretation of genome scans based on local genetic variation. A number of methods based on local genetic variation have been used to detect loci involved in different kinds of selective processes. For example, F_ST_ (differentiation), d_XY_ (divergence), and other population parameters are inferred to detect genomic islands of speciation (Delmore et al., 2018; Hejase et al., 2020; Huang et al., 2020; Malinsky et al., 2015). Reduced diversity (*π*) is a signature of selection (Delmore et al., 2018; Irwin et al., 2018; Pracana et al., 2017), and by combining it with variation among populations, loci associated with population-specific selection can be also inferred (Yi et al., 2010). Targets of adaptive introgression have been identified by applying statistics based on ABBA-BABA test, which is related to genetic variation (Peter, 2016, 2022), in sliding windows (Kronforst et al., 2013; Martin et al., 2015; Patterson et al., 2012; Reich et al., 2009). However, there are confounding factors that affect inference of these statistics. For example, it has been shown that low diversity can cause elevation in some of these statistics (Cruickshank & Hahn, 2014; Noor & Bennett, 2009). In addition to reduced diversity, this study and others (Booker et al., 2020; Lotterhos, 2019; Renaut et al., 2013) show that reduced recombination rate also causes distinct patterns of genetic variation which can lead to erroneous identification of regions under influence of selective factors. Examining recombination rates at identified regions and comparing them to other regions are necessary to avoid this. For instance, apparent outliers in only few (pairs of) populations at a low-recombining region may reflect high variance, while high variance at low-recombining regions alone cannot explain signals occurring in many (quasi-) independent populations or species at a low-recombining region. Furthermore, corroborating methods based on different aspects of distinct patterns of variation, such as site frequency spectrum (DeGiorgio et al., 2016; Fay & Wu, 2000; Tajima, 1989), LD (Sabeti et al., 2002, 2007; Voight et al., 2006), inferred genealogies (Hejase et al., 2020; Speidel et al., 2019; Stern et al., 2019), local landscape of variation (Setter et al., 2020), and sites of mutations in genes (Nei & Gojobori, 1986), as well as approaches with explicit simulation based on inferred demography (Hager et al., 2022), may be informative.

The biological implication is about evolution of recombination rates and genetic variation along the genome. Based on our findings of a link between these, we predict that organisms with more conserved recombination landscape along the genome may have more conserved genomic landscapes of distinct patterns of genetic variation (Fig. 6B). In other words, the more conserved recombination maps are, the more correlated genomic distribution of distinct genetic variation may be between species. In vertebrates including placental mammals (with some exceptions), recombination landscape along the genome evolves fast due to continuous turnovers of alleles of PRDM9 (the gene coding a protein that determines recombination hot spots) and its target DNA sequences (Baudat et al., 2010; Myers et al., 2008). For instance, in mammals that possess functional PRDM9, the genomic landscape of recombination rates is distinct between and even within species (Kong et al., 2010; Spence & Song, 2019; Stevison et al., 2016). Importantly, PRDM9 has been pseudogenised (Birtle & Ponting, 2006) or lost (Baker et al., 2017) independently in multiple vertebrate lineages. This shifted the determinants of recombination map from the PRDM9 allele and its target to genomic features such as CpG islands and transcription start sites, stabilising the recombination landscape (Auton et al., 2013; Baker et al., 2017; Singhal et al., 2015). Our results shown in birds, a group lacking PRDM9 (Birtle & Ponting, 2006; Singhal et al., 2015), raises a question whether the evolution of local recombination rates may play an even more important role in shaping local genetic variation in organisms with functional PRDM9. Comparative studies using taxa with and without functional PRDM9 will address this and may link the evolution of genomic landscape of distinct patterns of genetic variation and (in)stability of recombination maps.

## Materials and Methods

### Empirical analyses

#### *de novo* genome assembly

A chromosome-level blackcap reference genome was *de novo* assembled within the Vertebrate Genomes Project (VGP), following pipeline version 1.5 (Rhie et al., 2021). In brief, blood of a female blackcap from the resident Tarifa population in Spain was collected in 100% ethanol on ice and stored at −80 °C (NCBI BioSample accession SAMN12369542). The ethanol supernatant was removed and the blood pellet was resuspended in Bionano Cell Buffer in a 1:2 dilution. Ultra-long high molecular weight (HMW) DNA was isolated using Bionano agarose plug method (Bionano Frozen Whole Nucleated Blood Stored in Ethanol – DNA Isolation Guidelines (document number 30033)) using the Bionano Prep Blood and Cell Culture DNA Isolation Kit. Four DNA extractions were performed yielding a total of 13.5 µg HMW DNA. About 6 µg of DNA was sheared using a 26G blunt end needle (PacBio protocol PN 101-181-000 Version 05) to ∼40 kb fragments. A large-insert PacBio library was prepared using the Pacific Biosciences Express Template Prep Kit v1.0 following the manufacturer protocol. The library was then size selected (>15 kb) using the Sage Science BluePippin Size-Selection System. The library was then sequenced on 8 PacBio 1M v3 smrtcells on the Sequel instrument with the sequencing kit 3.0 and 10 hours movie with 2 hours pre-extension time, yielding 77.51 Gb of data (∼66.29X coverage) with N50 read length averaging around 22,927 bp. We used the unfragmented HMW DNA to generate a linked-reads library on the 10X Genomics Chromium (Genome Library Kit & Gel Bead Kit v2, Genome Chip Kit v2, i7 Multiplex Kit PN-120262). We sequenced this 10X library on an Illumina Novaseq S4 150 bp PE lane to ∼60X coverage. Unfragmented HMW DNA was also used for Bionano Genomics optical mapping. Briefly, DNA was labeled using the Bionano Prep Direct Label and Stain (DLS) Protocol (30206E) and run on one Saphyr instrument chip flowcell. 136.31 Gb of data was generated (N50 = 301.9kb with a label density = 16.91 labels/100kb). Optical maps were assembled using Bionano Access (N50 = 27.48 Mb and total length = 1.41 Gb). Hi-C libraries were generated by Arima Genomics and Dovetail Genomics and sequenced on HiSeq X at ∼60X coverage following the manufacturer’s protocols. Proximally ligated DNA was produced using the Arima-HiC kit v1, sheared and size selected (200 – 600 bp) with SRI beads, and fragments containing proximity-ligated DNA were enriched using streptavidin beads. A final Illumina library was prepared using the KAPA Hyper Prep kit following the manufacturer guidelines. FALCON v1.9.0 and FALCON unzip v1.0.6 were used to generate haplotype phased contigs, and purge_haplotigs v1.0.3 was used to further sort out haplotypes (Guan et al., 2020). The phased contigs were first scaffolded with 10X Genomics linked reads using scaff10X 4.1.0 software, followed with Bionano Genomics optical maps using Bionano Solve single enzyme DLS 3.2.1, and Arima Genomics in-vitro cross-linked Hi-C maps using Salsa Hi-C 2.2 software (Ghurye et al., 2019). Base call errors were polished with both PacBio long reads and Arrow short reads to achieve above Q40 accuracy (no more than 1 error every 10,000 bp). Manual curation was conducted using gEVAL software by the Sanger Institute Curation team (Howe et al., 2021). Curation identified 33 autosomes and Z and W chromosomes (plus 1 unlocalised W). Autosomes were named in decreasing order of size, and autosomes 1 through 30 and sex chromosomes had counterparts in the commonly used VGP reference zebra finch assembly (Sup. Table 2). The total length of the primary haplotype assembly was 1,066,786,587 bp, with 99.14% assigned to chromosomes. The final 1.1 Gb assembly consisted of 601 contigs in 189 scaffolds, with a contig N50 of 7.4 Mb, and scaffold N50 of 73 Mb, indicating a high-quality assembly that fulfills the VGP standard metrics.

#### Whole-genome resequencing

We resequenced 69 blackcap samples from various populations across the species distribution range (Sup. Table 1) to complement an existing dataset of 110 blackcaps, 5 garden warblers, and 3 African hill babblers that had been sequenced previously (Delmore et al., 2020b). Blood samples from the additional 69 blackcaps were collected from the brachial vein and stored in 100% ethanol. High molecular weight genomic DNA was extracted with a standard salt extraction protocol or through the Nanobind CBB Big DNA Kit Beta following the manufacturer’s instructions. Libraries for short insert fragments between 300 and 500 bp were prepared and were then sequenced for short paired-end reads on either Illumina NextSeq 500, HiSeq 4000 or NovaSeq 5000 (Sup. Table 1).

We performed quality control of the reads with FastQC version 0.11.8 (https://www.bioinformatics.babraham Reads from all samples were mapped against the blackcap reference genome following an adjusted pipeline of `Genome Analysis Toolkit` (`GATK` version 4.1.7.0, McKenna et al. (2010)) and `Picard` version 2.21.9 (http://broadinstitute.github.io/picard/). After resetting the base quality of adapter bases in the sequenced reads to 2 with `Picard MarkIlluminaAdapters`, paired-end reads were mapped to the reference using `BWA mem` (Li, 2013). To ensure that both unmapped mates and secondary/supplementary reads were marked for duplicates, we ran `Picard MarkDuplicates` for sorted reads with the default pixel distance of 100 for reads from Illumina NextSeq 500 or with a pixel distance of 2,500 for reads from HiSeq 4000 and NovaSeq 5000. Due to low coverage, 10 samples (Sup. Table 1) were sequenced multiple times. Alignment files for these samples (in BAM format) were merged with `Picard MergeSamFiles`. Per-sample quality control of BAM files were performed using `QualiMap` version 2.2.1 (Okonechnikov et al., 2016), `Picard CollectMultipleMetrics`, `CollectRawWgsMetrics` and `CollectWgsMetrics`; and `MultiQC` version 1.8 (Ewels et al., 2016). The minimum and median depth were 7.8X and 20.1X, and the minimum and median coverage were 0.88 and 0.97. We called bases at all positions per sample using `GATK HaplotypeCaller`. We combined gVCF files of 189 individuals into ten evenly sized subsets (to allow parallelisation of the following variant calling step) with `GATK CombineGVCFs`. We genotyped SNPs and INDELs using `GATK GenotypeGVCFs`, and the 10 subsets were concatenated using `Picard GatherVcfs` into one VCF file covering the entire genome. From the VCF file, SNPs were selected (i.e. indels were excluded) using `GATK SelectVariants`, after which we filtered SNPs with the following criteria: `QD < 2.5; FS > 45.0; SOR > 3.0; MG < 40; MQRankSum < −12.5; ReadPosRankSum < −8.0`. We removed garden warblers and African hill babblers from the multi-species VCF and kept only biallelic sites. We estimated blackcap haplotypes using `SHAPEIT2` (r837) (Delaneau et al., 2013) with the blackcap recombination map (Bascón-Cardozo et al., 2022a), yielding 142,083,056 SNPs.

#### Genome-wide PCA

To characterise the population structure of blackcaps, we performed principal component analysis (PCA) using `PLINK` (Purcell et al., 2007).

#### Local PCA

To identify genomic regions with distinct patterns of genetic variation in blackcaps, we performed local PCA in sliding genomic windows of 1,000 SNPs and summarised dissimilarity of windows by multidimensional scaling using `lostrct` (Li & Ralph, 2019) in R version First, we prepared a genotype and a haplotype table for each chromosome in which rows and columns represented positions and individuals from the phased VCF file using `BCFtools`. Specifically, genotypes were encoded 0, 1, and 2 for the reference allele homozygotes, heterozygotes, and non-reference allele homozygotes in the genotype table, and 0 and 2 for the reference and the non-reference allele in the haplotype table (encoding 0 and 1 instead of 0 and 2 in haplotype-based analysis gives the same results). Chromosomes shorter than 10 Mb were concatenated to avoid misidentification of short chromosomal background as an outlier region. Distance matrices of windows were computed based on the coordinates (PC1 and PC2) of samples (individuals for genotype-based local PCA, and haplotype for haplotype-based local PCA) within R using `lostruct`. Multidimensional scaling (MDS) was performed to summarise similarities of local genetic variation patterns among windows into 20 axes (MDS1 through MDS20).

Using the `lostruct` output, we identified chromosomal intervals with distinct patterns of genetic variation. In each chromosome, windows with MDS value apart from the mode of the distribution by greater than 0.3 for any one of the 20 axes were defined as outlier windows. This threshold was determined by visualising the distribution of MDS values in each chromosome (Sup. Fig. 2). For each MDS axis, we defined genomic intervals with at least five outlier windows longer than 100 kb as “outlier regions” with distinct patterns of genetic variation. Overlapping intervals across different MDS axes as well as intervals identified based on genotypes and haplotypes were merged using `BEDtools`. To verify that the outliers show pattern of genetic variation distinct from the whole-genome PCA, we performed PCA using all SNPs within each outlier region using `PLINK`. Genomic regions showing similar pattern to the whole genome PCA were identified with visual inspection and discarded from the outliers.

To assess consistency between the pipelines using genotypes and haplotypes, we compared MDS results of genotype- and haplotype-based `lostruct`. We calculated Euclidean distance of windows from the centre of the 20 dimensional space to enable comparison of the same window in genotype- and haplotype-based MDS. We measured this distance instead of comparing the coordinates directly to account for possible rotations of MDS patterns between genotype- and haplotype-based `lostruct`. Because dissimilarity of windows in terms of the pattern of genetic variation was computed per chromosome, we calculated correlation of the above distance between genotype- and haplotype-based methods per chromosome. The comparison of genotype-based and haplotype-based `lostruct` is in Sup. Fig. 4.

To assess whether `lostruct` can identify outliers irrespective of presence/absence of other outliers on the same chromosome as well as the chromosome length, we ran `lostruct` treating either one part of a blackcap chromosome (“split chromosomes”) or multiple blackcap chromosomes as a single chromosome (“joined chromosome”). If `lostruct` is robust to the chromosomal background, it is expected that the same regions should be detected as outliers with distinct patterns of genetic variation in both split and joined chromosomes compared to per-chromosome results. We prepared four split chromosomes by splitting chromosomes 1 and 2 at the middle, and one joined chromosome by concatenating chromosomes 20, 21, and 28. We performed `lostruct` analysis based both on genotype and haplotype and merged the identified regions. The comparison of `lostruct` between using single chromosomes and split/joined chromosomes is in Sup. Fig. 5.

#### LD and recombination landscape

To calculate LD around outlier regions, we first extracted SNPs within and 30% length outside each outlier. We then thinned SNPs so that all neighbouring SNP positions were at least 10 kb away from each other. Linkage disequilibrium (LD) between all pairs of thinned SNPs was calculated with `VCFtools` with the `--geno-r2`.

We inferred recombination landscape along blackcap chromosomes using `Pyrho` (Spence & Song, 2019). `Pyrho` infers demography-aware recombination rates with a composite-likelihood approach from SNPs data of unrelated samples making use of likelihood lookup tables generated by simulations based on demography and sample size of each population. In all inferences, we used demography of focal populations inferred in Delmore et al. (2020b). Before the recombination inference, focal samples were filtered and singletons were removed. We ran `Pyrho` with mutation rate of 4.6 × 10*^−^*^9^ per site per generation (Smeds et al., 2016), block penalty of 20, and window size of 50 kb to infer population-level recombination landscape in Azores, Cape Verde, continental resident, and medium-long distance migrants (represented by medium distance south-west migrants). We computed mean recombination rate in 10 kb sliding windows for each population.

We defined low-recombining regions and evaluated overlaps between outlier regions and low-recombining regions in the following four steps. 1. define low-recombining regions for each population recombination map, 2. test for association between all outlier regions and the low-recombining regions for each population, 3. define species-wide and population-specific low-recombining regions, and 4. label outlier regions with species-wide or population-specific low-recombining regions or no overlap with any low-recombining region.

1. For the recombination map of each of the four populations (med_sw, cont_res, Azores, Cape Verde), we defined low-recombining regions as the set of 10 kb windows with recombination rate lower than 20 percentile for each chromosome. This mild threshold was set to account for large variation in the recombination landscapes among chromosomes and to capture population-specific reduction in recombination rate which could be with weaker reduction in recombination rate than at species-wide low-recombining regions.
2. For the set of low-recombining regions of each population, we performed a permutation test by shuffling observed outlier regions within the chromosome and counted the total length of overlap with (any) low-recombining regions (in bp) using `BEDTools`. We repeated this 1,000 times, and compared the empirical null distribution of the overlap length (in bp) with observed overlaps.
3. To define species-wide and population-specific low-recombining regions, at all positions along the genome we counted the number of population recombination maps sharing low-recombining regions. If a region was labelled low-recombining in three or four populations at step 1, we defined it to be species-wide low-recombining region. If a region was labelled low-recombining in one or two populations, we defined it to be a population-specific low-recombining region, recording which populations were low-recombining.
4. We first labelled outlier regions overlapping species-wide low-recombining regions. We intersected the species-wide low-recombining regions defined in step 3 and outlier regions. We labelled an outlier region with species-wide low-recombining if it had a coverage of species-wide low-recombining regions greater than 0.5. Two exceptions were outlier_12_3 and outlier_30_1, which are putative inversions. They had coverage of species-wide low-recombining regions of 0.30 and 0.23 but this is largely due to heterokaryotype-specific recombination suppression and inclusion of homokaryotypes in recombination rate inference. Because these putative inversions were segregated in most populations, we defined them to be species-wide low-recombining regions. We next labelled outlier regions overlapping population-specific low-recombining regions. We intersected the population-specific low-recombining region defined in step 3 with outlier regions excluding those overlapping with species-wide low-recombining regions. We labelled an outlier region overlapping with population-specific low-recombining regions if it had a coverage greater than 0.01 for any (pair of) population(s). Finally, the remaining outlier regions were labelled no reduction in recombination rate.

To characterise genotype-specific LD and recombination landscape at the five outlier regions with three clusters of individuals in PCA, we applied `vcftools --geno-r2` and `Pyrho` (Spence & Song, 2019) to our empirical data using each genotype (AA, AB, and BB in Sup. Fig. 11) separately. Validation of this procedure is described in “Simulation: Validation of LD-based inference of recombination landscape using non-randomly chosen samples”.

#### Inversion breakpoints

Three clusters of individuals observed in PCA with genotype-specific LD at two outlier regions on chromosomes 12 and 30 were indicative of polymorphic inversion (Ma & Amos, 2012; Ruiz-Arenas et al., 2019). To further characterise whether they represent polymorphic inversions, we intended to locate breakpoints by two independent approaches.

##### Soft-clip reads

We attempted to identify positions where presence of soft-clipping of mapped reads is associated with PCA-based genotype of the putative inversions. First, we extracted focal regions around boundaries of the outliers (Sup. Table 4) from read mapping file of all individuals using `SAMtools` (Danecek et al., 2021). Next, we identified soft clip reads in each extracted region using `samextractclip` (Lindenbaum, 2015), and obtained reference position corresponding to the position of soft clipping in mapped reads using a custom script. At all extracted soft-clip positions, we counted the number of reads that switch to soft-clip (“soft-clip depth”), as well as the depth of mapped reads, using `SAMtools`. At each of all positions with at least one read supporting soft-clip switch, we calculated proportion of reads with soft-clip switch relative to all mapped reads (depth of the position) for each individual (“soft-clip proportion”). This resulted in “position-by-individual” matrix whose entry depicts the proportion of soft-clip in all reads mapped at the focal position for the focal individual. Using this matrix, we fit a linear model (soft-clip proportion *∼* PCA − basedgenotype) in `R` at each position treating genotypes AA, AB, and BB as 0, 1, and 2. Based on the significance of genotype and *R*^2^ of the linear models, we generated a list of 14 positions at which soft-clip proportion was significantly associated with genotype of the putative inversions. We visualised the distribution of the soft-clip proportion at these positions (Sup. Fig. 15) and selected six positions for which the soft-clip proportion of BB was high enough and that of AB was around a half of BB based on the assumption that soft clip reads covering an inversion breakpoint should originate from haplotype B and non-soft clip reads should originate from haplotype A (Sup. Table 5). To investigate whether some of these six positions represent inversion breakpoints, we asked whether the soft-clipped segments of the reads have homologous sequences at the other end of the outlier regions. We extracted soft-clipped segments of reads mapped at the focal six positions in AB and BB individuals using a custom script, and re-mapped these segments (instead of the entire reads) to the blackcap reference using `BWA mem`. We computed the depth of mapped segments in each position using `SAMtools` (Sup. Table 5).

##### 10x linked read

We used an independent set of blackcap individuals (hereafter “10x individuals”) whose genomes were sequenced with the 10x linked-read technology (Delmore et al., 2023, NCBI BioProject PRJEB65115). We genotyped the 10x individuals at the two putative inversion loci (i.e. AA, AB, or BB) based on genotypes at diagnostic SNP positions. We started by determining diagnostic SNP positions using our Illumina short read-based resequence data. Because usable diagnostic SNP positions should have genotypes perfectly associated with PCA-based genotype, we focused on positions at which F_ST_ was 1 between AA and BB, and all AB were heterozygous, using `VCFtools` and `BCFtools`. We also recorded mapping between an allele at the diagnostic positions and a genotype of the putative inversion (“A- and B-diagnostic alleles”, e.g. G for haplotype A, T for haplotype B).

We then counted the number of sites with A- and B-diagnostic allele in each of 10x samples. To convert coordinates of 10x assemblies to the reference coordinate, we mapped the 10x pseudo-haplotyped assemblies to the blackcap reference using `minimap2` (Li, 2018).

To determine the putative inversion genotype in the 10x individuals, we counted the number of positions with A-diagnostic and B-diagnostic alleles for each 10x pseudo-haplotype, and calculated the proportion of sites with A-diagnostic and B-diagnostic sites. In principle, an AA and a BB individual respectively are expected to have proportion of 100% and 0% of A-diagnostic sites in both of two pseudo-haplotypes, while an AB individual is expected to have 100% of A-diagnostic sites in one pseudo-haplotype and 0% for the other. For genotyping, we set the following three thresholds.

1. Missingness at the diagnostic positions is less than 10%, after removing positions with non-unique `minimap2` mapping (i.e. at least 90% of all diagnostic positions should have depth of 1x).
2. More than 90% of all diagnostic sites should agree per pseudo-haplotype.
3. The second criterion should be fulfilled for both pseudo-haplotypes of an individual.

We identified two BB individuals for each of the putative inversions on chromosomes 12 and 30. There were no AB individuals passing the above threshold, indicating 10x pseudo-haplotyping is not accurate in separating two diverged non-recombining alleles at a long range in an individual that has both. To identify breakpoints, we aligned the pseudo-haplotype assemblies of these BB individuals as well as one AA individual for each putative inversion to the blackcap reference using `Nucmer4` (Marçais et al., 2018), and generated dot plots (Sup. Fig. 16).

### Sequence analysis at breakpoint of putative inversion on chromosome 12

10x contigs of pseudo-haplotype B aligned next to the putative breakpoint position of blackcap reference chromosome 12 had an un-aligned flanking sequence. To characterise the DNA sequence of these flanking segments, we extracted the flanking sequences using `SAMtools`, aligned the sequences to themselves using `minimap2`, and generated self-dot plots (Sup. Fig. 17), revealing presence of tandem repeats. To identify unit of tandem repeats within the flanking sequences, we ran `TandemRepeatsFinder` against these extracted sequences, resulting in four consensus unit sequences of 144 bp based on two contigs from two individuals. To confirm that the four consensus sequences represent the same tandem repeat (because the unit of identical tandem repeat can have different phases), we ran `BLASTn` (version 2.10.1, Altschul et al., 1990) with each consensus as query against dimers of the consensus. To investigate whether the tandem repeat found at the putative breakpoint of chromosome 12 in haplotype B is present in chromosome 12 and other chromosomes of the reference and corresponding position of haplotype A, we ran `BLASTn` with the 144 bp consensus of the tandem repeat unit as the query against blackcap reference and a contig of an AA individual that spans the breakpoint position, and counted how many copies were found in each reference chromosome/scaffold and the 10x contig (Sup. Fig. 18).

### Selection in blackcaps

To test for selection in different outlier regions and to compare them with the genome-wide base line, we computed nucleotide diversity (*π*) and Tajima’s D in 10 kb sliding windows per population using `PopGenome` (Pfeifer et al., 2014) and `VCFtools` (Danecek et al., 2011) respectively. The effects of the outlier regions on these statistics were tested using a linear mixed effects model (`nlme::lme` (Pinheiro et al., 2021)) and a generalised linear mixed effects model with a Gamma distribution (`lme4::glmer` (Bates et al., 2015)). To test for selection in genes d_N_*/*d_S_ were computed following the counting method by Nei & Gojobori (1986). Gene annotation of the blackcap was obtained from Bascón-Cardozo et al. (2022b).

### Tandem repeats within and outside outlier regions

To characterise correlation between outlier regions with distinct patterns of genetic variation and tandem repeats, we identified tandem repeats in the reference genome and compared the distribution of the tandem repeats with genomic regions with distinct patterns of genetic variation. First, `TandemRepeatsFinder` (Benson, 1999) was run on the blackcap reference genome with the parameter set recommended on the documentation (`trf </path/to/fasta> 2 7 7 80 10 50 500 -f -d -m -h`). The output was formatted and summarised for visualisation using custom scripts. Briefly, distribution of tandem repeats with a different unit size along the genome was summarised in 100 kb sliding windows in blocks of repeat unit sizes of 10 bp step (Sup. Fig. 33). Tandem repeats with the six longest repeat unit size were extracted per chromosome, and copy number for each tandem repeat was counted (Sup. Fig. 34).

Next, we tested whether the number of tandem repeats with long repeat unit were enriched in outlier regions at species-wide and population-specific low-recombining regions. We extracted tandem repeats with repeat unit size greater than or equal to 150 bp, and counted the number of tandem repeats (instead of total copy number) within and outside outlier regions. We performed Fisher’s exact tests to test independence between the number of long tandem repeats and the mode of recombination suppression (species-wide/population-specific) (Sup. Table 7) using `fisher.test` function in R.

## Simulation

### Validation of LD-based inference of recombination landscape using non-randomly chosen samples

We asked whether LD-based recombination map inference using individuals chosen based on the karyotype instead of at random is informative of the underlying mode of recombination suppression. To this end, we simulated two 5 Mb-long chromosomes with neutral mutation rate of 4.6 × 10*^−^*^8^ in a population of 1,000 individuals in `SLiM`. The purpose of these simulations was to investigate the effect of an inversion and additional recombination suppression on recombination rate inference and LD in general, rather than investigating the effects specific to blackcap demography. As such, we kept the population size smaller than the blackcap effective population size and the mutation rate greater than assumed in order to minimise the time and computational resource for simulations. We introduced a mutation (inversion marker) on one chromosome at 1 Mb position at the 50th generation. We simulated an inversion on the chromosome by suppressing recombination in an interval from 1 Mb to 4 Mb position if the inversion marker site was heterozygous. We defined additional suppression according to different scenarios (models 1-6 in Sup. Table 6). We applied negative frequency-dependent selection (fitness of inversion is 1 − (*p_inv_* − 0.2) where *p_inv_* is the frequency of the inversion allele). 1,000 generations after the inversion event, we recorded the mutations in all samples, making a VCF file including all samples. Although 1,000 generations is relatively short given the population size of 1,000, the haplotype structure at the inversion locus was stable in test runs of model-1 (inversion frequency of 0.2 without additional recombination suppression).

Based on the genotype at 1 Mb position, we randomly chose 10 samples for each inversion genotype. `Pyrho` was run to estimate recombination rates using the chosen 10 samples, with the block penalty 50 and window size 50. The inferred recombination maps are in Sup. Fig. 13.

### Effects of recombination suppression model on recombination rate inference at an inversion

Three clusters of individuals observed in PCA at five outlier regions indicate presence of distinct haplotypes. Polymorphic inversions are known to show this pattern due to suppression of recombination between the normal and inverted alleles (Wellenreuther & Bernatchez, 2018). To test whether some of the five outlier regions represent polymorphic inversions, we intended to infer recombination rates using AA, AB, and BB individuals separately based on linkage disequilibrium (LD) patterns. Before addressing this in blackcaps empirically, we assessed how different types of recombination suppression at a haplotype block affect inference of recombination landscape using a set of individuals with a certain combination of haplotypes. To investigate the effect of a genotype-specific suppression of recombination on LD-based inference of recombination rate, we simulated different modes of recombination suppression using `SLiM` version 3.5 (Haller & Messer, 2019) under six scenarios listed in Sup. Table 6. Specifically, we performed 1,000 replicates of forward-time simulations of two 500 kb-long chromosomes with neutral mutation rate of 1×10*^−^*^7^ [per site per generation] and recombination rate of 1 × 10*^−^*^6^ [per site per generation] in a population of 1,000 diploid individuals under the Wright-Fisher model (We downscaled the population size and upscaled mutation rate to minimise the time and computational resource for simulation). We introduced a mutation (inversion marker) on one chromosome at 100 kb position at the 50th generation. We modelled an inversion by suppressing recombination in an interval from 100 kb to 400 kb position if the inversion marker site was heterozygous. We defined additional suppression according to different scenarios (models 1-6). To allow for the inversion to remain in the population, we applied negative frequency-dependent selection (fitness of inversion is 1 − (*p_inv_* − 0.2) for models 1-3 and 1 − (*p_inv_* − 0.8) for models 4-6 where *p_inv_* is the frequency of the inversion allele). 1,000 generations after the inversion event, we recorded the mutations in all samples, making a VCF file including all individuals. Although 1,000 generations is relatively short given the population size of 1,000, the haplotype structure at the inversion locus was stable in test runs of model-1 (inversion frequency of 0.2 without additional recombination suppression). Based on the genotype at the marker, we randomly sampled 10 individuals for each inversion genotype. `Pyrho` was run to estimate recombination rates using the sampled 10 individuals, with the block penalty 50 and window size 50. The inferred recombination landscape is in Sup. Fig. 13.

### Coalescent simulation of species-wide reduction of recombination rate

To discern the effect of reduced recombination rate, demographic history, and unequal sample sizes among population on outlier regions identified by `lostruct`, we performed neutral coalescent simulations using msprime version 1.2.0 (Baumdicker et al., 2022). We simulated a 1-Mb long recombining chromosome with a mutation rate of 4.6 × 10*^−^*^9^ [per site per generation]. We implemented 11 models differing in the recombination maps, population subdivision, and demographic history (Sup. Fig. 19). In models 1-3, the recombination rate was set to 4.6×10*^−^*^9^ [per site per generation] throughout the entire chromosome, and they differ in population subdivision (model 1: panmictic, model 2, subdivision of five equal populations without gene flow, model 3: subdivision of equally-sized populations with gene flow between two pairs of populations (symmetric migration rate of 0.025 [per generation])). In models with five populations, we distributed the sample of 100 individuals unequally, as in our blackcap dataset (50, 20, 10, 10, 10 individuals for five populations). In models 4-7, we introduced reduced recombination rate in the middle of the chromosome (0.4 to 0.6 Mb) with the same demographic histories as models 2 and 3. In addition to the uniform recombination map, we prepared two recombination maps with reduced recombination rate: “low-rec” with one-hundreth the background recombination rate, and “no-rec” with recombination rate of 0. In models 8-11, we used the same two recombination maps with reduced recombination rate in the middle, with different demography: 10 times increase in effective population size in one population, and 10 times descrease in effective population size in three populations, which roughly reflects inferred demography of blackcap populations (Delmore et al., 2020b). For each model, we ran 1,000 replicates of simulations and recorded SNPs in VCF format.

To identify outlier regions, we ran `lostruct` the same way as in the empirical analysis. To evaluate how reduced recombination rate affects the mean and variance of population genetic summary statistics, we computed nucleotide diversity (*π*), Tajima’s D, and F_ST_, using `VCFTools`. The outliers detected by `lostruct` are in Sup. Fig. 20. The summary statistics are in Sup. Figs. 21, 22, 23.

### Forward simulation of species-wide reduction of recombination rate

To investigate how species-wide low-recombining regions affect patterns of local genetic variation depicted in local PCA, we performed forward simulation with `SLiM` version 4.0.1 (Haller & Messer, 2022). We simulated 100 replicates of two 500 kb-long chromosomes with neutral mutation rate of 1 × 10*^−^*^7^ [per site per generation] and recombination rate of 1 × 10*^−^*^6^ [per site per generation] except for an interval from 100 to 400 [kb] of the first chromosome where recombination rate was set to 1 × 10*^−^*^9^, which is 1/1000 of the normally recombining chromosome. First, we ran a burn-in of 4,000 generations for an ancestral population of 1,000 diploids. After the burn-in, we made three populations of 1,000 diploids (pop1, pop2, and pop3) split from the ancestral population. We sampled 50 individuals per population every 20 generations over 1,000 generations after the population split and recorded SNPs in VCF. For each time point of each of 100 simulation replicates, we performed PCA with `PLINK`, using SNPs either within 100 to 400 [kb] of the first chromosome (pop1-specific suppression) or the normally recombining chromosome.

We investigated how reduced recombination rate affects representation of population subdivision in local PCA. To evaluate whether the individuals from different populations were distributed differently in local PCA at the low-recombining region, we performed Fasano-Franceschini test (Fasano & Franceschini, 1987), which is a multi-dimensional extension of Kolmogorov-Smirnov test, in three pairs of populations (pop1-pop2, pop1-pop3, pop2-pop3). We counted the number of significant pairs of populations (0, 1, 2, or 3) for each time point of each replicate. We compared between the low-recombining and normally recombining regions the number of pairs of populations with distinct distribution in PCA (Sup. Fig. 31).

### Forward simulation of population-specific reduction of recombination rate

To investigate how evolution of low-recombining regions in population(s) affect patterns of local genetic variation depicted in local PCA, we performed forward simulation with `SLiM` version 4.0.1. We simulated two 500kb-long chromosomes with neutral mutation rate and recombination rate of 1 × 10*^−^*^7^ [per site per generation] and 1 × 10*^−^*^6^ [per site per generation]. First, we ran a burn-in of 4,000 generations for an ancestral population of 1,000 diploids. After the burn-in, we made three populations of 1,000 diploids (pop1, pop2, and pop3) split from the ancestral population, after which gene flow between all pairs of populations were set to 0.0025. We introduced recombination suppression in pop1 from 100 to 400 [kb] of the first chromosome in two scenarios. In the first scenario, recombination suppression was introduced at the same time of the split. In the second scenario, recombination suppression was introduced 4,000 generations after the population split event, allowing the three populations to differentiate before population-specific recombination suppression was introduced in pop1. We sampled 50 individuals per population every 20 generations over 1,000 generations after the introduction of the population-specific suppression of recombination and recorded SNPs in VCF. For each time point of each of 1,000 simulation replicates, we performed PCA with `PLINK`, using SNPs either within 100 to 400 [kb] of the first chromosome (pop1-specific suppression) or the normally recombining chromosome.

To characterise factors represented in the primary axes of distinct local PCA at population-specific low-recombining regions, we performed one replicate of `SLiM` simulation with the same scenarios of models 1 and 2 recording the full ancestry and mutations in tree sequence, with an increased duration of burn-in (40,000 generations) to make sure that all lineages at sampling time coalesce. We loaded the tree sequence with mutations in `tskit` (Kelleher et al., 2018) and sampled 50 diploids per population, and saved SNPs in VCF. Using the VCF files for each time point for each model, we performed PCA using `PLINK` at the population-specific low-recombining region, and determined one time point per model showing typical spread of individuals from the low-recombining population in PCA (Sup. Fig. 27A, E). For these PCAs we identified 5% SNPs with the highest loadings to the first two PC axes. We analysed these mutations on the underlying genealogies using `tskit`. Specifically, we investigated whether mutations originating from the low-recombining population were enriched in the high-loading mutations (Sup. Fig. 27C, G) with a *χ*^2^ test. We also assessed whether multiple mutations originating in the low-recombining population occurring on the same genealogical branches (i.e. mutations on the same ancestral haplotypes) were enriched in the high-loading mutations (Sup. Fig. 27D, H). For this, we compared the number of mutations sharing the same genealogical branches among the high-loading mutations originating from the low-recombining population and the same number of randomly-selected mutations originating from the low-recombining population by a Kolmogorov-Smirnov test.

### Effects of linked selection on local PCA

#### Background selection

To investigate the linked effect of purifying selection at low-recombining regions (background selection) on patterns of local genetic variation represented in local PCA, we performed forward simulation with `SLiM` version 4.0.1. We simulated a species-wide low-recombining region in three populations as described above, except we changed the distribution of fitness effect of mutations with three different ratios between neutral (“n”, *s* = 0) and deleterious (“d”, *s* = −0.05 and *h* = 0.5) mutations of *n/*(*n* + *d*) = 0, 0.25, 0.5, 0.75. To evaluate whether individuals from different populations were distributed differently in the local PCA at the low-recombining region, we performed Fasano-Franceschini test between three pairs of populations (pop1-pop2, pop1-pop3, pop2-pop3). We counted the number of significant pairs of populations (0, 1, 2, or 3) for each sampled time point of each replicate (out of 100) for each DFE (Sup. Fig. 31).

#### Positive selection

To investigate the linked effect of positive selection at low-recombining regions on patterns of local genetic variation represented in local PCA, we performed forward simulation with `SLiM` version 4.0.1 under four scenarios: population-specific sweep and sweep before populations split, with and without reduced local recombination rate. We simulated 10 replicates of one 500 kb-long chromosome with neutral mutation rate of 1 × 10*^−^*^7^ [per site per generation] and recombination rate of 1 × 10*^−^*^6^ [per site per generation]. In scenarios with reduced recombination rate, we introduced a reduced recombination rate within an interval from 100 to 400 [kb] of the chromosome where recombination rate was set to 1 × 10*^−^*^9^, which is 1/1000 of the normally recombining regions. For all scenarios, we ran a burn-in of 4,000 generations for an ancestral population of 1,000 diploids. In the scenarios with population-specific sweep, we made three populations of 1,000 diploids (pop1, pop2, and pop3) split from the ancestral population at the 4000-th generation. We introduced a strongly beneficial mutation (*s* = 1 and *h* = 0.5) in the middle of a chromosome of one randomly selected sample of the first population at the 100-th generation after the populations split. In the scenarios with sweep before split, we introduced a strongly beneficial mutation (*s* = 1 and *h* = 0.5) in the middle of the chromosome of one randomly selected sample of the ancestral population, and made the three populations of 1,000 diploids split at the 100-th generation after the introduction of the beneficial mutation. We sampled 100 diploid individuals per population every 20 generations since the introduction of the beneficial mutation (scenarios of population-specific sweep) or the split (scenarios of ancestral sweep) and recorded the SNPs in VCF format. We performed PCA using `PLINK`.

## Acknowledgment

This work was supported by the Max Planck Society (Max Planck Research Group grant MFFALIMN0001 to ML), the DFG (project Z02 and Nav05 within SFB 1372 – Magnetoreception and Navigation in Vertebrates to ML), and DFG Research Infrastructure NGS_CC (project 407495230) as part of the Next Generation Sequencing Competence Network (project 423957469). AR was supported by the Intramural Research Program of the NHGRI, NIH (1ZIAHG200398). JCI was funded by two research grants from the Spanish Ministry of Science, Innovation and Universities, and the European Regional Development Fund (PGC2018-097575-B-I00; PID2022-140091NB-I00). We thank Britta Meyer, Tianhao Zhao, Hanna Koch, Conny Burghardt, and Sven Künzel for DNA extraction, library preparation, and/or sequencing. We are grateful to Julien Dutheil, Diethard Tautz, Linda Odenthal-Hesse, Tobias Kaiser, Carolina Peralta, and Matthias Weissensteiner for constructive discussion. We are grateful to Thord Fransson, Christos Barboutis, Zura Javakhishvili, Martim Melo, Álvaro Ramírez, and Helena Batalha for providing us with samples. Permits were provided to JCI for samples collected on Cape Verde (Ministerio do Ambiente - Habitacao e Ordenamento do Territorio, 18/CITES/DNA, 17 Dec 2015), Canary Islands (Ref.: 2012/0710), Madeira (Ref.: 02/2016), and the Azores (Instituto da Conservacao da Natureza e da Biodiversidade, 171/2008, 31 Mar 2009); to JP-T for samples collected on Mallorca (CAP 64/2009); to Thord Fransson for samples collected on Crete (6Υ0Ξ4653Π8-ΥΓ 5 issued by the Hellenic Ministry of Environment and Energy), and to Zura Javakhishvili for samples collected in Georgia (889-0-2-202303291450 by the Ministry of Environment and Agriculture of Georgia). A preprint version of this article has been peer-reviewed and recommended by PCIEvolBiol (https://doi.org/10.24072/pci.evolbiol.100711).

## Data availability

The primary and alternate haplotype assemblies of the blackcap reference genome can be found under NCBI BioProject PRJNA558064 (accenssion GCA_009819655.1) and PRJNA558065 (accession GCA_009819715.1). Raw Illumina reads for the resequencing data can be accessed under NCBI BioProject PRJEB66075 (SRA accession ERP151147). Processed data and scripts for analysis and simulation are found in Zenodo (https://doi.org/10.5281/zenodo.10623362).

## Conflict of interest

The authors declare no conflict of interest.

## Author contributions

JI and ML designed the study. Reference genome was generated by JF, AR, JM, BH, WC, JC, KH, MU, OF, and EDJ. JP-T and JCI collected samples for resequencing. AB performed read mapping, variant calling, and data filtration. KB-C inferred recombination maps. JI conducted haplotype inference, population genomics analyses, simulations, sequence analyses, statistical modelling, and data visualisation. JI and ML wrote the manuscript with inputs from other authors.

## Notes

### Competing Interest Statement

The authors have declared no competing interest.

### Summary of Updates

In Results, exact p-values have been included. The number of permutations has been specified. In Figure 6 legend, two references have been added. A PCI badge has been added to the front page. Acknowledgment to PCI EvolBiol has been added.

