## Supplementary information for "Distinct patterns of genetic variation at low-recombining genomic regions represent haplotype structure"

**4 References** **42**

### 1 Supplementary Notes

#### 1.1 Discussion: Contribution of genealogical and mutational noise on local genetic variation

The amount of genealogical and mutational noise on realised genetic variation is influenced by the number of genealogies, the length of the genomic interval, and the length of branches. First, the effect of genealogical noise on realised genetic variation reduces as the number of genealogies increases (i.e. as the local recombination rate increases). If a genomic interval contains only one genealogy with enough mutations, then the realised genetic variation tends to reflect this particular tree (i.e. without removal of the genealogical noise) instead of a set of trees with different probabilities expected under the population history. On the contrary, if a genomic interval contains many genealogies each with enough mutations (i.e. with removal of genealogical noise), then the realised genetic variation tends to reflect the expectation under the population history. Second, the effect of mutational noise on realised genetic variation reduces as the product of branch length and the genomic interval increases. This is because the product represents total amount of substrate for mutations to occur in one ancestral haplotype (Shipilina et al., 2023). When this product is small, length of genealogical branches may be disproportionately represented by the number of mutations, increasing the relative effect of mutational noise.

#### 1.2 Discussion: Effects of selection on distinct genetic variation

Using neutral simulations and comparing them with simulations with selection, we showed that primary axes of PCA represent effect of genealogical noise at low-recombining regions, instead of effect of selection (distorted genealogies). The prominent effect of reduced recombination rate over selection may be partly due to the scenarios tested in our simulations. For example, we tested the effects of positive selection under the hard sweep model, in which selection acted on a novel mutation in a single copy, resulting in fixation of a single haplotype. Other scenarios of selection causing multiple haplotypes, such as selection on standing genetic variation (Stephan, 2019), may result in greater contribution by selection on haplotype structure than by reduced recombination rate.

#### 1.3 Results and discussion: Spread of individuals of low-recombining populations in local PCA

We note that the observed patterns of genetic variation at the population-specific low-recombining regions in the blackcap are also qualitatively similar to transient effect of population-specific selection (spread of individuals of population without selection (Sup. Fig. 32B)). However, the selection scenario cannot explain the empirical observations of moderate levels of nucleotide diversity (Sup. Fig. 29) and Tajima's D values around the genome-wide average (Sup. Fig. 28), and is hence unlikely.

### 2 Supplementary Tables

**Supplementary Table 1:** Samples used in this study. The spreadsheet can be downloaded from bioRxiv.

**Supplementary Table 2:** Zebra finch chromosomes (GCA_003957565.4) and corresponding blackcap chromosomes in our *de novo*-assembled genome. Note that Sylvioidea, to which the blackcap belongs, has neo-sex chromosome segments (in Z and W) originating from a part of chromosome 4A (zebra finch, homologous to blackcap chromosome 21) (Leroy et al., 2021; Pala et al., 2012; Sigeman et al., 2020, 2021).

| Chr. (Zebra finch) | Length [bp] | Chr. (Blackcap) | Length [bp] |
|---|---|---|---|
| 1 | 114,020,016 | 2 | 115,287,908 |
| 1A | 71,569,005 | 5 | 72,605,035 |
| 2 | 151,653,088 | 1 | 153,194,430 |
| 3 | 112,598,992 | 3 | 113,359,798 |
| 4 | 70,982,421 | 4 | 72,978,054 |
| 4A | 19,491,698 | 21 | 10,058,197 |
| 5 | 61,663,524 | 6 | 63,423,007 |
| 6 | 35,592,820 | 8 | 35,969,369 |
| 7 | 37,979,234 | 7 | 38,862,568 |
| 8 | 31,251,066 | 9 | 31,650,914 |
| 9 | 25,529,500 | 10 | 26,389,481 |
| 10 | 20,543,327 | 13 | 20,608,123 |
| 11 | 21,108,237 | 12 | 22,249,080 |
| 12 | 20,384,687 | 11 | 22,342,531 |
| 13 | 18,052,104 | 14 | 19,084,229 |
| 14 | 16,340,488 | 15 | 16,052,774 |
| 15 | 13,771,869 | 17 | 14,194,216 |
| 16 | 1,095,113 | 17 | 14,194,216 |
| 17 | 11,257,394 | 19 | 11,408,529 |
| 18 | 11,152,170 | 18 | 11,994,622 |
| 19 | 8,335,140 | 20 | 11,247,436 |
| 20 | 14,749,488 | 16 | 15,230,834 |
| 21 | 7,924,511 | 22 | 7,560,935 |
| 22 | 4,608,043 | 28 | 4,652,909 |
| 23 | 6,804,446 | 24 | 6,883,216 |
| 24 | 7,620,594 | 23 | 7,468,107 |
| 25 | 2,880,192 | 30 | 2,016,562 |
| 26 | 6,804,938 | 25 | 6,775,589 |
| 27 | 5,378,928 | 27 | 5,004,062 |
| 28 | 5,866,305 | 26 | 5,062,550 |
| 29 | 2,636,872 | 29 | 2,201,499 |
| W | 20,846,394 | W | 21,688,951 |
| Z | 75,396,176 | Z | 88,576,696 |

**Supplementary Table 3:** Genomic regions with distinct pattern of genetic variation ("outliers" based on local PCA using `lostruct`).

| Outlier ID | Chromosome | From [bp] | To [bp] | Length [bp] | Red. rec. |
|---|---|---|---|---|---|
| outlier_1_1 | 1 | 56,264,814 | 57,679,298 | 1,414,485 | Spp-wide |
| outlier_1_2 | 1 | 76,931,639 | 77,195,852 | 264,214 | Spp-wide |
| outlier_2_1 | 2 | 17,876,150 | 18,183,386 | 307,237 | Spp-wide |
| outlier_2_2 | 2 | 18,301,736 | 19,824,380 | 1,522,645 | Spp-wide |
| outlier_2_3 | 2 | 114,618,118 | 114,874,768 | 256,651 | CapeVerde |
| outlier_3_1 | 3 | 1,317,747 | 1,464,754 | 147,008 | No reduction |
| outlier_3_2 | 3 | 108,194,892 | 108,406,259 | 211,368 | Azores |
| outlier_4_1 | 4 | 11,647,463 | 11,966,907 | 319,445 | med_sw |
| outlier_5_1 | 5 | 68,065,279 | 68,294,444 | 229,166 | Spp-wide |
| outlier_6_1 | 6 | 5,687,267 | 6,323,968 | 636,702 | Spp-wide |
| outlier_6_2 | 6 | 62,340,071 | 62,456,329 | 116,259 | No reduction |
| outlier_8_1 | 8 | 30,282,659 | 30,636,429 | 353,771 | Spp-wide |
| outlier_10_1 | 10 | 11,602,450 | 13,202,422 | 1,599,973 | Spp-wide |
| outlier_12_1 | 12 | 61 | 208,254 | 208,194 | Spp-wide |
| outlier_12_2 | 12 | 1,850,450 | 2,310,703 | 460,254 | CapeVerde |
| outlier_12_3 | 12 | 14,118,030 | 22,229,395 | 8,111,366 | Spp-wide |
| outlier_14_1 | 14 | 43 | 207,189 | 207,147 | Spp-wide |
| outlier_14_2 | 14 | 14,789,430 | 15,024,671 | 235,242 | Azores;CapeVerde |
| outlier_14_3 | 14 | 15,956,522 | 16,206,154 | 249,633 | Azores;CapeVerde |
| outlier_15_1 | 15 | 13,458,327 | 14,146,280 | 687,954 | Azores;CapeVerde |
| outlier_15_2 | 15 | 15,846,846 | 16,049,116 | 202,271 | Spp-wide |
| outlier_16_1 | 16 | 1,510,566 | 1,842,318 | 331,753 | Spp-wide |
| outlier_17_1 | 17 | 13,936,617 | 14,179,092 | 242,476 | Spp-wide |
| outlier_20_1 | 20 | 25,274 | 338,180 | 312,907 | Spp-wide |
| outlier_20_2 | 20 | 1,499,745 | 1,627,522 | 127,778 | Azores;CapeVerde |
| outlier_21_1 | 21 | 611,083 | 757,333 | 146,251 | Azores |
| outlier_21_2 | 21 | 3,003,124 | 3,440,115 | 436,992 | Azores;CapeVerde |
| outlier_28_1 | 28 | 939,876 | 1,143,638 | 203,763 | Azores;CapeVerde |
| outlier_30_1 | 30 | 72 | 1,471,845 | 1,471,774 | Spp-wide |
| outlier_Z_1 | Z | 14,585,807 | 14,978,909 | 393,103 | Spp-wide |
| outlier_Z_2 | Z | 23,428,533 | 24,628,650 | 1,200,118 | Spp-wide |
| outlier_Z_3 | Z | 47,230,133 | 47,412,498 | 182,366 | Spp-wide |

**Supplementary Table 4:** Genomic regions within which soft-clipped reads were analysed to identify breakpoints of putative inversions at outlier_12_3 and outlier_30_1.

| Chromosome | From | To |
|---|---|---|
| 12 | 14,080,000 | 14,250,000 |
| 12 | 20,500,000 | 22,249,080 |
| 30 | 0 | 160,000 |
| 30 | 1,340,000 | 1,475,000 |

**Supplementary Table 5:** Soft-clipped segments remapped to the blackcap reference. The first two columns are filtered positions (detailed in Materials and Methods and Sup. Fig. 15) of putative breakpoints based on soft clip reads associated with PCA-based genotype at putative inversions at outlier_12_3 and outiler_30_1. The third through the sixth columns show summary of remapped soft-clipped segments. The absence of mapping to the other end of the outlier region indicates that our analysis based on soft-clip proportion of Illumina short reads failed to locate breakpoints of the putative inversions.

| Chromosome | Position [bp] | Chromosome | From [bp] | To [bp] | Mean depth |
|---:|---:|---:|---:|---:|---:|
| 12 | 14,174,953 | 13 | 20,601,341 | 20,601,412 | 66.96 |
| 12 | 14,174,953 | 3 | 113,350,210 | 113,350,264 | 96.27 |
| 12 | 14,174,961 | 5 | 68,337,734 | 68,337,769 | 30.61 |
| 12 | 14,174,961 | 5 | 68,337,771 | 68,337,824 | 79.78 |
| 12 | 21,703,198 | 12 | 21,703,358 | 21,703,473 | 108.40 |
| 12 | 21,800,952 | scaffold_096 | 14,484 | 14,525 | 17.31 |
| 12 | 21,800,952 | scaffold_096 | 21,919 | 21,967 | 26.78 |
| 12 | 21,800,952 | scaffold_133 | 19,291 | 19,372 | 111.66 |
| 12 | 22,046,831 | 1 | 141,582,638 | 141,582,679 | 29.71 |
| 12 | 22,046,831 | 2 | 85,518,469 | 85,518,525 | 22.98 |
| 12 | 22,046,831 | W | 3,399,537 | 3,399,588 | 58.15 |
| 12 | 22,046,831 | W | 3,765,336 | 3,765,394 | 22.64 |
| 12 | 22,046,831 | W | 19,239,977 | 19,240,059 | 50.27 |
| 30 | 142,877 | 30 | 143,588 | 143,705 | 217.43 |

**Supplementary Table 6:** Six models of recombination suppression at a haplotype block simulated with SLiM. We investigated whether inference of genotype-specific recombination maps can capture heterozygote-specific suppression of recombination at polymorphic inversions by performing simulation under the six models of recombination suppression. N: normal (non-inversion) allele; I: inversion allele. The results are shown in Sup. Fig. 13.

| | Model 1 | Model 2 | Model 3 | Model 4 | Model 5 | Model 6 |
|---|---|---|---|---|---|---|
| Derived haplotype frequency | 0.2 | 0.2 | 0.2 | 0.8 | 0.8 | 0.8 |
| Recombination suppression N-I | Yes | Yes | Yes | Yes | Yes | Yes |
| Recombination suppression I-I | No | Yes | Yes | No | Yes | Yes |
| Recombination suppression N-N | No | No | Yes | No | No | Yes |

**Supplementary Table 7:** Result of Fisher's exact tests for the number of tandem repeats with unit sizes longer than or equal to 150 bp within and outside of outlier regions. Genomic distribution of tandem repeats is visualised in Sup. Fig. 33.

| class | n. long TRs in | n. long TRs out | total length [bp] | 95% CI of odds ratio | p-value |
|---:|---:|---:|---:|:---:|:---:|
| spp-wide | 10 | 62 | 19,191,668 | 3.9-16.8 | <0.001 |
| pop-specific | 0 | 72 | 3,335,331 | 0.0-16.3 | 1.000 |

**Supplementary Table 8:** 11 models of demography and recombination maps simulated with msprime.

| Scenario | Relative rec. rate | Pop. subdivision | Unequal pop. size | Gene flow |
|---|---|---|---|---|
| 1 | 1.00 | No | - | - |
| 2 | 1.00 | Yes | No | No |
| 3 | 1.00 | Yes | No | Yes |
| 4 | 0.01 | Yes | No | No |
| 5 | 0.00 | Yes | No | No |
| 6 | 0.01 | Yes | No | Yes |
| 7 | 0.00 | Yes | No | Yes |
| 8 | 0.01 | Yes | Yes | No |
| 9 | 0.01 | Yes | Yes | Yes |
| 10 | 0.00 | Yes | Yes | No |
| 11 | 0.00 | Yes | Yes | Yes |

**Supplementary Table 9:** Linear mixed-effects model testing the fixed effects of outlier regions on Tajima's D, treating chromosome, population, and species-wide/population-specific low-recombining region as random effects variables.

| | Estimate | SE | t-value | p-value |
|---|---|---|---|---|
| Intercept | -0.2196099 | 0.12894037 | -1.70319 | 0.0885 |
| outlier_1_1 | -0.7610205 | 0.01809935 | -42.04684 | 0.0000 |
| outlier_1_2 | -0.2288175 | 0.03922094 | -5.83406 | 0.0000 |
| outlier_2_1 | -0.0892122 | 0.03723778 | -2.39574 | 0.0166 |
| outlier_2_2 | -0.7153382 | 0.01943918 | -36.79879 | 0.0000 |
| outlier_2_3 | 0.3177984 | 0.01801459 | 17.64116 | 0.0000 |
| outlier_3_1 | 0.2517975 | 0.02484491 | 10.13477 | 0.0000 |
| outlier_3_2 | -0.0288526 | 0.02100110 | -1.37386 | 0.1695 |
| outlier_4_1 | -0.3321943 | 0.01922048 | -17.28335 | 0.0000 |
| outlier_5_1 | -0.6917180 | 0.04515554 | -15.31856 | 0.0000 |
| outlier_6_1 | -1.2596634 | 0.02867480 | -43.92929 | 0.0000 |
| outlier_6_2 | 0.3531770 | 0.02928475 | 12.06010 | 0.0000 |
| outlier_8_1 | -1.4311088 | 0.03868169 | -36.99706 | 0.0000 |
| outlier_10_1 | -0.1856898 | 0.01995984 | -9.30317 | 0.0000 |
| outlier_12_1 | 0.0202982 | 0.04782549 | 0.42442 | 0.6713 |
| outlier_12_2 | 0.3645219 | 0.01734986 | 21.01008 | 0.0000 |
| outlier_12_3 | 0.1153698 | 0.00717804 | 16.07261 | 0.0000 |
| outlier_14_1 | 0.0395049 | 0.04697090 | 0.84105 | 0.4003 |
| outlier_14_2 | 0.0401297 | 0.01945566 | 2.06263 | 0.0391 |
| outlier_14_3 | 0.2814038 | 0.02327442 | 12.09069 | 0.0000 |
| outlier_15_1 | 0.2511851 | 0.01110711 | 22.61479 | 0.0000 |
| outlier_15_2 | -0.5317589 | 0.04838627 | -10.98987 | 0.0000 |
| outlier_16_1 | -0.3228744 | 0.03712518 | -8.69691 | 0.0000 |
| outlier_17_1 | 0.1660338 | 0.04238137 | 3.91761 | 0.0001 |
| outlier_20_1 | -1.2035238 | 0.04124938 | -29.17678 | 0.0000 |
| outlier_20_2 | -0.0907998 | 0.02876938 | -3.15613 | 0.0016 |
| outlier_21_1 | 0.1726014 | 0.02510559 | 6.87502 | 0.0000 |
| outlier_21_2 | 0.1919318 | 0.01356198 | 14.15220 | 0.0000 |

| | Estimate | SE | t-value | p-value |
|---|---|---|---|---|
| outlier_28_1 | -0.0454387 | 0.02369477 | -1.91767 | 0.0552 |
| outlier_30_1 | 0.1352691 | 0.01609042 | 8.40681 | 0.0000 |
| outlier_Z_1 | 0.6877002 | 0.05775299 | 11.90761 | 0.0000 |
| outlier_Z_2 | -0.6255465 | 0.02026285 | -30.87160 | 0.0000 |
| outlier_Z_3 | -0.5121660 | 0.05049917 | -10.14207 | 0.0000 |

**Supplementary Table 10:** Generalised linear mixed-effects model (GLMM) with Gamma distribution with log-link function testing the fixed effects of outlier regions on $\pi$, treating chromosome, population, and species-wide/population-specific low-recombining region as random effects variables.

| | Estimate | SE | t-value | p-value |
|---|---|---|---|---|
| Intercept | -5.683314 | 0.054636 | -104.022 | <2e-16 |
| outlier_1_1 | -1.356542 | 0.014813 | -91.576 | <2e-16 |
| outlier_1_2 | -0.815256 | 0.032498 | -25.087 | <2e-16 |
| outlier_2_1 | -1.149724 | 0.029935 | -38.408 | <2e-16 |
| outlier_2_2 | -1.973864 | 0.015217 | -129.712 | <2e-16 |
| outlier_2_3 | 0.314815 | 0.031690 | 9.934 | <2e-16 |
| outlier_3_1 | -1.068258 | 0.046546 | -22.951 | <2e-16 |
| outlier_3_2 | 0.440471 | 0.035297 | 12.479 | <2e-16 |
| outlier_4_1 | -0.615527 | 0.030914 | -19.911 | <2e-16 |
| outlier_5_1 | -1.426243 | 0.032550 | -43.817 | <2e-16 |
| outlier_6_1 | -2.233589 | 0.021155 | -105.584 | <2e-16 |
| outlier_6_2 | 0.156340 | 0.046986 | 3.327 | 0.000877 |
| outlier_8_1 | -2.110366 | 0.027630 | -76.380 | <2e-16 |
| outlier_10_1 | -1.681078 | 0.016472 | -102.056 | <2e-16 |
| outlier_12_1 | -1.059585 | 0.035687 | -29.691 | <2e-16 |
| outlier_12_2 | -2.917098 | 0.027999 | -104.185 | <2e-16 |
| outlier_13_3 | -0.006737 | 0.008072 | -0.835 | 0.403969 |
| outlier_14_1 | -0.973331 | 0.034501 | -28.212 | <2e-16 |
| outlier_14_2 | 0.186851 | 0.033156 | 5.636 | 1.75e-08 |
| outlier_14_3 | -2.485696 | 0.038057 | -65.315 | <2e-16 |
| outlier_15_1 | 0.377986 | 0.020905 | 18.081 | <2e-16 |
| outlier_15_2 | -1.212848 | 0.034497 | -35.158 | <2e-16 |
| outlier_16_1 | -1.482747 | 0.028237 | -52.510 | <2e-16 |
| outlier_17_1 | -0.800511 | 0.032404 | -24.704 | <2e-16 |
| outlier_20_1 | -2.039013 | 0.029654 | -68.761 | <2e-16 |
| outlier_20_2 | -0.268680 | 0.042148 | -6.375 | 1.83e-10 |
| outlier_21_1 | 0.002279 | 0.040307 | 0.057 | 0.954921 |
| outlier_21_2 | 0.211965 | 0.025634 | 8.269 | <2e-16 |
| outlier_28_1 | -0.661910 | 0.035531 | -18.629 | <2e-16 |
| outlier_30_1 | 0.487019 | 0.025900 | 18.804 | <2e-16 |
| outlier_Z_1 | 0.809887 | 0.039414 | 20.548 | <2e-16 |
| outlier_Z_2 | -0.886677 | 0.015829 | -56.017 | <2e-16 |
| outlier_Z_3 | -0.666644 | 0.036808 | -18.111 | <2e-16 |

### 3 Supplementary Figures

#### 3.1 Whole-genome PCA

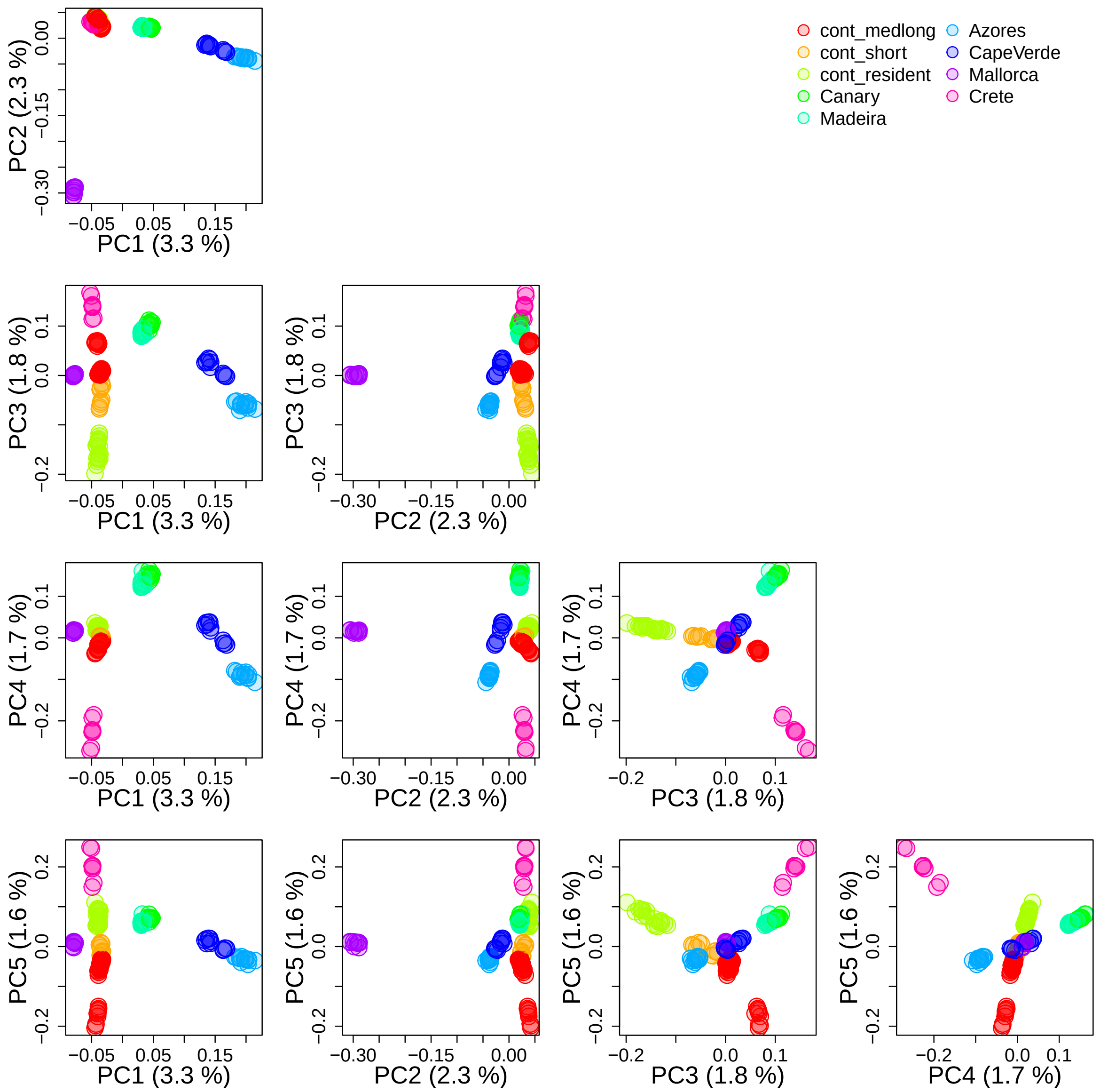

**Supplementary Figure 1: Population structure analysed with whole-genome PCA.** Supplementary data related to Fig. 2B, C. The first five PCs are shown.

#### 3.2 Local PCA and recombination map

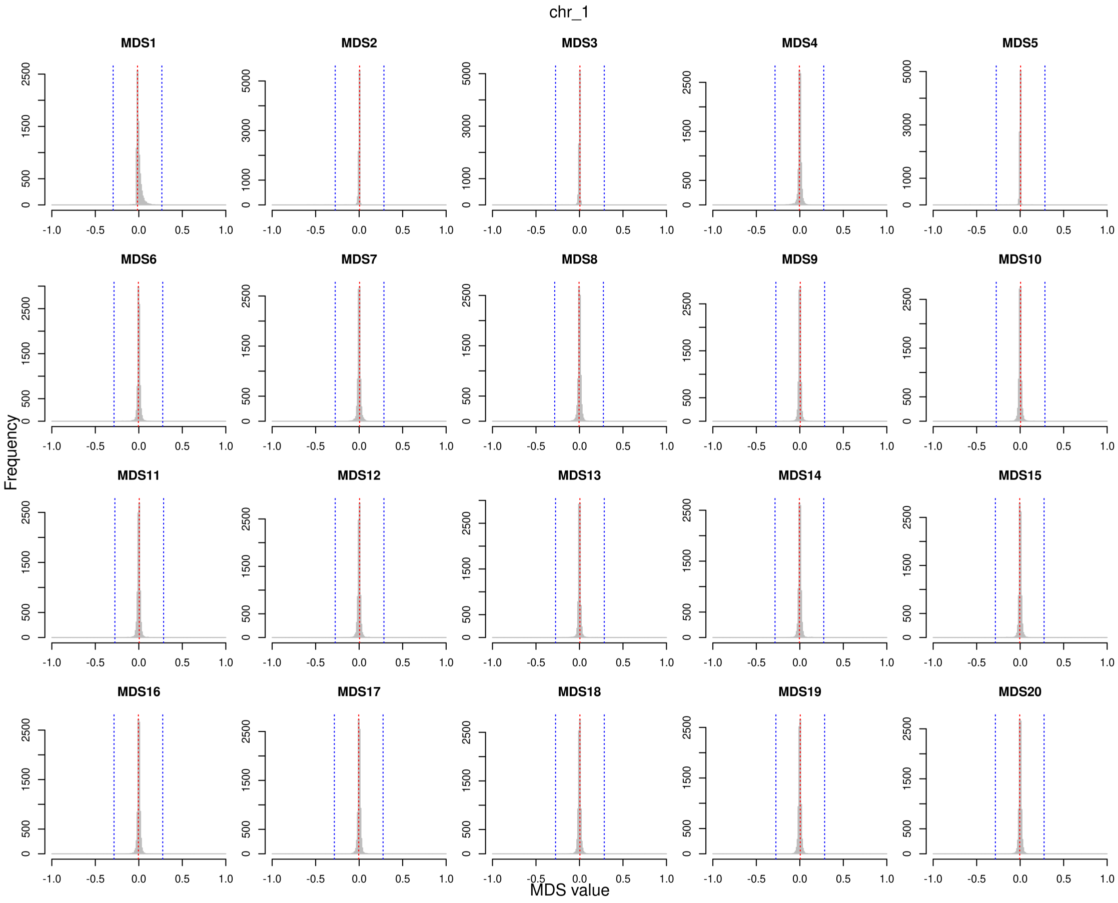

**Supplementary Figure 2: Distribution of MDS values of windows of a chromosome in `lostruct`.** Each panel shows the distribution of MDS values of windows in an exemplified chromosome (chromosoem 1) in local PCA using `lostruct` based on genotypes. Red lines show median of MDS values. Genomic regions with at least 10 windows with MDS values away from the median by 0.3 (blue lines) were defined as "outliers" (see Materials and Methods).

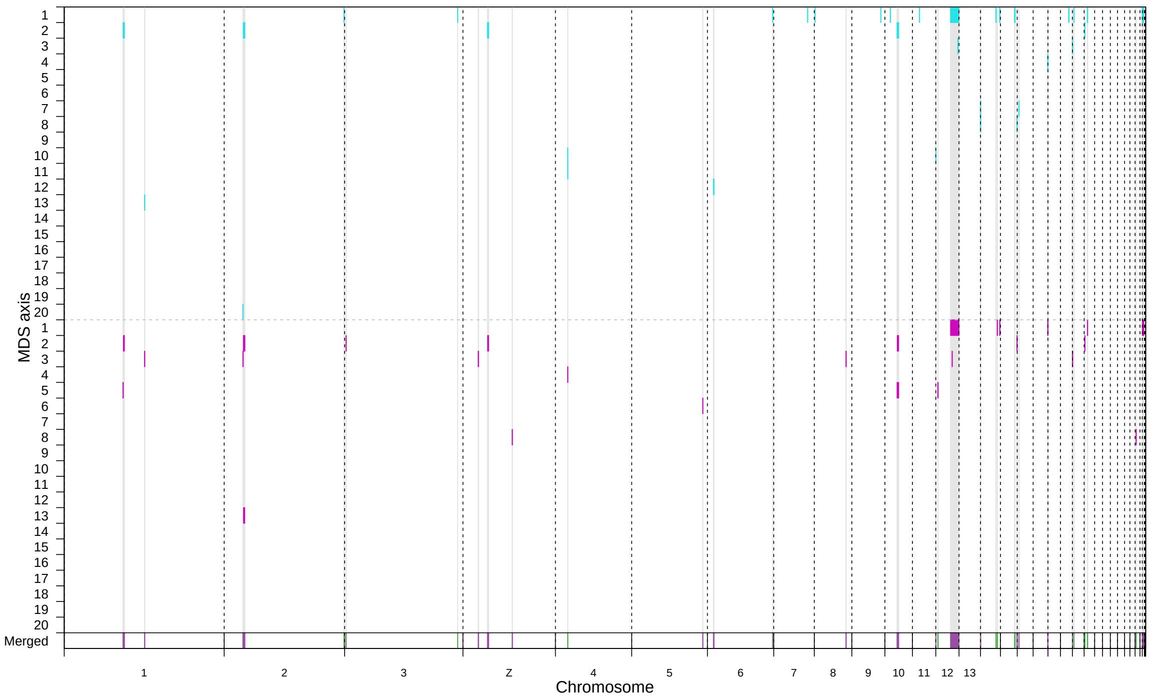

**Supplementary Figure 3: Genomic regions with distinct patterns of genetic variation identified with `lostruct`.** Supplementary content related to Fig. 2D. The y-axis shows the MDS axes along which projected component of genetic variation is deviated in genotype-based (top half, cyan) and haplotype-based (bottom half, purple) analysis of `lostruct`. The x-axis shows the genomic position. The bottom row (colours correnspond to species-wide, population-specific, and no low-recombining in Fig. 2D) shows coordinates of outlier intervals which we determined as final outliers by merging results of the genotype- and haplotype-based analyses.

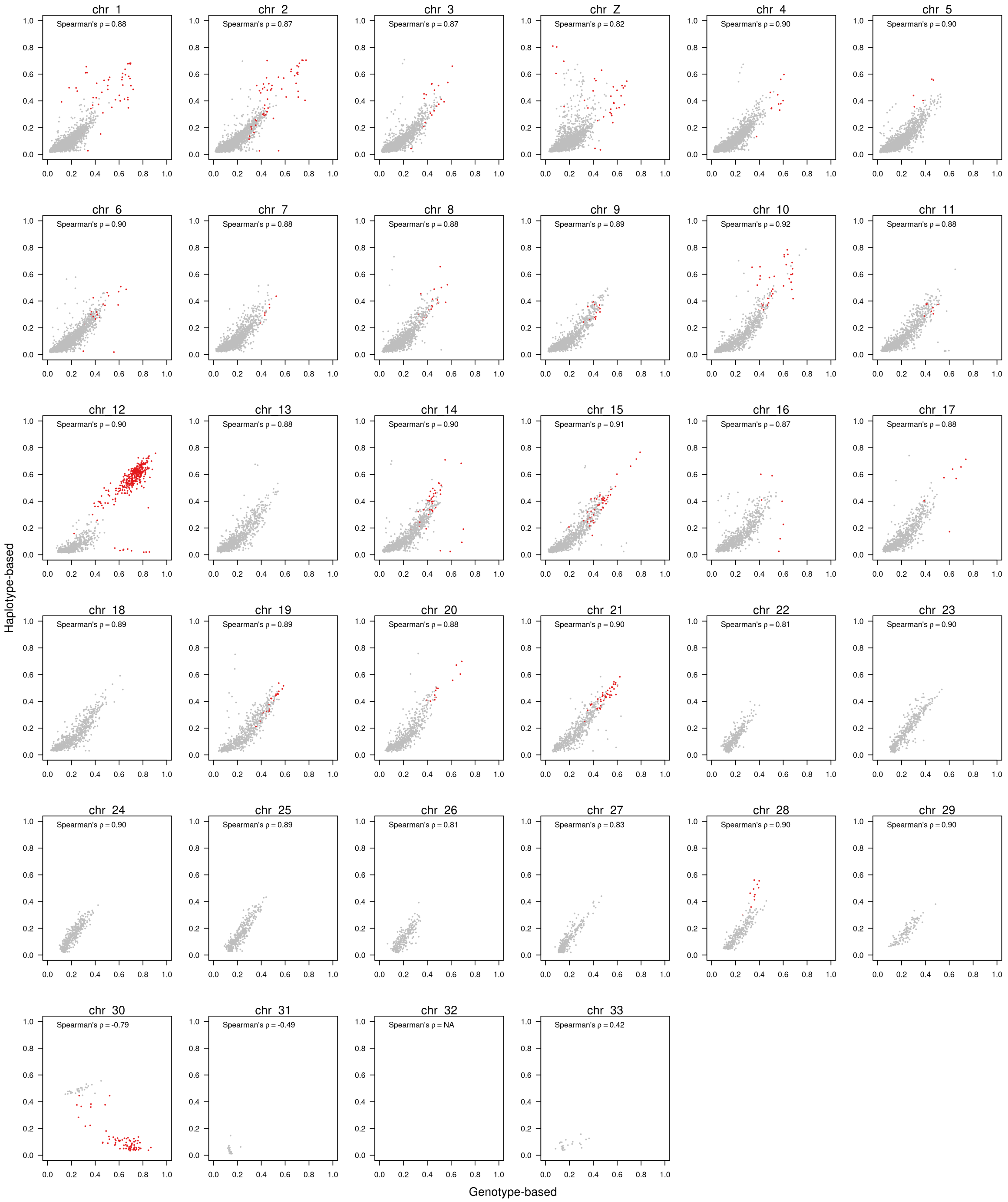

**Supplementary Figure 4: Consistency of genotype- and haplotype-based `lostruct`.** To answer whether genotype- and haplotype-based local PCA using `lostruct` were consistent, Euclidean distance of windows in the 20 dimensional space was compared between these approaches. Red points depict windows with deviated MDS value along a MDS axis in at least one of genotype- or haplotype-based analyses.

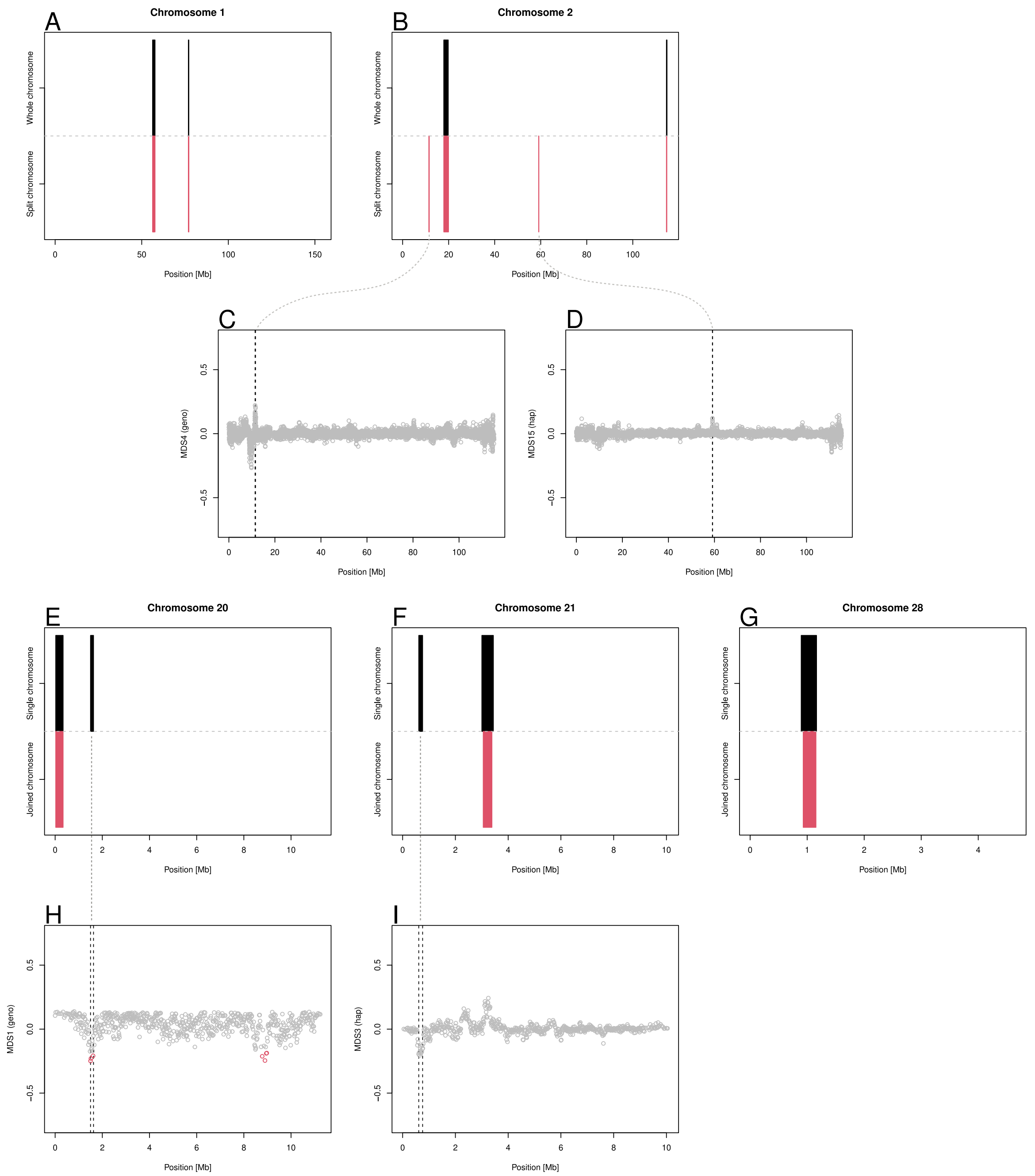

**Supplementary Figure 5: Robustness of `lostruct`.** To address whether `lostruct` is robust to the chromosomal background and the length of the chromosome, we performed `lostruct` using artificially synthesised chromosomes. Blackcap chromosomes 1 and 2 were split into halves in the middle, and chromosomes 20, 21, 28 were joined into a single chromosome. `lostruct` was performed for these split/joined chromosomes and results were compared with single chromosome analysis. **A, B, E-G.** Summary of `lostruct` analysis based on a single chromosome (black) and split/joined chromosome (red). **C, D, H, I.** MDS of single chromosome (**C, D**) and joined chromosome (**H,I**) `lostruct`. Red points depict windows with MDS value beyond the threshold. Per-chromosome analysis was mostly consistent with split/joined chromosome analysis. In regions with inconsistent results, MDS values still show sub-threshold deviation.

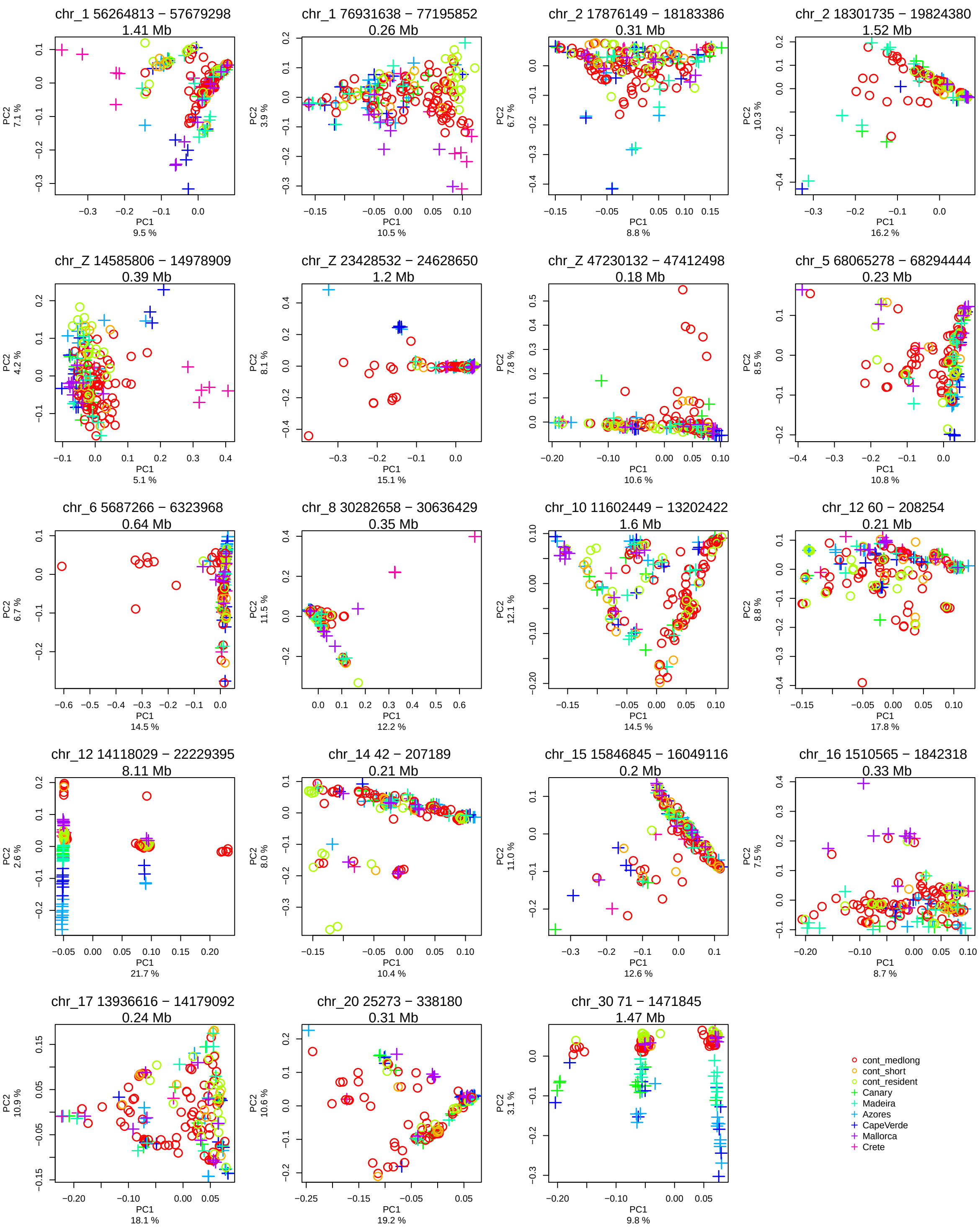

**Supplementary Figure 6: PCA for outliers overlapping species-wide low-recombining regions.** Supplementary data related to Fig. 3. PCA plots represent patterns of genetic variation at 19 outlier regions in the blackcap genome overlapping species-wide low-recombining regions (data points represent blackcap individuals and colours depict populations). The patterns were distinct from population structure (Fig. 2B, C).

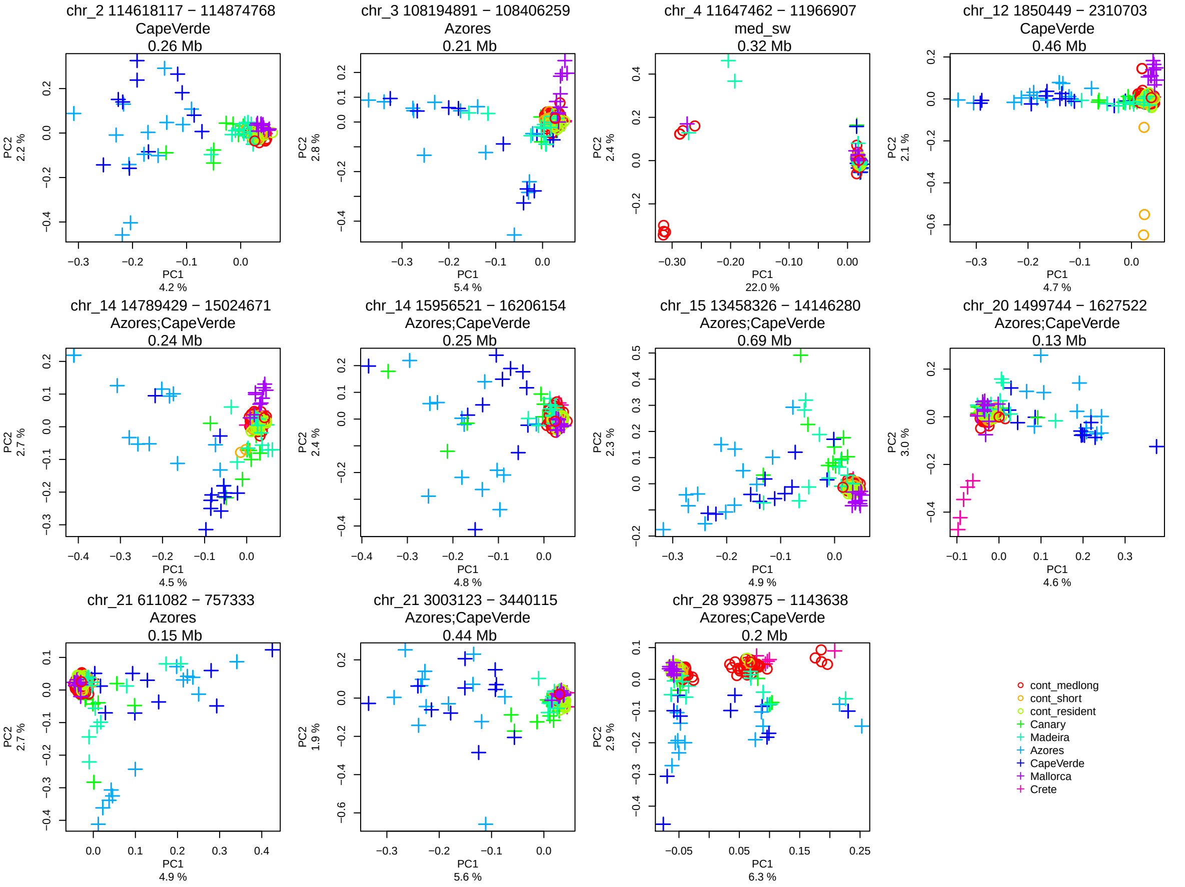

**Supplementary Figure 7: PCA for outliers overlapping only with population-specific low-recombining regions.** Supplementary data related to Fig. 3. PCA plots represent patterns of genetic variation at 11 outlier regions in the blackcap genome overlapping only with population-specific low-recombining regions (data points represent blackcap individuals and colours depict populations). The patterns were distinct from population structure (Fig. 2B, C). Population labels on top of each panel depict the population(s) in which low-recombining regions are found within the outlier.

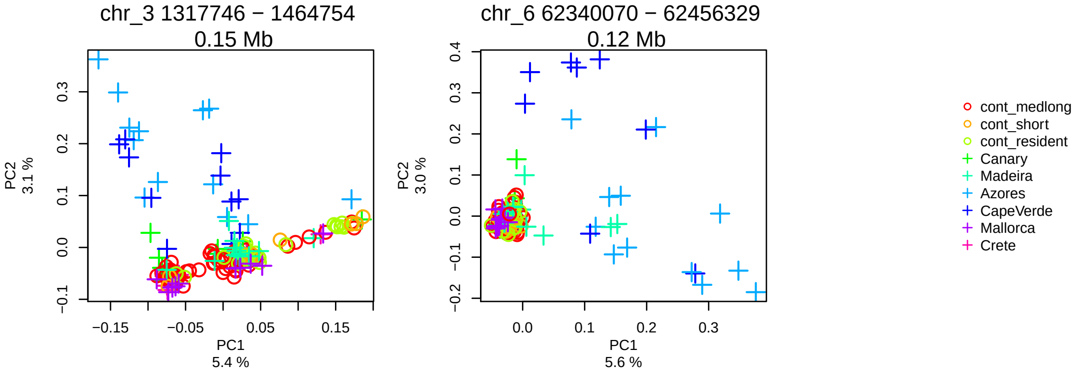

**Supplementary Figure 8: PCA for outlier regions without overlaps with low-recombining regions.** Supplementary data related to Fig. 3. PCA plots represent patterns of genetic variation at two outlier regions in the blackcap genome (data points represent blackcap individuals and colours depict populations).

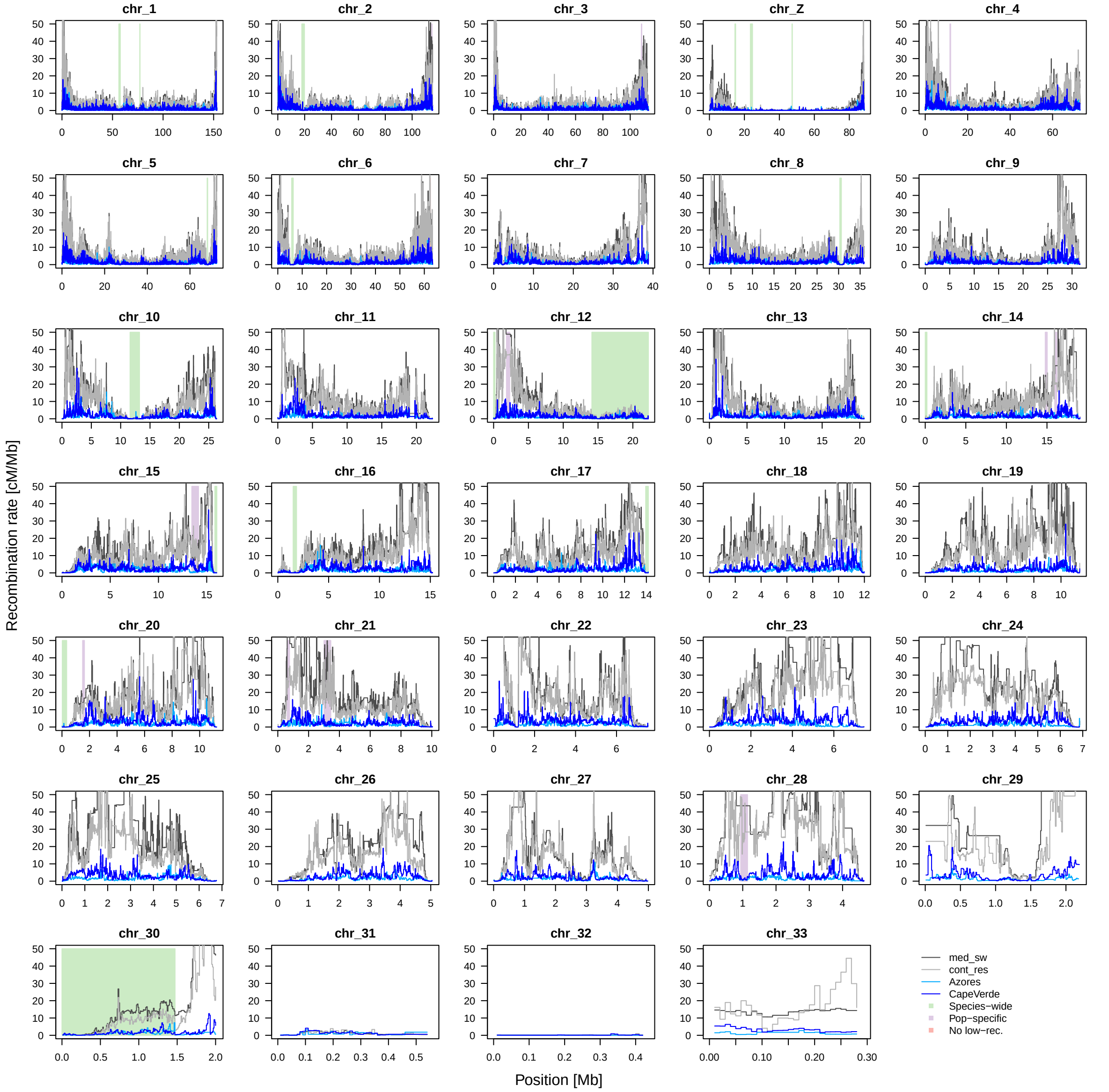

**Supplementary Figure 9: Recombination landscape and `lostruct` outliers.** Supplementary data related to Fig. 2E, F. Four lines depict recombination maps inferred for four blackcap populations. Background shades depict positions of outliers identified by local PCA using `lostruct`.

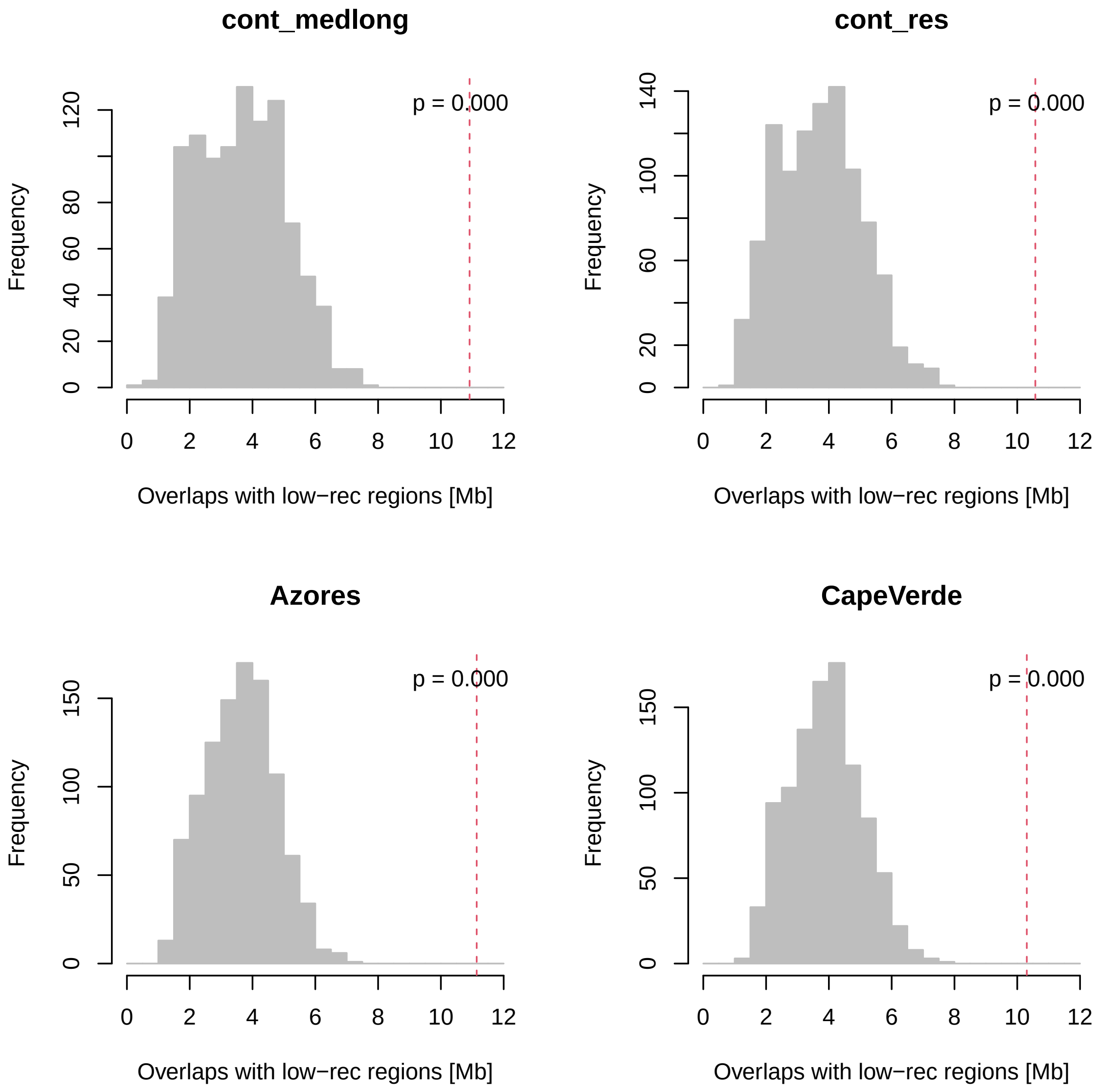

**Supplementary Figure 10: Permutation tests for the number of overlaps between local PCA outliers and low-recombining regions.** To test whether the 32 outlier regions in the blackcap genome based on local PCA significantly overlap with low-recombining regions, permutation tests were performed (n = 1,000). In each permutation, intervals of observed outlier regions were shuffled in each chromosome, and the total length of overlaps with low-recombining regions (below 20 percentile per chromosome) was recorded (see Materials and Methods). Red lines represent observed length of overlaps.

#### 3.3 Putative inversions

##### 3.3.1 PCA

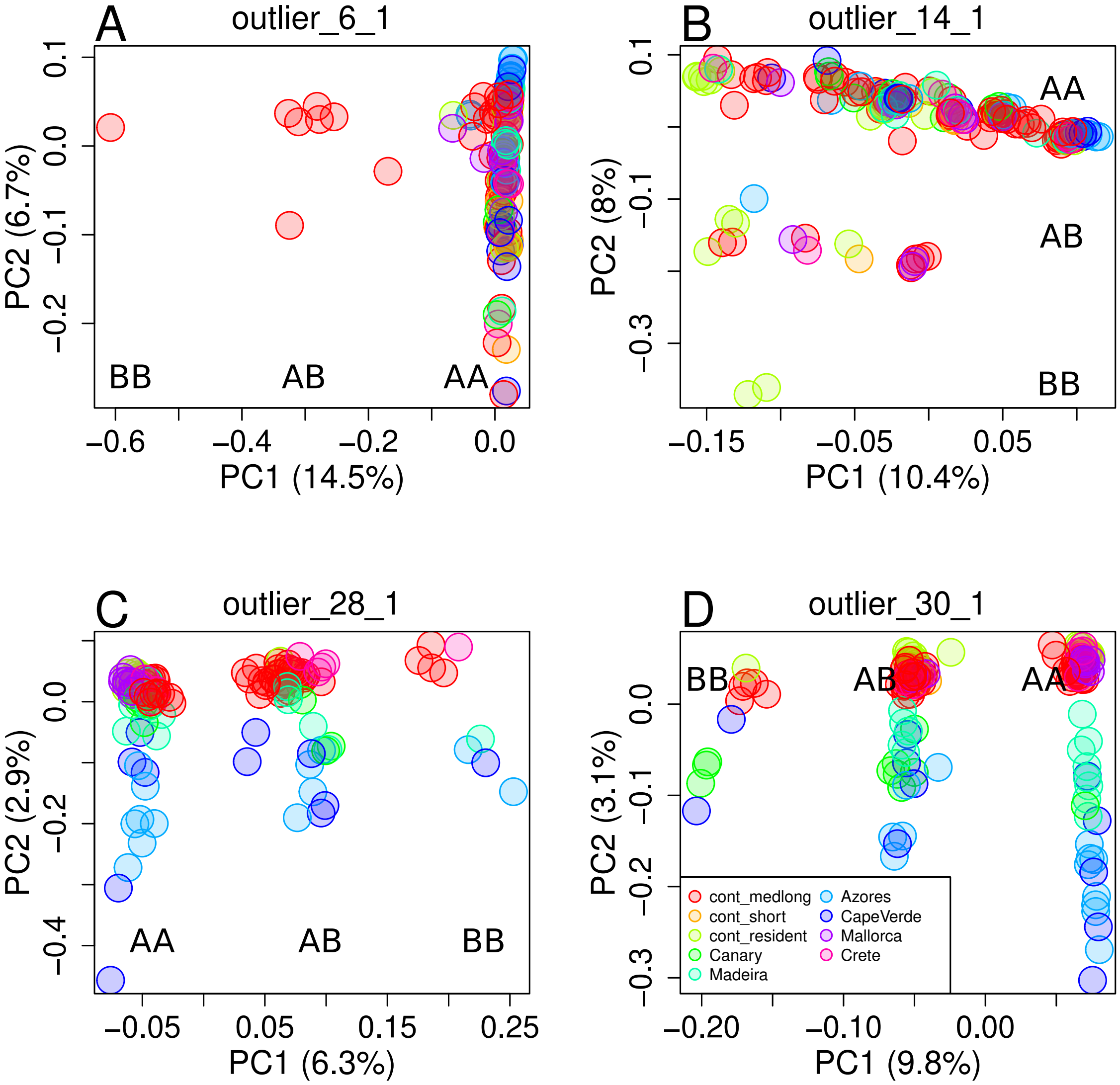

**Supplementary Figure 11: PCA at outliers with three clusters of individuals.** Supplementary data related to Fig. 3A (top). Five outlier regions found in the blackcap genome show patterns of genetic variation in PCA with three clusters of individuals, some of which may represent polymorphic inversions (Huang et al., 2020; Ma & Amos, 2012; Todesco et al., 2020). We investigated this possibility in Sup. Figs. 12, 13, 14. In each region, we named the major and minor alleles A and B, and the three genotypes AA, AB, and BB.

##### 3.3.2 LD and genotype-specific recombination map

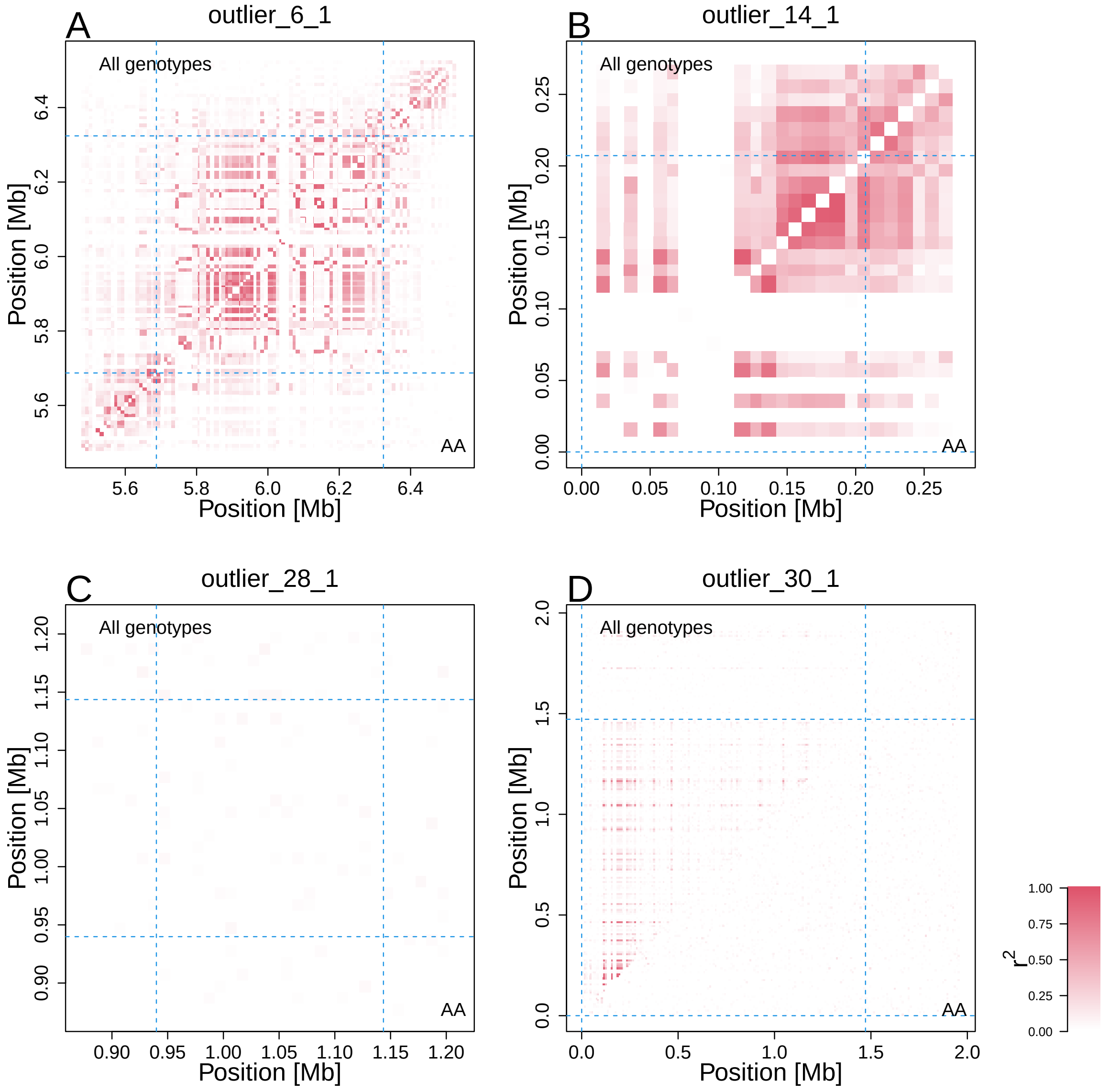

**Supplementary Figure 12: Linkage disequilibrium (LD) at outliers with three clusters in PCA.** Supplementary data related to Fig. 3A (bottom). We investigated whether the five outliers in Sup. Fig. 11 represent five inversions. An inversion locus is expected to show high LD in all individuals but not in homozygous individuals while a non-inversion haplotype block is expected to show high LD irrespective of the genotype. In two outlier regions (outlier_12_3 (Fig. 3A) and outlier_30_1 (**D**)), LD was elevated when all individuals were used, but not when only AA individuals were used, indicating that these two outliers represent putative inversions. The other three did not show this pattern and thus represent non-inversion low-recombining regions. The top-left diagonal shows LD calculated using all blackcap samples. The bottom right diagonal shows LD calculated using AA individuals.

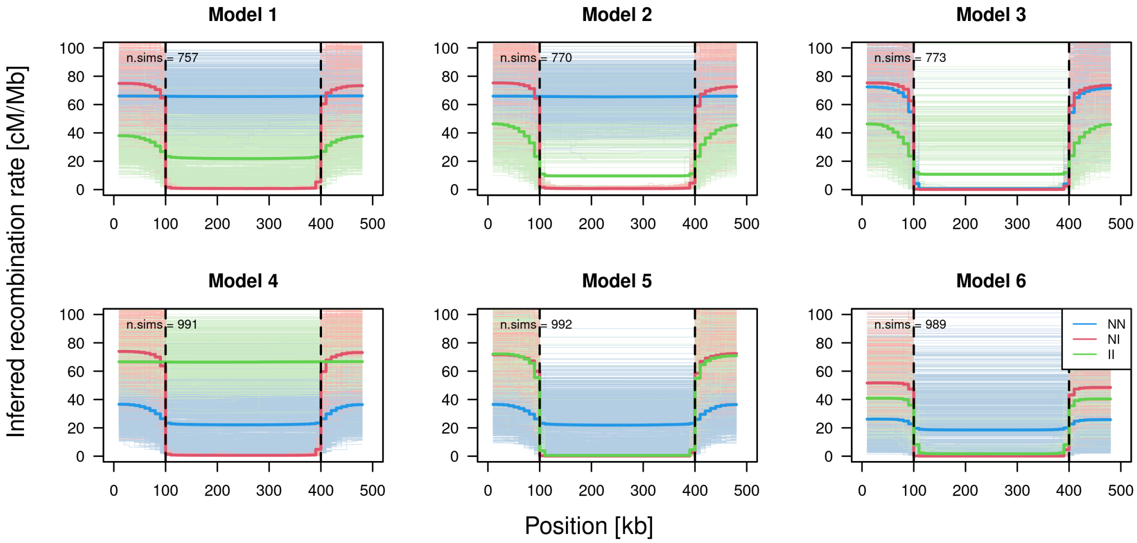

**Supplementary Figure 13: Recombination rate inference by `Pyrho` performed on a chromosome simulated by `SLiM` with different models of recombination suppression.** Supplementary data related to Sup. Fig. 14. We investigated whether inference of genotype-specific recombination maps can capture heterozygote-specific suppression of recombination at polymorphic inversions using simulation before applying this approach to the blackcap data. Six different models of genotype-specific recombination suppression was simulated with `SLiM` (see Materials and Methods and Sup. Table. 6), and recombination map was inferred for each genotype (heterozygotes in red, homozygotes in blue (normal-normal) and green (inversion-inversion)). Thin lines depict inferences for individual simulation replicates, and thick lines depict average of all replicates. The vertical black dotted lines depict the boundary positions of the haplotype block defined by recombination suppression. The result shows that heterozygote-specific suppression can be captured (red lines in models 1 and 4) but normal recombination rates in minor allele homozygotes (green in model 1 and blue in model 4) are not well reflected in inferred maps. Recombination suppression irrespective of genotypes (models 3 and 6) is reflected in all inferred genotype-specific recombination maps.

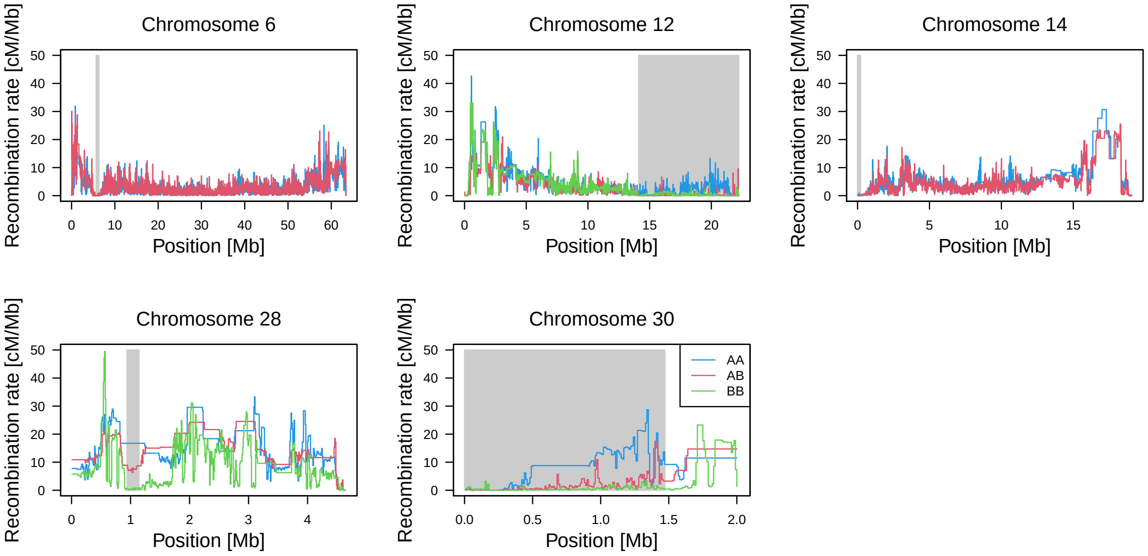

**Supplementary Figure 14: Recombination maps inferred with AA, AB, and BB samples in outlier regions with three clusters of individuals in PCA.** Supplementary data related to Fig. 3A (bottom) and Sup. Fig. 12. Gray shades depict positions of the outlier regions with three clusters of individuals in PCA (Sup. Fig. 11). In line with Sup. Figs. 12, 13, reduced recombination rate in AB with moderate levels of recombination rate in AA was observed at outlier_12_3 and outlier_30_1, indicating these two represent polymorphic inversions. The other threes showed reduced recombination rate both in AA and AB, indicating they represent low-recombining regions without inversions.

##### 3.3.3 Breakpoint analysis

###### 3.3.3.1 Illumina short reads with soft clip

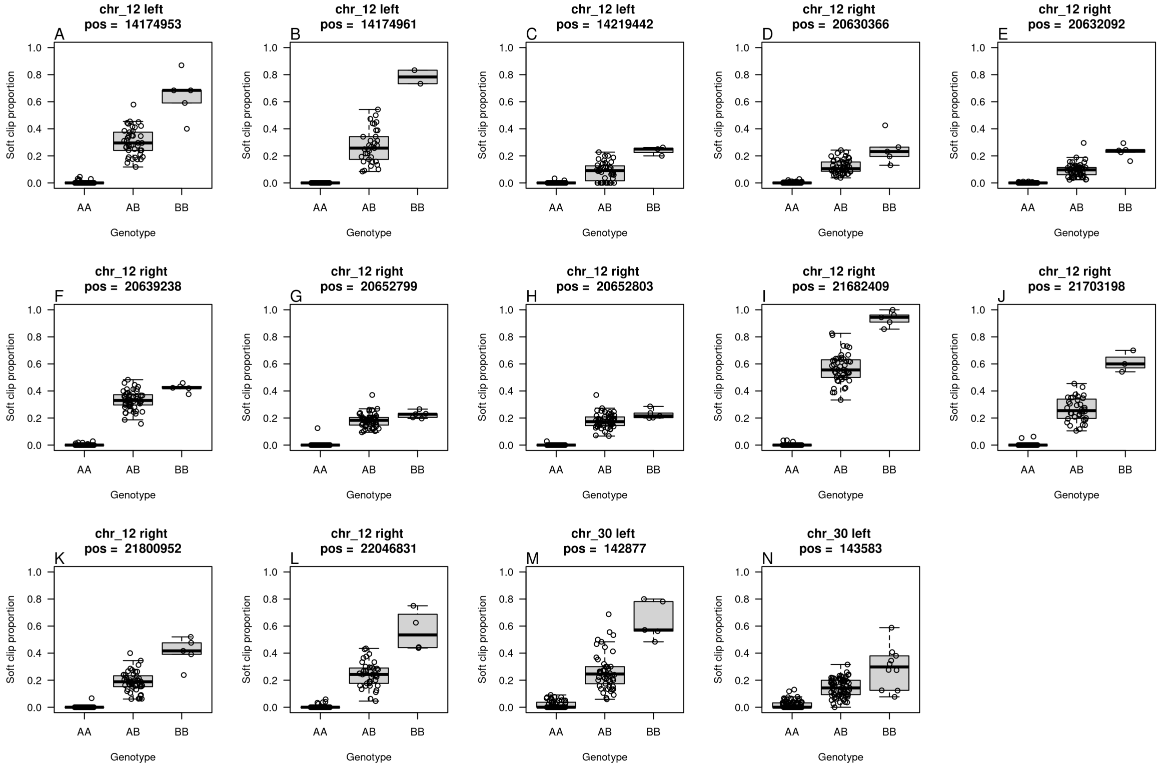

**Supplementary Figure 15: Soft-clip proportion at 14 candidate positions where soft-clip proportion and genotype of putative inversions are associated.** Supplementary data related to Sup. Table. 5. We analysed alignment of Illumina short reads to identify breakpoints of putative inversions at outlier_12_3 and outlier_30_1 (detailed in Materials and Methods). At 14 positions, the proportion of soft-clipped reads was correlated with the genotypes (AA, AB, BB. Detailed in Materials and Methods). The plots show soft-clip proportion of all individuals with three putative inversion genotypes at these 14 positions. Based on these plots, we excluded positions in **C-H** from further analysis in Sup. Table. 5 because the soft-clip proportion of BB was not high or that of AB was not around a half that of BB.

##### 3.3.3.2 10x linked-read sequencing

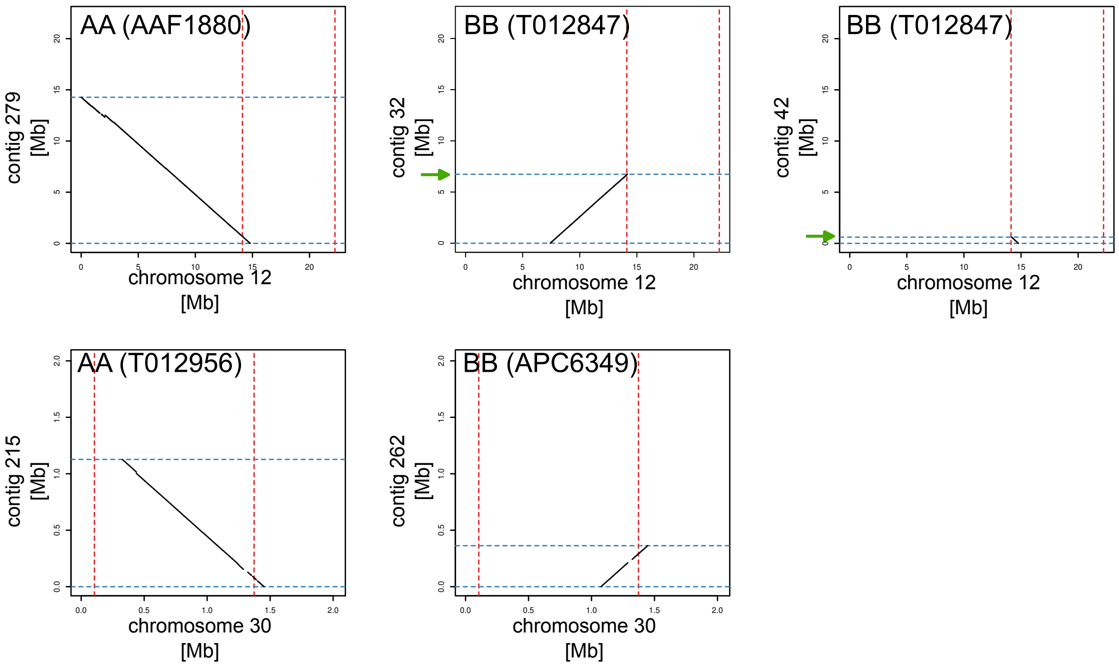

**Supplementary Figure 16: Dot plots between blackcap reference chromosomes 12 or 30 and 10X pseudo-haplotyped contigs of AA and BB individuals.** To identify breakpoints of the putative inversions, we aligned pseudo-haplotyped contigs (based on 10x linked-read sequencing) of AA and BB individuals for outlier_12_3 (top) and outlier_30_1 (bottom) to the blackcap reference genome (Detailed in Materials and Methods). The x-axis shows the coordinate of the blackcap reference chromosomes, and the y-axis shows the coordinate of the 10x pseudo-haplotyped contigs. The red dotted lines depict local PCA-based breakpoint positions in the reference coordinate. The blue dotted lines depict two ends of 10x contigs. In AA individuals (AAF1880 for outlier_12_3 and T012956 for outlier_30_1), one contig was aligned across the putative breakpoint based on local PCA (red dotted lines), supporting the absence of large structural variation. In BB individuals (T012847 for outlier_12_3 and APC6349 for outlier_30_1), no contigs were aligned across the ends of local PCA-based outlier regions, but instead alignment of two 10x contigs terminated (blue dotted lines) at either side of the putative breakpoint, indicating the presence of structural variation in B allele for both outlier_12_3 and outlier_30_1. Presence of large inversions is inconclusive because the ends of these pseudo-haplotyped contigs were not aligned to the other ends of the outlier regions. We investigated why the sequences of B allele is not contiguous across the putative breakpoints in Sup. Fig. 17. Green arrows depict regions in which repeat analyses were performed in Sup. Figs. 17, 18.

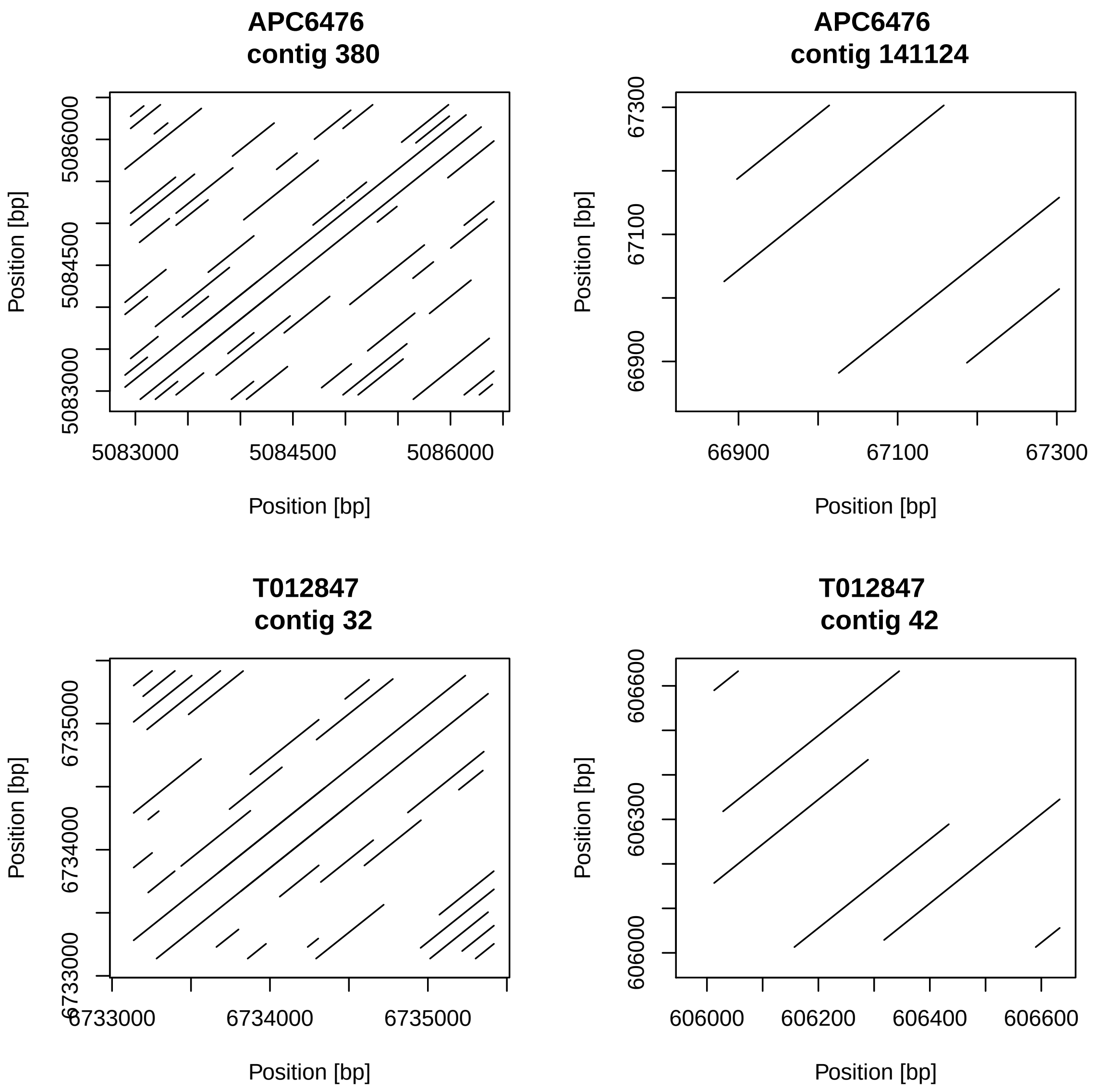

**Supplementary Figure 17: Self dot plots of flanking sequences of 10x linked-read contigs around putative breakpoint of chromosome 12.** Supplementary data related to Sup. Fig. 16. To investigate why 10x contigs of individuals with B allele (presumably inversion) break at the putative breakpoints, we aligned the flanking sequence of the 10x contigs that was aligned next to the putative breakpoint of outlier_12_3 to itself. The plots show the presence of repeats in the sequence flanking the putative breakpoint.

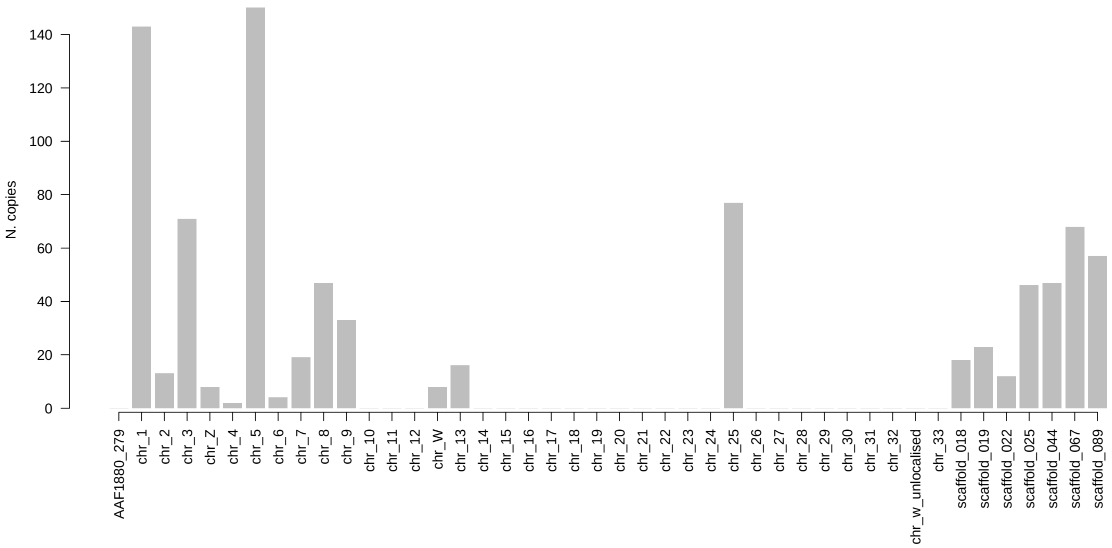

**Supplementary Figure 18: Tandem repeats found at putative inversion breakpoint of chromosome 12 are absent in ancestral allele but present in other chromosomes.** Supplementary data related to Sup. Fig. 17. A 144 bp-long tandem repeat was identified in flanking sequence of the 10x contigs aligned next to the putative inversion breakpoint of outlier_12_3 (Detailed in Materials and Methods). To investigate whether this repeat is present elsewhere in the pseudohaplotype of A allele and the reference, `BLASTn` was performed with the consensus sequence of the tandem repeat unit as the query and contig 279 of AAF1880 (A haplotype contig of 10x) and the blackcap reference as target. The bar plot depicts the number of hits in each chromosome/contig. The result shows the tandem repeat is present in other chromosomes but not on the A allele of chromosome 12.

#### 3.4 Effect of reduced recombination rate on pattern of local genetic variation

##### 3.4.1 Effect of demography and local recombination rate (coalescent simulation)

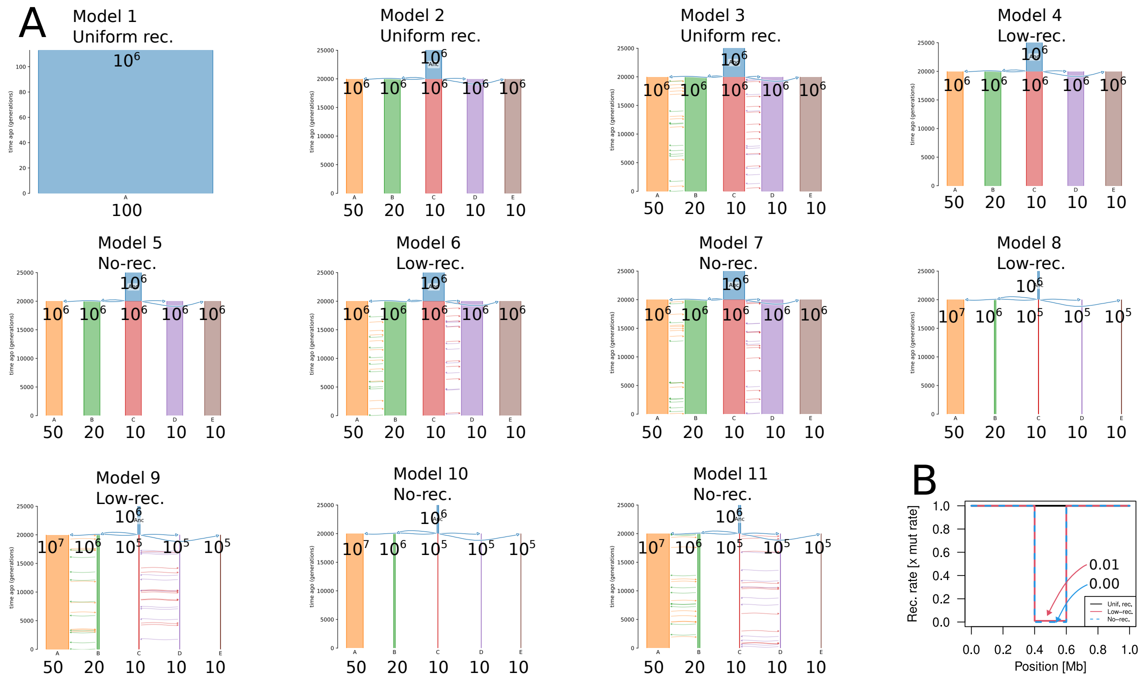

**Supplementary Figure 19: Demography models and recombination maps for neutral coalescent simulations.** Supplementary data related to Sup. Table. 8. To investigate the effects of local recombination rate and demography on genetic variation, we implemented 11 scenarios of demographic history and recombination landscapes ([tab:sup.msp__models]) and simulated SNPs under coalescent with recombination with `msprime` 1,000 times. **A.** 11 models of demographic history. The numbers shown on the populations depict the effective population size, and the numbers below the populations depict the numbers of sampled diploid individuals. Note that demographies for models 2, 4 and 5, models 3, 6 and 7, models 8 and 9, and models 10 and 11 are respectively the same, differing by the recombination map (**B**). **B.** Three recombination maps differing by the recombination rate in the middle of the chromosome. Within the middle interval (0.4 to 0.6 Mb), recombination rate is reduce to 1/100 of the background region in the "low-recombining" scenarios, while in the "no-recombining" scenarios recombination rate is reduced to 0.

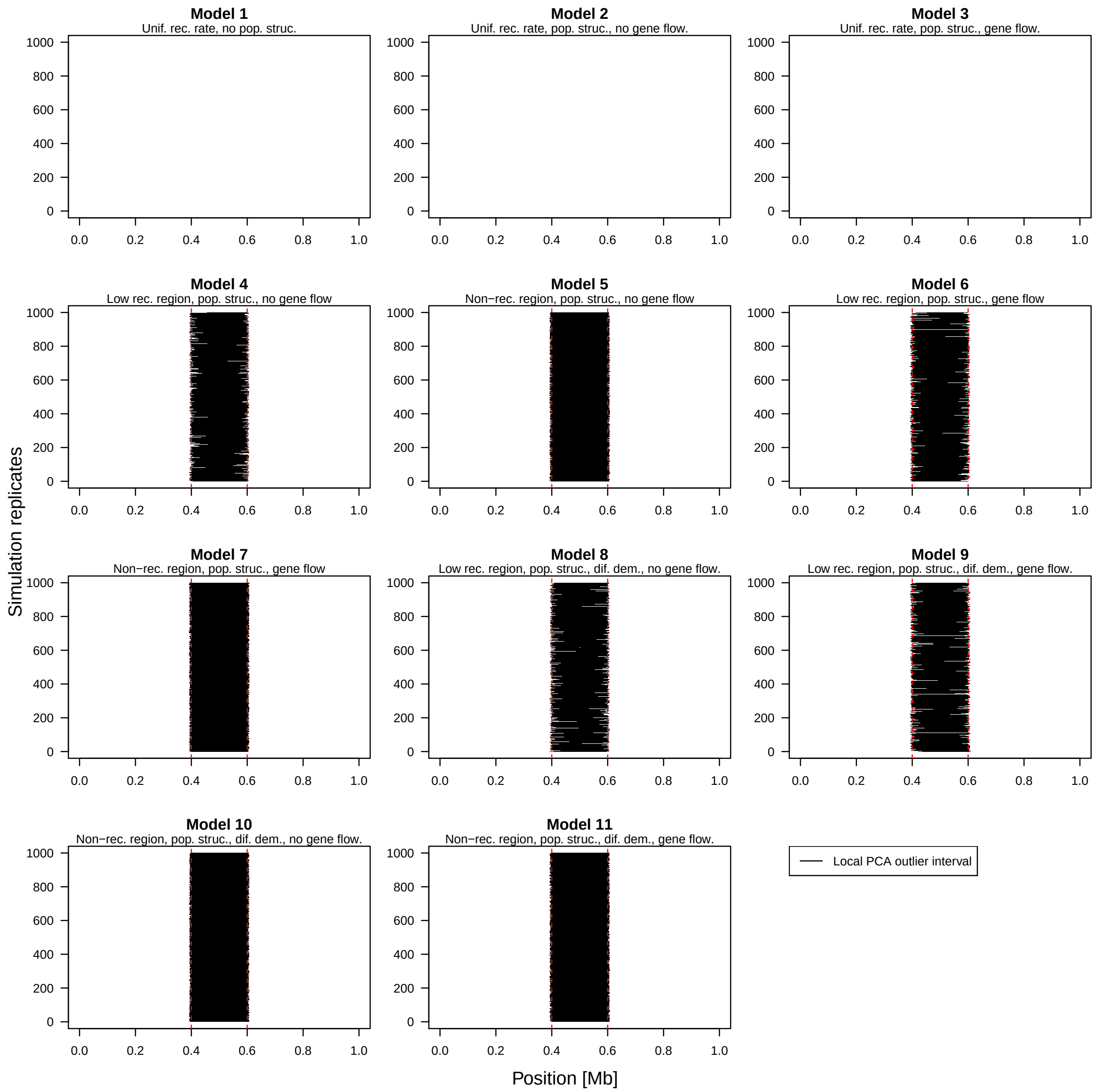

**Supplementary Figure 20: Reduced recombination rate, but not demography, causes distinct patterns of genetic variation.** Supplementary data related to Sup. Fig. 19 and Sup. Table. 8. To identify outlier regions, `lostruct` was performed for each of 1,000 replicates of the 11 scenarios in Sup. Fig. 19. The results show that no outlier regions were detected without reduced recombination rate (models 1-3), and outliers were always detected with reduced recombination rates (models 4-11). This is irrespective of the presence of population structure, unequal demographic history, and unbalanced sample size among populations. These results indicate that reduced recombination rate, but not demography, causes distinct patterns of genetic variation.

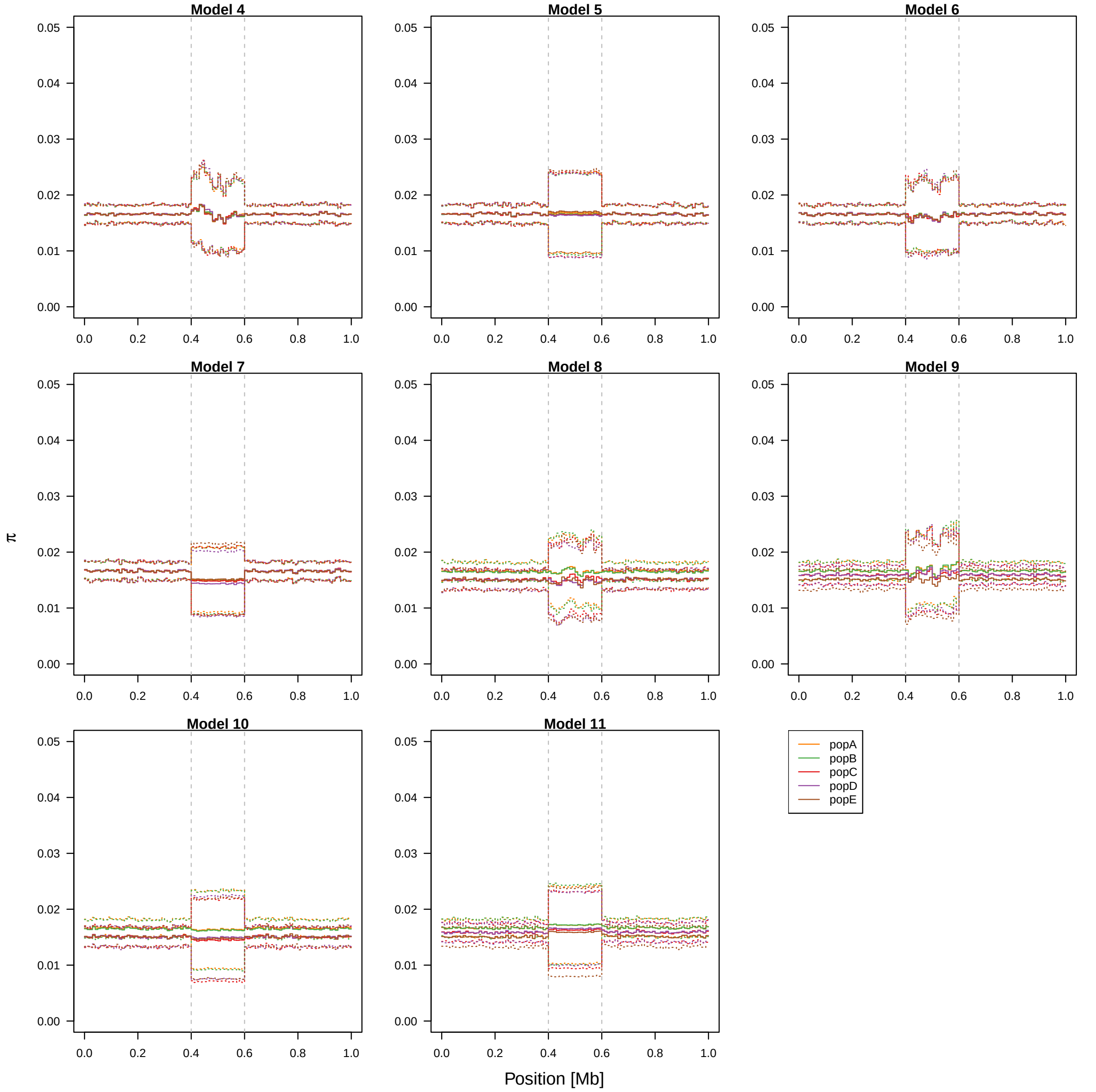

**Supplementary Figure 21: Reduced recombination rate increases variance of summary statistics.** Supplementary data related to Sup. Fig. 19 and Sup. Table. 8. Using the same data as Sup. Fig. 19, we asked how reduced recombination rates affect mean and variance of nucleotide diversity. Solid lines show the meean of the first 100 replicates (out of 1,000 simulated) and dotted lines show the standard deviation.

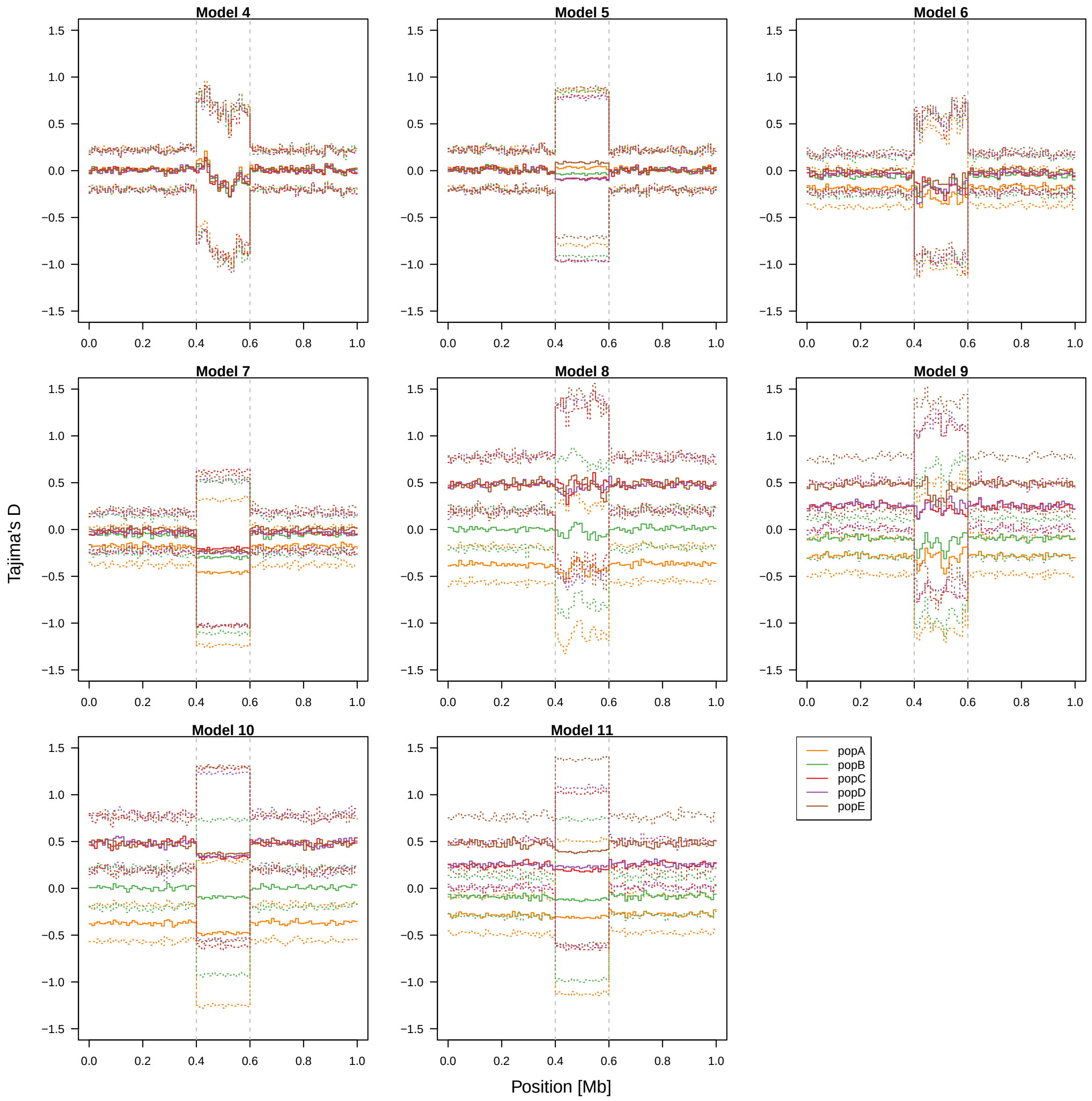

**Supplementary Figure 22: Reduced recombination rate increases variance of summary statistics.** Supplementary data related to Sup. Fig. 19 and Sup. Table. 8. Using the same data as Sup. Fig. 19, we asked how reduced recombination rates affect mean and variance of Tajima's D. Solid lines show the meean of the first 100 replicates (out of 1,000 simulated) and dotted lines show the standard deviation.

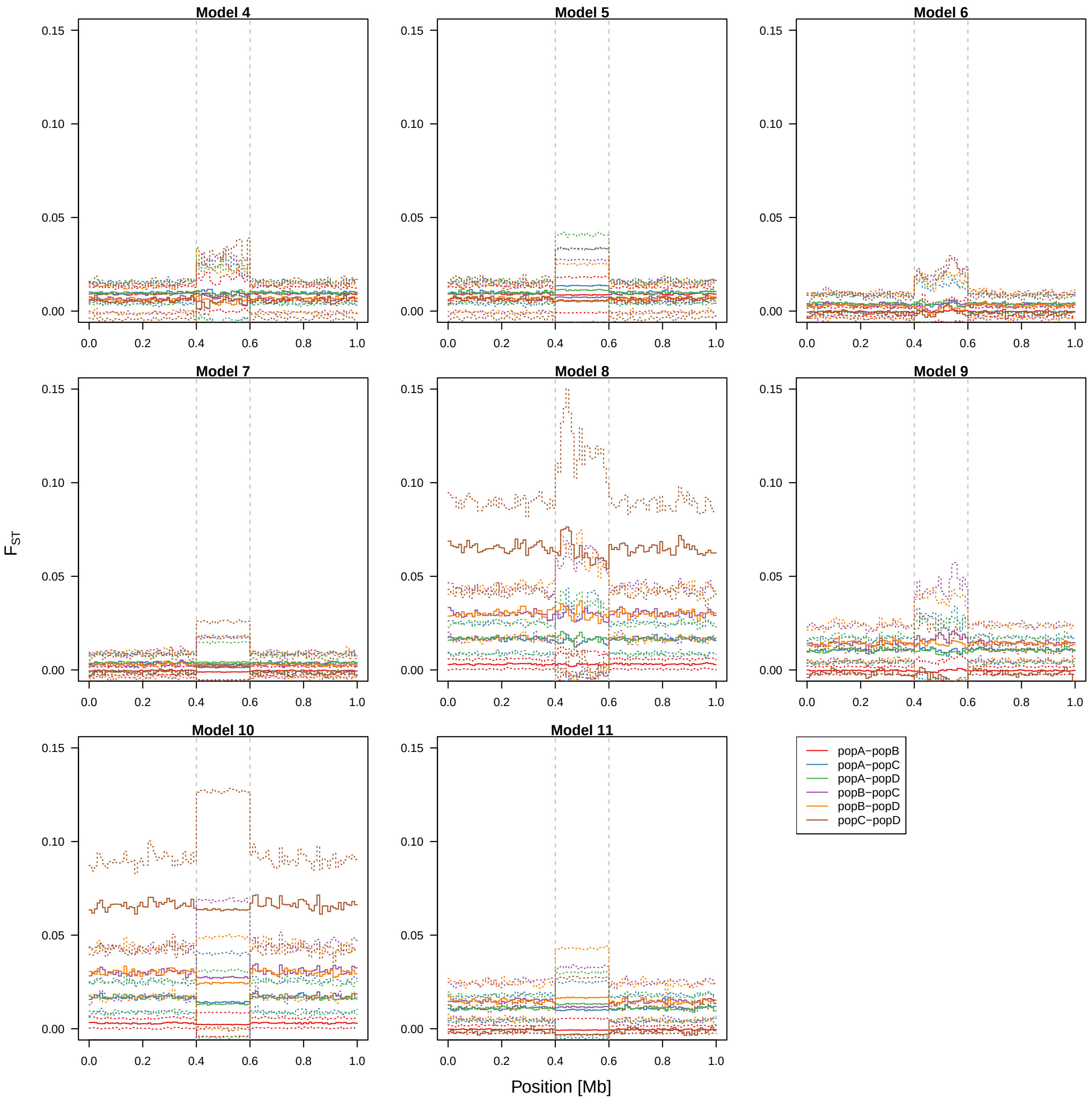

**Supplementary Figure 23: Reduced recombination rate increases variance of summary statistics.** Supplementary data related to Sup. Fig. 19 and Sup. Table. 8. Using the same data as Sup. Fig. 19, we asked how reduced recombination rates affect mean and variance of $F_{ST}$ between five population pairs (of all 10 pairs). Solid lines show the meean of the first 100 replicates (out of 1,000 simulated) and dotted lines show the standard deviation.

##### 3.4.2 Species-wide reduction of local recombination rate (forward simulation)

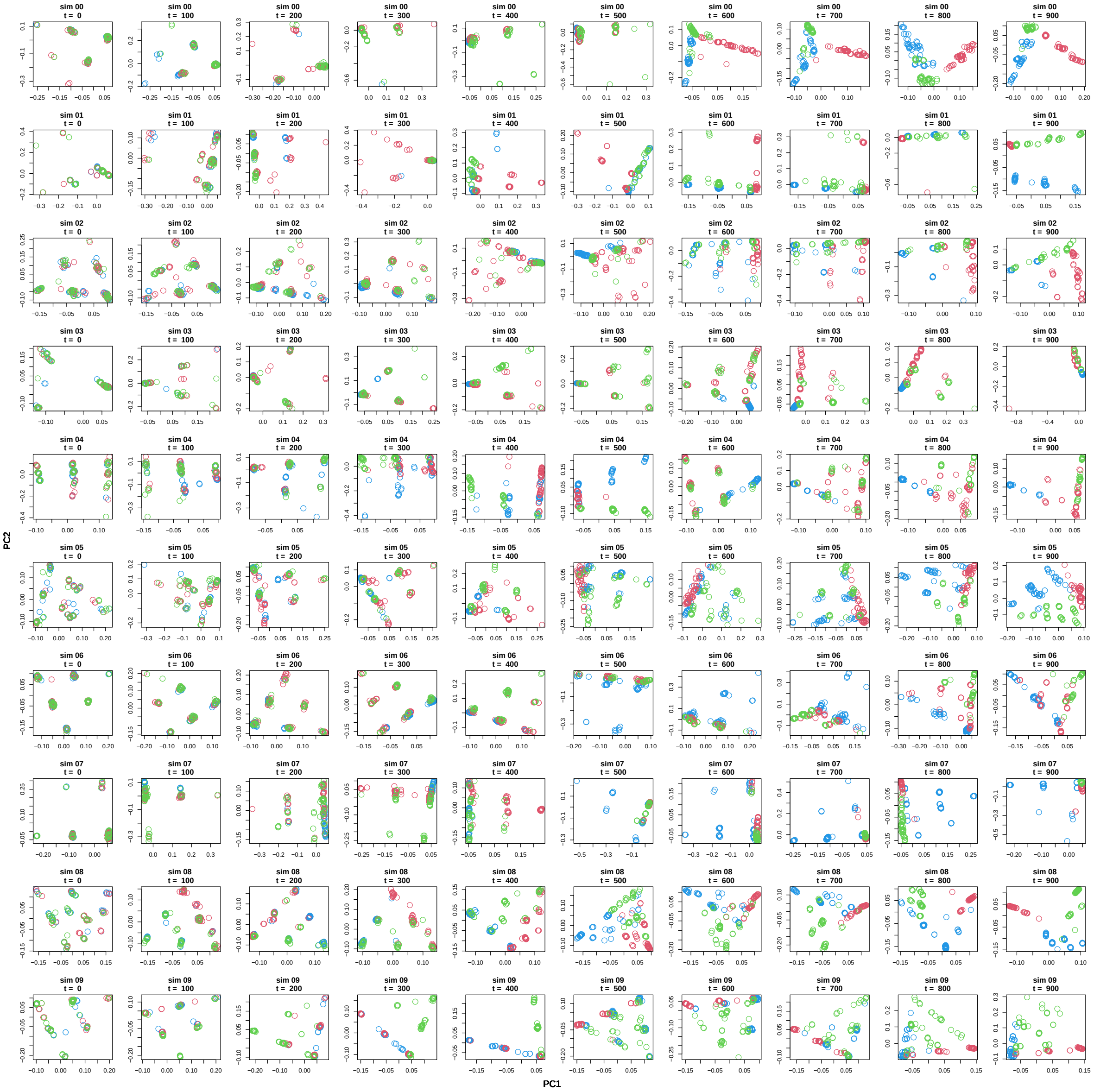

**Supplementary Figure 24: PCA at a species-wide low-recombining region at ten time points of ten exemplified simulation replicates.** Supplementary data related to Fig. 4. To investigate the effects of species-wide reduction in local recombination rate, we simulated one ancestral population of 1,000 diploids with a low-recombining genomic region that splits into three subpopulations (pop1, pop2, pop3. Fig. 4A). All mutations were neutral. We sampled individuals over time after the population split and conducted PCA in the low-recombining genomic region. 10 rows represent 10 simulation replicates (out of 100, see Materials and Methods). 10 columns represent 10 time points. Data points with three different clours depict individuals from three different populations. In addition to Fig. 4B and C, these exemplified results show high variability in realised genetic variation at low-recombining regions across replicates with three to six clusters with different degrees of mixture of individuals in PCA and transitioning from haplotype structure to population structure over time.

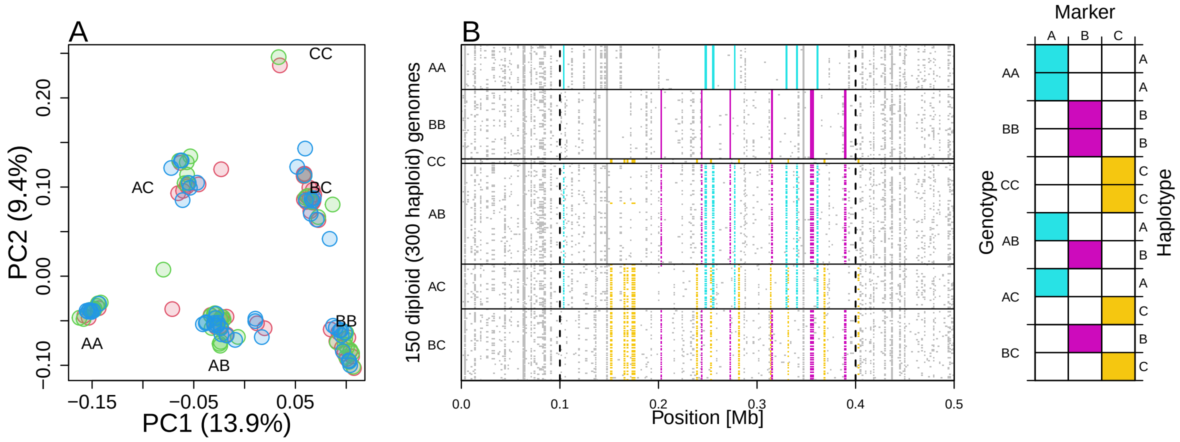

**Supplementary Figure 25: Clusters of individuals in local PCA represent haplotype structure.** Supplementary data related to Figs. 3, 4. **A.** Local PCA of a species-wide low-recombining region simulated showing six clusters of individuals (the same as Fig. 4C, t=0). **B.** Genotypes at thinned mutation sites. Cyan, magenta, and yellow correspond to A-, B-, and C-specific mutations. The distribution of these mutations in homozygous (AA, BB, CC) and heterozygous(AB, AC, BC) individuals suggests that the clusters of individuals in PCA represent combination of haplotypes possessed by diploid individuals.

###### 3.4.3 Population-specific reduction of local recombination rate (forward simulation)

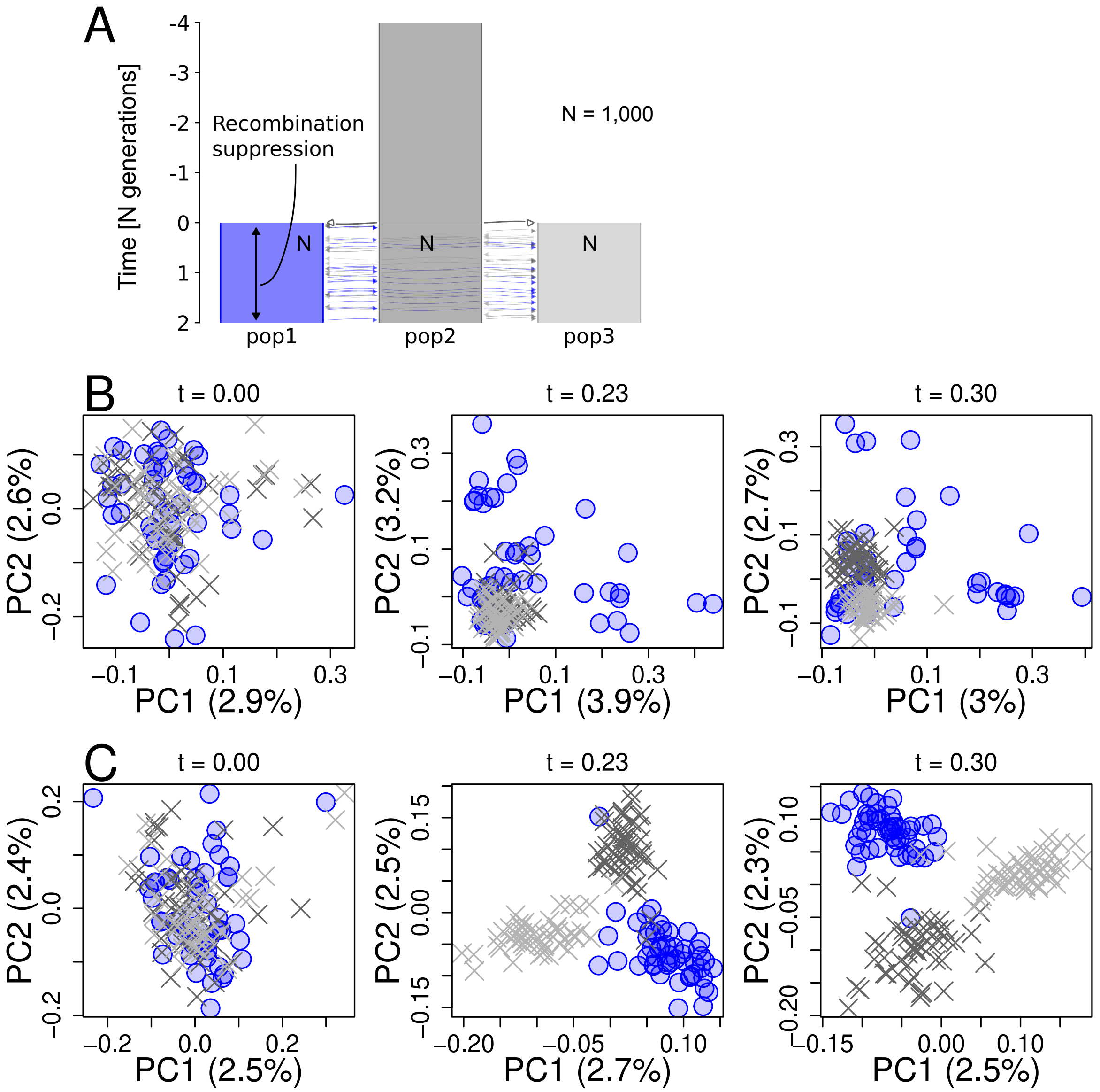

**Supplementary Figure 26: Summary of model 1 of population-specific recombination suppression.** Supplementary data related to Fig. 5. **A.** Simulated scenario. Simulated genome contained two chromosomes, one with a population-specific low-recombining region and the other without. **B, C.** PCA showing patterns of genetic variation at the population-specific low-recombining region (**B**) and the normally recombining chromosome (**C**) at three time points in one exemplified simulation replicate.

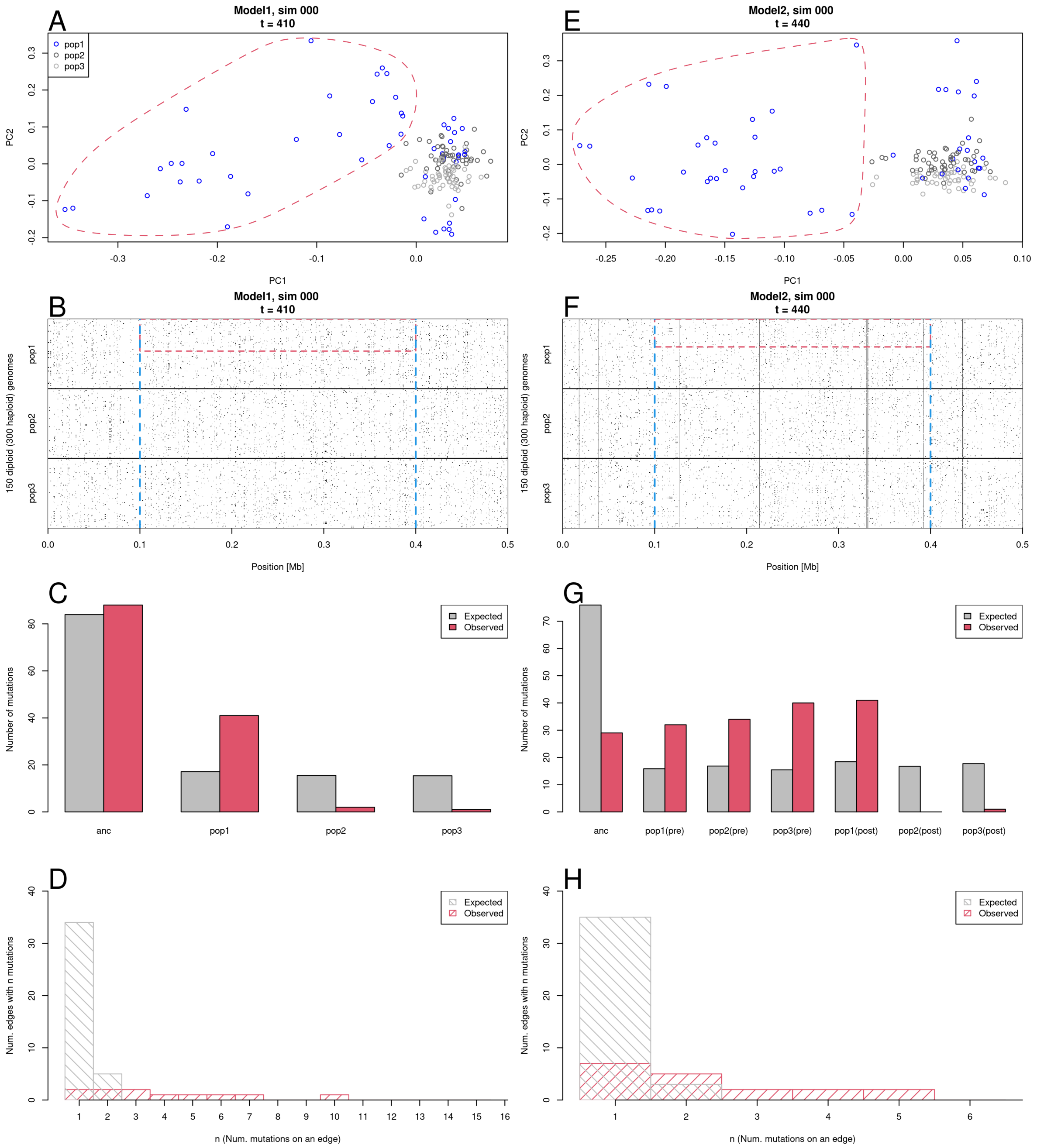

**Supplementary Figure 27: Distinct local PCAs at population-specific low-recombining region represent cryptic haplotype structure.** To characterise factors represented in the primary axes of distinct local PCA at population-specific low-recombining regions, we performed one replicate of SLiM simulation with the scenarios of models 1 (**A-D**) and model 2 (**E-H**) recording the full ancestry and mutations in tree sequence in addition to VCF files (Detailed in Materials and Methods). We performed PCA, and identified mutations with the highest contributions to the PC1 and PC2. We analysed the tree sequence to address whether mutations that occurred in certain population (e.g. ancestral population, low-recombining population) were enriched in the set of mutations contributing to the PC1 and PC2. **A, E.** Local PCA at population-specific low-recombining region. **B, F.** No visible haplotype structure were found at population-specific low-recombining region. **C, G.** Mutations originating from the low-recombining population after the population-specific recombination suppression were enriched in mutations with high loading to the distinct pattern of local PCA. **D, H.** Mutations originating from the low-recombining population with high PCA loading share common genealogical edges. Due to recombination suppression, the mutations on the same edge are on the same haplotype in the current sample.

#### 3.5 Effect of selection

##### 3.5.1 Effects of selection in blackcap genome

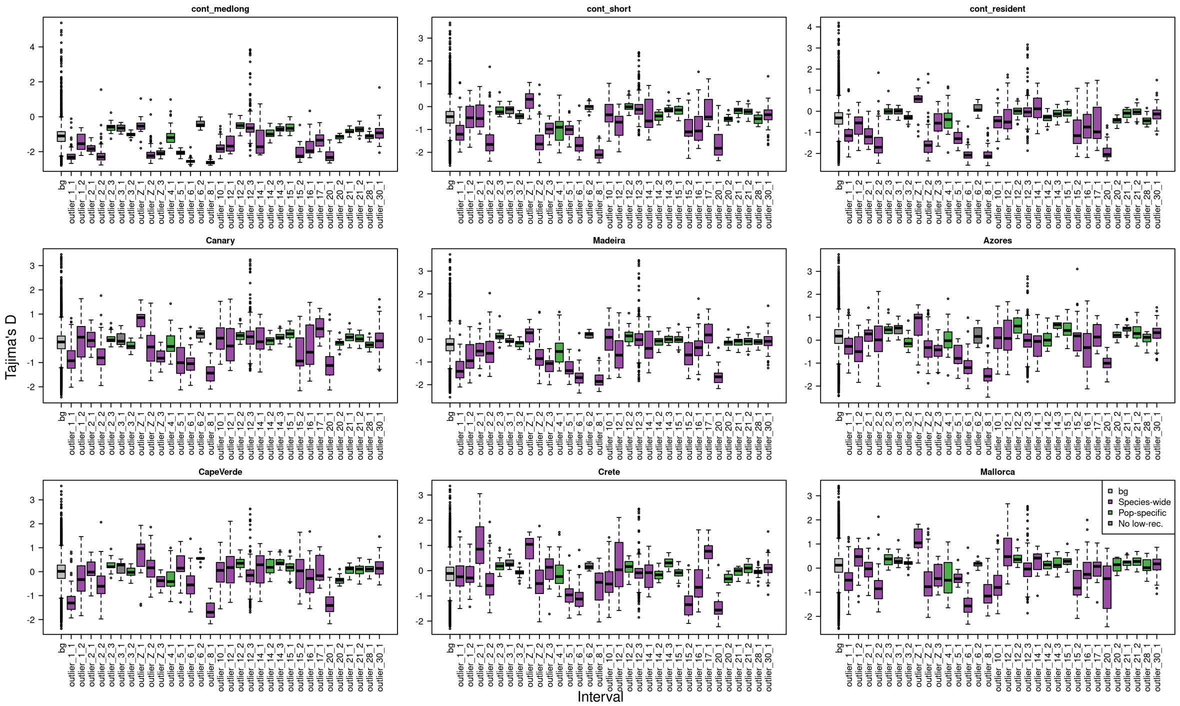

**Supplementary Figure 28: Tajima's D in genomic regions with distinct pattern of genetic variation.** Supplementary data related to *Results: Effect of selection on patterns of genetic variation* and Sup. Table. 9. Box plots depict the distribution of Tajima's D in windows within 32 outlier regions and in non-outlier regions ("bg").

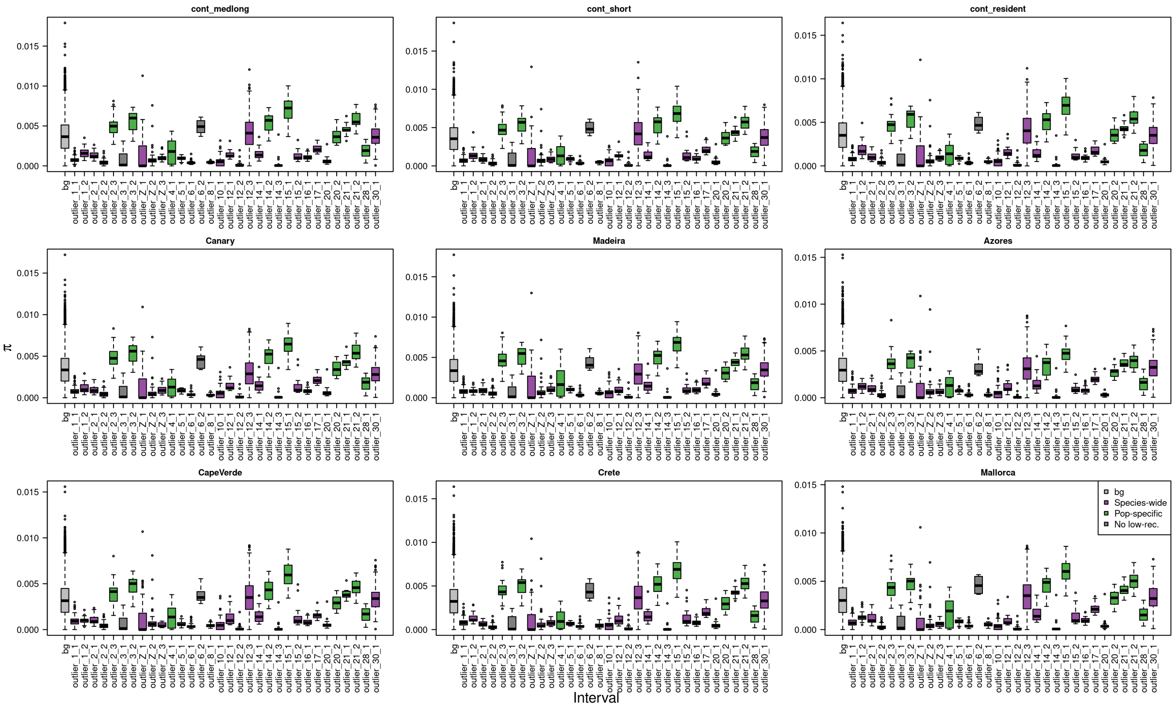

**Supplementary Figure 29: Nucleotide diversity ($\pi$) in genomic regions with distinct pattern of genetic variation.** Supplementary data related to *Results: Effect of selection on patterns of genetic variation* and Sup. Table. 10. "bg" depicts distribution of $\pi$ in non-outlier regions of local PCA.

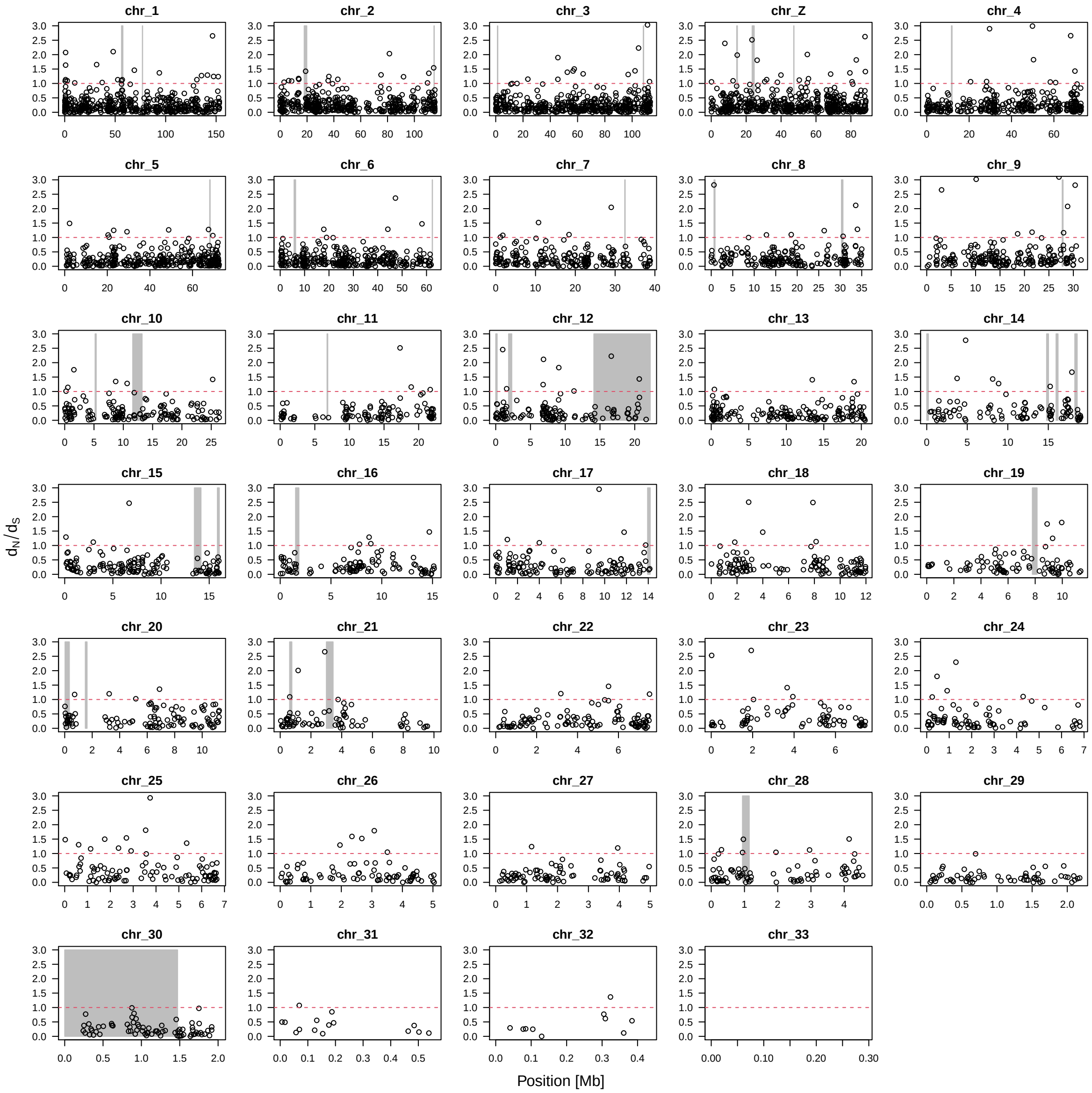

**Supplementary Figure 30:** $d_N/d_S$ **of blackcap genes.** Supplementary data related to *Results: Effect of selection on patterns of genetic variation.* Gray shades depict positions of 32 outlier regions.

##### 3.5.2 Effects of selection at low-recombining regions (forward simulation)

###### 3.5.2.1 Purifying selection

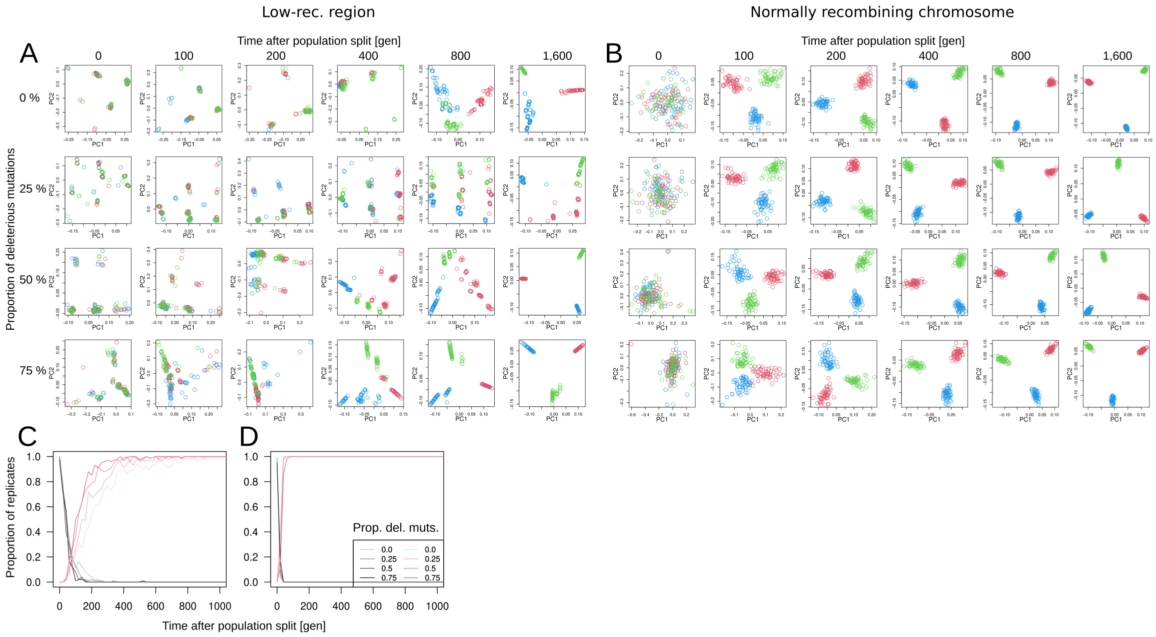

**Supplementary Figure 31: Species-wide reduction in recombination rate and purifying selection affect rate at which local genetic variation changes.** To investigate the effect of purifying selection on patterns of genetic variation at low-recombining regions, we simulated three populations with a species-wide low-recombining region, varying the distribution of fitness effect (DFE) of mutations by specifying three different ratios between neutral and deleterious mutations (Detailed in Materials and Methods). We performed PCA within and outside the low-recombining region over time, and tested whether the distribution of individuals in PCA is distinct between populations by applying Fasano-Franceschini tests. **A.** PCA at low-recombining region over time in one exemplified replicate under the four scenarios of DFE. Points with three different colours depict individuals of three different populations. **B.** PCA with the normally recombining chromosome over time in the same replicate in **A** under the four scenarios of DFE. Clusters of individuals representing haplotype structure is present at all scenarios, but population structure emerges at different rates across DFEs. **C, D.** Summary of Fasano-Franceschini tests in low-recombining (**B**) and normally recombining (**C**) regions. Red lines show the proportion of simulation replicates in which Fasano-Franceschini tests were significant in all three pairs of populations. Black/gray lines show the proportion of simulation replicates in which none of the three pairs of populations were significant in Fasano-Franceschini test. These results show that (1) species-wide recombination reduction slows down change in pattern of local genetic variation (differentiation at a genomic local scale) after a population split event, and (2) purifying selection accelerates the slowed change in pattern of local genetic variation in a chromosomal region with species-wide recombination reduction.

###### 3.5.2.2 Positive selection

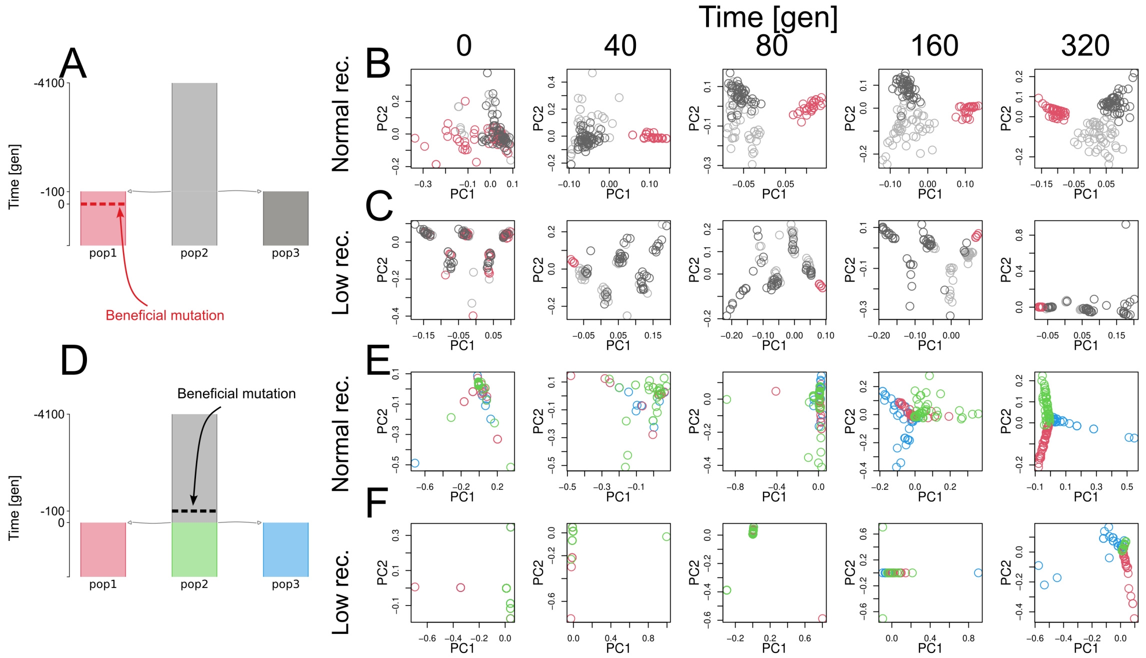

**Supplementary Figure 32: Positive selection at species-wide low-recombining region.** To investigate how positive selection affects patterns of genetic variation at low-recombining regions, we simulated three populations with or without a species-wide low-recombining region and introduced a beneficial mutation before or after population split. We performed PCA over time. **A.** Scenario of population-specific positive selection. A beneficial mutation was introduced in pop1 100 generations after the population split. **B.** PCA around the position of beneficial mutation in the scenario of **A** without a low-recombining region. **C.** PCA around the position of beneficial mutation within a low-recombining region in the scenario of **A**. **D.** Scenario of positive selection in an ancestral population. A beneficial mutation was introduced in the ancestral population 100 generations before the population split. **E.** PCA around the position of beneficial mutation in the scenario of **D** without a low-recombining region. **F.** PCA around the position of beneficial mutation within a low-recombining region in the scenario of **D**. These results show that positive selection affects local genetic variation which is overlaid on the effect of haplotype structure by reduced recombination rate. Only five time points of one simulation replicate are shown in this figure.

#### 3.6 Pericentromeric regions in outlier regions

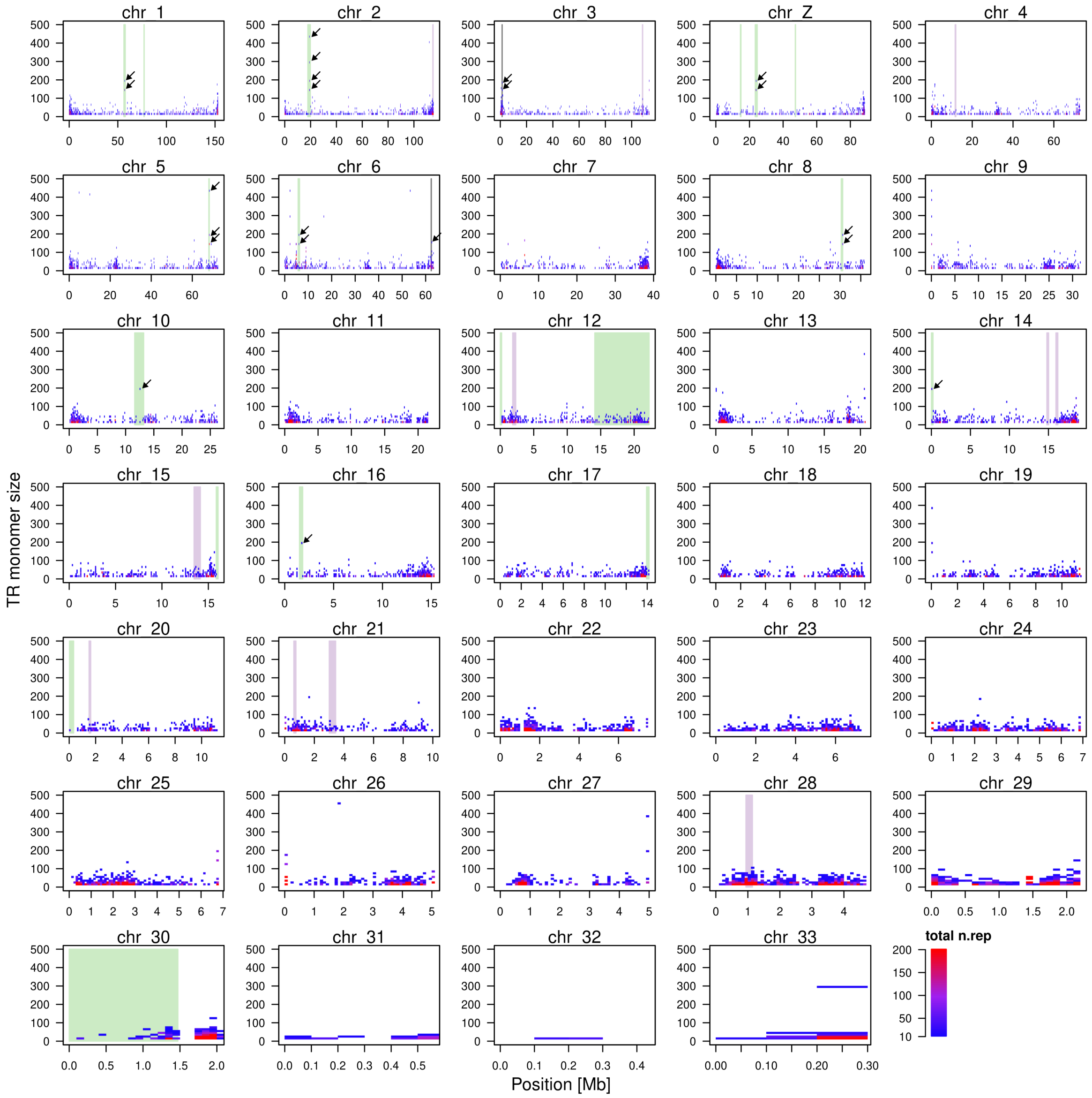

**Supplementary Figure 33: Genomic distribution of tandem repeats.** Shades indicate outlier regions based on `lostruct`. Each cell in the heatmap shows the number of copies of tandem repeats (colour-coded) with unit size of the focal range (y-axis) found in the focal genomic window of 100 kb (x-axis). Black arrows point long tandem repeats coinciding with `lostruct` outlier regions.

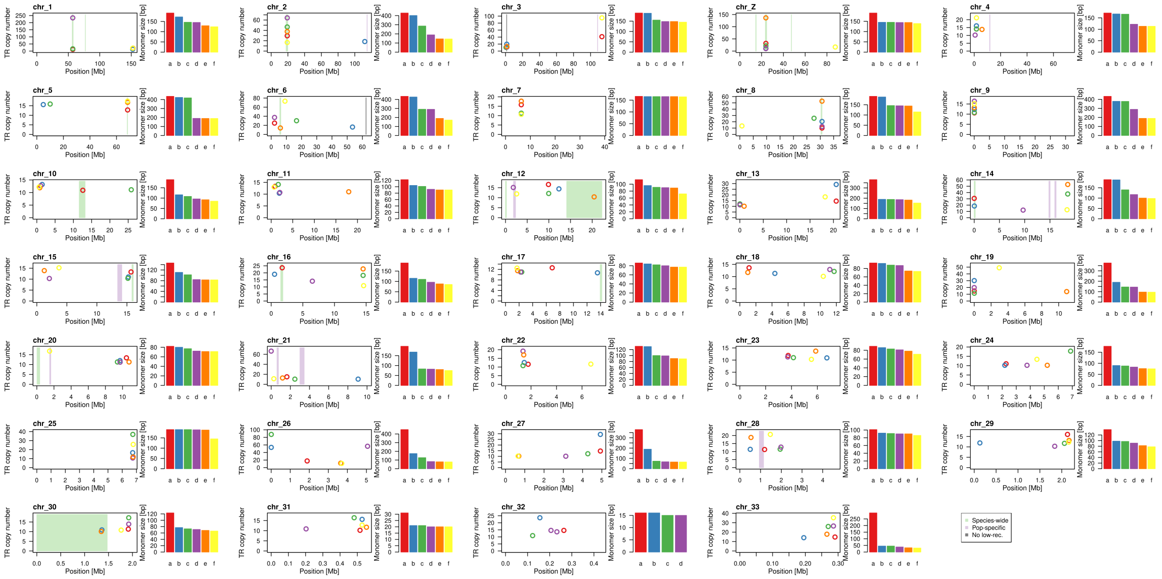

**Supplementary Figure 34: Genomic distribution of tandem repeats with long repeat unit.** In each chromosome, six tandem repeats with the longest repeat units are shown. Shades indicate genomic regions with distinct patterns of genetic variation.

##### 3.7 Genealogical interpretation

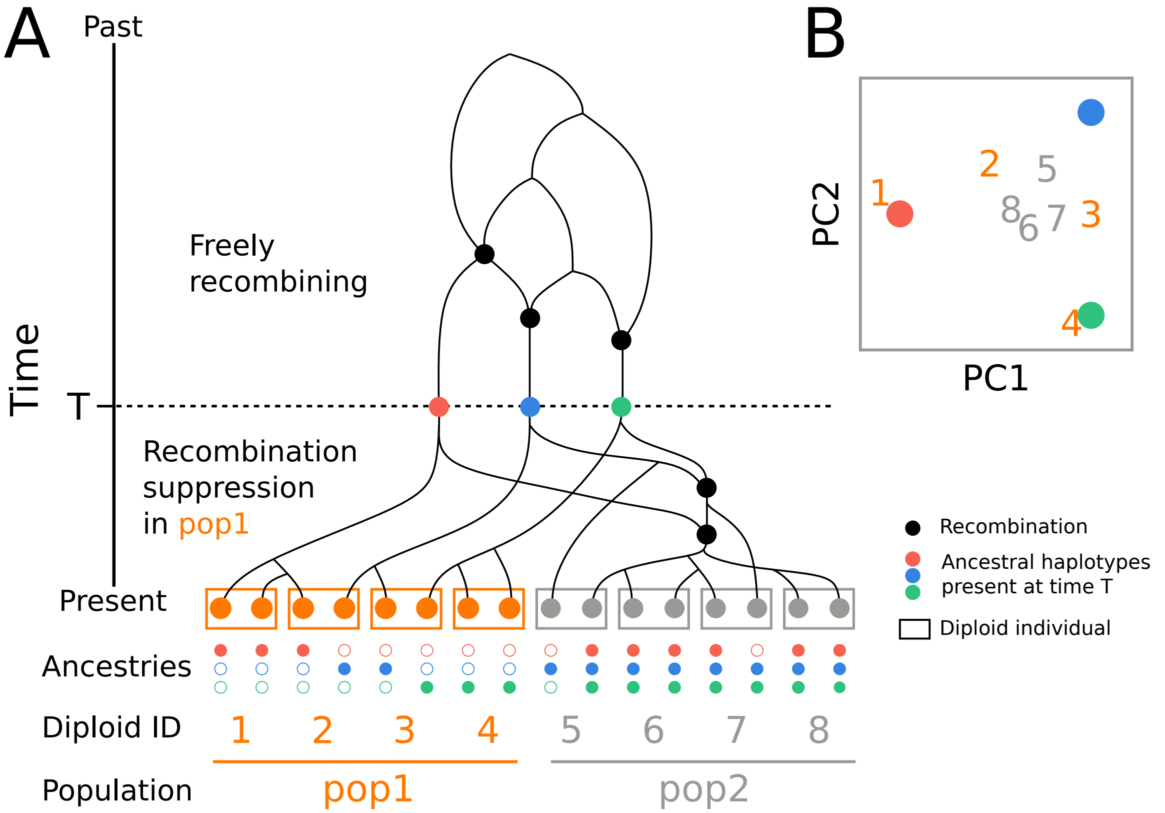

**Supplementary Figure 35: Genealogical interpretation of the effect of population-specific recombination suppression on local genetic variation. A.** An ancestral recombination graph (ARG) representing ancestries of 16 hypothetical haploid sequences of 8 diploid sampled individuals from two populations. Their ancestries can be traced back to $n_1 = n_2 = 3$ ancestral haplotypes (the same set for simplicity) present at time $T$ when population-specific recombination suppression initiated in pop1. The ancestries of these $n_1$ and $n_2$ ancestral haplotypes freely recombine at times older than $T$. At the bottom, the ancestries of each current haplotype are shown. Closed and open circles represent presence and absence of contribution from the respective ancestral haplotype. **B.** A hypothetical PCA representing the pattern of genetic variation of the focal region. Individuals from the low-recombining population (pop1) are spread in the hypothetical PCA because each diploid has a combination of discrete ancestries. The individuals of normally recombining populations (pop2) are clustered around the centre because they have mixed haplotypes due to continued recombination after $T$.
